# An open resource of structural variation for medical and population genetics

**DOI:** 10.1101/578674

**Authors:** Ryan L. Collins, Harrison Brand, Konrad J. Karczewski, Xuefang Zhao, Jessica Alföldi, Laurent C. Francioli, Amit V. Khera, Chelsea Lowther, Laura D. Gauthier, Harold Wang, Nicholas A. Watts, Matthew Solomonson, Anne O’Donnell-Luria, Alexander Baumann, Ruchi Munshi, Mark Walker, Christopher Whelan, Yongqing Huang, Ted Brookings, Ted Sharpe, Matthew R. Stone, Elise Valkanas, Jack Fu, Grace Tiao, Kristen M. Laricchia, Valentin Ruano-Rubio, Christine Stevens, Namrata Gupta, Lauren Margolin, Genome Aggregation Database Production Team, Genome Aggregation Database Consortium, Kent D. Taylor, Henry J. Lin, Stephen S. Rich, Wendy Post, Yii-Der Ida Chen, Jerome I. Rotter, Chad Nusbaum, Anthony Philippakis, Eric Lander, Stacey Gabriel, Benjamin M. Neale, Sekar Kathiresan, Mark J. Daly, Eric Banks, Daniel G. MacArthur, Michael E. Talkowski

## Abstract

Structural variants (SVs) rearrange large segments of the genome and can have profound consequences for evolution and human diseases. As national biobanks, disease association studies, and clinical genetic testing grow increasingly reliant on genome sequencing, population references such as the Genome Aggregation Database (gnomAD) have become integral for interpreting genetic variation. To date, no large-scale reference maps of SVs exist from high-coverage sequencing comparable to those available for point mutations in protein-coding genes. Here, we constructed a reference atlas of SVs across 14,891 genomes from diverse global populations (54% non-European) as a component of gnomAD. We discovered a rich landscape of 433,371 distinct SVs, including 5,295 multi-breakpoint complex SVs across 11 mutational subclasses, and examples of localized chromosome shattering, as in chromothripsis. The average individual harbored 7,439 SVs, which accounted for 25-29% of all rare protein-truncating events per genome. We found strong correlations between constraint against damaging point mutations and rare SVs that both disrupt and duplicate protein-coding sequence, suggesting intolerance to reciprocal dosage alterations for a subset of tightly regulated genes. We also uncovered modest selection against noncoding SVs in *cis*-regulatory elements, although selection against protein-truncating SVs was stronger than any effect on noncoding SVs. Finally, we benchmarked carrier rates for medically relevant SVs, finding very large (≥1Mb) rare SVs in 3.8% of genomes (~1:26 individuals) and clinically reportable incidental SVs in 0.18% of genomes (~1:556 individuals). These data have been integrated directly into the gnomAD browser (https://gnomad.broadinstitute.org) and will have broad utility for population genetics, disease association, and diagnostic screening.

## INTRODUCTION

Structural variants (SVs) are genomic rearrangements that alter segments of DNA larger than 50 nucleotides.^1^ By virtue of their size and abundance, SVs represent an important mutational force shaping genome evolution and function,^2,3^ and contribute to germline and somatic diseases.^4–6^ The profound impact of SVs is partially attributable to the varied mechanisms by which these intra- and inter-chromosomal rearrangements alter linear and three-dimensional genome structure, which can disrupt protein-coding genes or *cis*-regulatory architecture.^7–9^ SVs can be grouped into distinct mutational classes, including “unbalanced” SVs associated with gains or losses of DNA (*e.g.*, copy-number variants [CNVs]), and “balanced” SVs that rearrange genomic segments without corresponding dosage alterations (*e.g.,* inversions and translocations) (Figure 1a).^10^ Other common forms of SVs include mobile elements such as transposons that insert themselves throughout the genome,^11^ and multiallelic CNVs (MCNVs) that may exist at high copy numbers.^12^ Beyond these canonical classes, more exotic species of complex SVs exist in all individuals.^13,14^ These variants do not conform to a single canonical class, and instead involve two or more SV signatures in a single mutational event interleaved within the same allele, ranging from CNV-flanked inversions (*e.g.*, dupINVdup) to rare instances of localized chromosome shattering, such as chromothripsis.^15,16^ The variant spectrum of germline SVs in all humans is therefore broad, as is their influence on genome structure and function.

**Figure 1.**
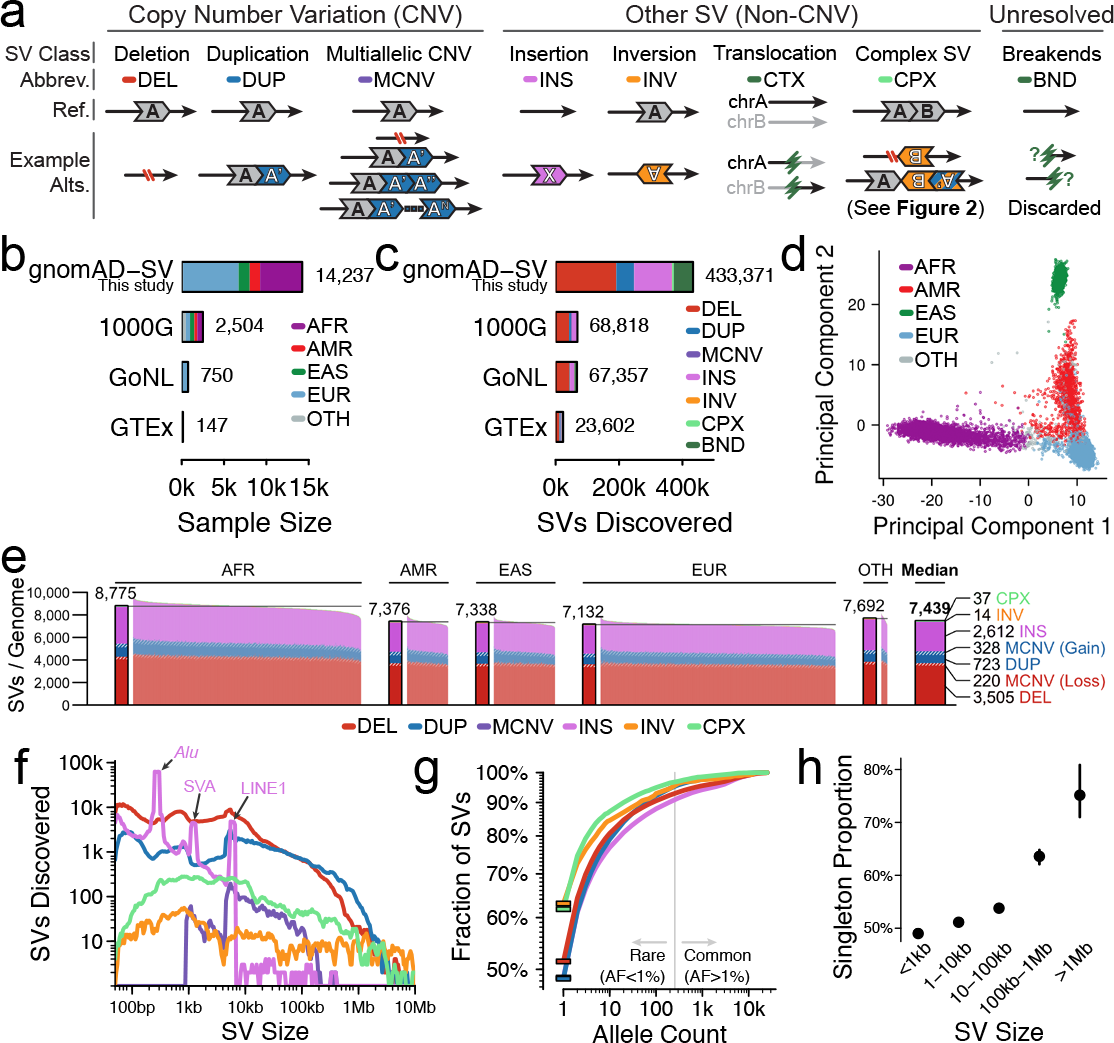
Properties of SVs across human populations. (**a**) SV classes catalogued in this study. Complex SVs were further categorized into 11 subclasses (see **Figure 2**). We also documented unresolved non-reference sequence junctions (i.e., breakends; BND), but they were excluded from all analyses. (**b**) After sample quality control, we processed 14,237 samples from four major continental populations: African/African-American (AFR), Latino (AMR), East Asian (EAS), and European (EUR). A small subset of samples came from admixed or other populations (OTH). Three publicly available WGS-based SV datasets are included for comparison (1000 Genomes Project [1000G], ~7× coverage; Genome of the Netherlands Project [GoNL], ~13× coverage; Genotype-Tissue Expression Project [GTEx], ~50× coverage).^1,20,29^ (**c**) We discovered 433,371 SVs, and provide counts from prior studies for comparison.^1,20,29^ (**d**) A principal component analysis of common SV genotypes separated samples along axes corresponding to genetic ancestry. (**e**) The median genome harbored 7,439 SVs. (**f**) Most SVs were small. Expected insertion peaks are marked at ~300 bp, ~2.1 kb, and ~6 kb, corresponding to three classes of mobile element insertion (Alu, SVA, and LINE1, respectively). (**g**) Most SVs were rare (AF<1%), and 49.8% of SVs were singletons (solid bars). (**h**) AFs were inversely correlated with SV size, which was accounted for in all subsequent analyses. Color codes are consistent between panels a, c, e-h, and between panels b and d.

While SVs alter more nucleotides per genome than single nucleotide variants (SNVs) and small insertion/deletion variants (indels; <50 bp),^1^ surprisingly little is known about their mutational spectra, patterns of natural selection, and functional impact on a global scale. This paucity of population-scale characterization of SVs to date is attributable to several factors, including the limited availability of deep coverage whole-genome sequencing (WGS) datasets and the myriad technical challenges of SV ascertainment. In contrast to the established gold-standard methods for profiling SNVs and indels from WGS data, like the Genome Analysis Toolkit (GATK),^17^ the uniform detection of SVs from short-read WGS has presented a much greater challenge.^18,19^ Analyses of SVs require specialized computational methods that consider multiple SV signatures, and even high-coverage short-read WGS fails to capture a component of the variant spectrum accessible to more expensive niche data types such as long-read WGS, optical mapping, or strand-specific sequencing.^18,19^ Current population references of SVs from WGS are thus restricted to the 1000 Genomes Project (N=2,504; 7× sequence coverage) or smaller European-centric cohorts.^1,20^ These references are dwarfed by contemporary resources for coding SNVs and indels such as the Exome Aggregation Consortium (ExAC) and its second iteration, the Genome Aggregation Database (gnomAD), which have jointly analyzed >140,000 exomes.^21,22^ Publicly available resources like ExAC and gnomAD have transformed most aspects of population genetics and disease association research, including defining a set of genes constrained against predicted loss-of-function (pLoF) variation,^21,23^ and have become integral in the medical interpretation of small coding variants.^24^ As short-read WGS is becoming the prevailing platform for large-scale human disease studies, and will likely displace conventional technologies in diagnostic screening, there is a critical need for similar resources of SVs across diverse global populations.

In this study, we developed gnomAD-SV, a reference map of SVs from WGS of 14,891 samples with an average coverage of 32× aggregated as part of gnomAD. Our analyses revealed diverse mutational patterns among SVs, and principles of selection acting against reciprocal dosage changes in genes and noncoding *cis*-regulatory elements. We found that SVs contributed approximately 25-29% of all rare protein-truncating events accessible to short-read WGS per genome, and that 0.18% of individuals in the general population harbored a clinically reportable SV that is likely to influence phenotype. These reference maps have been directly incorporated into the gnomAD browser (http://gnomad.broadinstitute.org) with no restrictions on reuse, and can be mined for new insights into genome biology and will provide an open resource for interpretation of SVs in diagnostic screening.

## RESULTS

### SV discovery and genotyping

We analyzed 14,891 samples in gnomAD-SV, of which 14,237 (95.6%) passed all data quality thresholds (**Supplementary Tables 1** and **SupplementaryFigure 1**). Samples were aggregated across numerous large-scale sequencing projects, and collectively represented a general adult population depleted for severe Mendelian diseases (median age = 49 years; **Supplementary Figure 2**). This gnomAD-SV reference included 46.1% European (N=6,559), 34.9% African/African-American (N=4,969), 9.2% East Asian (N=1,307), and 8.7% Latino (N=1,232) samples, as well as 1.2% samples from admixed or other populations (N=170; **Figure 1)**. Following SV discovery (described below) and family-based analyses of 970 parent-child trios as a quality assessment, we pruned all first-degree relatives from the cohort, retaining 12,653 unrelated genomes for subsequent analyses.

We discovered and genotyped SVs using a cloud-based version of a multi-algorithm pipeline for short-read WGS (**Supplementary Figure 3**), which has been previously detailed in a study of 519 autism quartet families.^25^ In brief, this pipeline integrated four orthogonal SV signatures to delineate variants across the size and allele frequency (AF) spectrum accessible to short-read WGS, including six classes of canonical SVs (**Figure 1a**; deletions, duplications, MCNVs, inversions, insertions, translocations) and 11 subclasses of complex SVs (**Figure 2**).^14^ We augmented this pipeline with new methods to account for the technical heterogeneity of aggregated WGS datasets (**Extended Data Figure 1** and **Supplementary Figures 4-5**). In total, these methods discovered 433,371 SVs (**Figure 1c** and **Supplementary Table 3**). We further pruned this dataset for the thousands of incompletely resolved non-reference breakpoint junctions per genome that are labeled by some algorithms as ‘breakends’ (BNDs; **Figure 1a**). These BNDs lack interpretable alternate allele structures for biological annotation, substantially inflated our variant counts (13.9% of all SVs detected), were enriched in false positives (**Extended Data Figure 2a**),^25^ and cannot be interpreted for functional impact, so we removed them from our final dataset (335,470 completely resolved SVs; **Supplementary Table 3**). The gnomAD-SV callset is freely available as a resource for the community via the gnomAD browser (https://gnomad.broadinstitute.org) and NCBI dbVar (accession nstd166).

**Figure 2.**
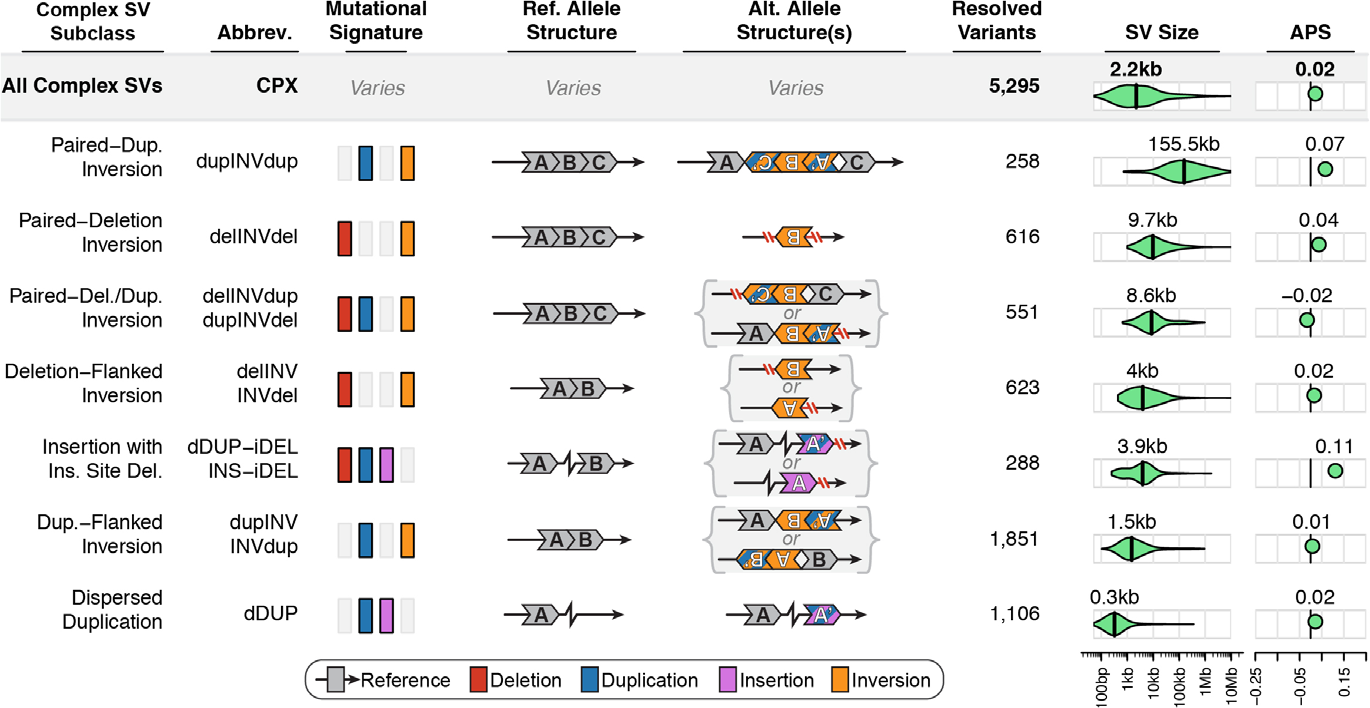
Complex SVs are abundant in the human genome. We discovered and fully resolved 5,295 complex SVs across 11 mutational subclasses, 73.7% of which involved at least one inversion. Each subclass is detailed here, including their mutational signatures, non-reference allele structures, abundance, sizes, and allele frequencies. For clarity, five pairs of subclasses have been collapsed into single rows due to mirrored or highly similar alternate allele structures (e.g., delINV vs INVdel). Two complex SVs that did not conform to any subclass are not included in this table (e.g., see **Figure 6e** and **Extended Data Figure 8**). APS = adjusted proportion of singletons.

There are currently no universally accepted, gold-standard benchmarking procedures for SV datasets from WGS, so we evaluated the technical qualities of gnomAD-SV using seven orthogonal approaches. These analyses are provided in complete detail in **Extended Data Figures 2**–**3**, **Supplementary Figures 6-12**, **Supplementary Tables 4-5**, and **Supplementary Note 1**. Overall, we found comparable specificities for gno-mAD-SV and our previous application of the same methods to 519 autism quartets, where we attained a 97% molecular validation rate for all *de novo* SV predictions.^25^ To highlight just a few measures in gnomAD-SV, we observed a Mendelian violation rate of 3.8% and a heterozygous *de novo* rate of 3.0% on average across 970 parent-child trios, which reflects a combination of false positives in children and/or false negatives in parents because we expect less than one true *de novo* SV per genome.^1,25,26^ We also found that 86% of SVs were in Hardy-Weinberg equilibrium, and common SVs were generally in strong linkage disequilibrium (LD) with nearby SNVs/indels (median peak R^2^ = 0.85). Finally, we leveraged matched long-read WGS data available for four individuals to perform *in silico* confirmation of our SVs predicted from short-read WGS.^19,27,28^ These analyses yielded a confirmation rate of 94.0% for SVs with break-point-level read evidence (92.8% of all SVs), and revealed that 59.8% of breakpoint coordinates from the gnomAD-SV callset were accurate within a single nucleotide of the long-read data, while 75.9% were accurate within ±10bp. In conclusion, despite the limitations of short-read WGS, the seven benchmarking approaches we applied here suggest that these data conform to many fundamental principles of population genetics, including Mendelian segregation, Hardy-Weinberg equilibrium, population stratification, and linkage disequilibrium, and that gnomAD-SV is sufficiently sensitive and specific to provide a contemporary resource for most applications in human genomics.

### Insights into population genetics and genome biology

Investigation of population substructure from SVs followed the expectations set by human demographic history,^29^ with the top three principal components providing clear separation between populations (**Figure 1d** and **Supplementary Figure 13**). African/African-American samples exhibited the greatest genetic diversity (**Figure 1e**) and their common SVs exhibited weaker LD with nearby SNVs and indels than Europeans (**Supplementary Figure 7**). East Asian genomes featured the highest levels of homozygosity (**Extended Data Figure 4a-d**). The spectrum of SVs present in gnomAD-SV was diverse: we completely resolved 5,295 complex SVs across 11 mutational subclasses, of which 3,901 (73.7%) involved inverted segments (**Figure 2**), confirming prior predictions that most inversion variation accessible to short-read WGS comprises complex SVs rather than canonical inversions.^1,30^ Across all SV classes, most SVs were small (median SV size = 331 bp; **Figure 1f**) and rare (AF < 1%; 92% of SVs; **Figure 1g**), with nearly half of all SVs (49.8%) appearing as “singletons” (*i.e.*, only one allele observed across all samples). While singleton proportion varied by SV class, it was strongly dependent on SV size across all classes, suggesting that the amount of genetic material rearranged is a principal determinant of selection against most SVs (**Figure 1h** and **Extended Data Figure 5a**).

Mutation rates for SVs have remained difficult to quantify due to the limited resolution of conventional technologies, the technical challenges of SV discovery from short-read WGS, and the frequent use of cell line-derived DNA in population studies.^1,26^ Given that nearly all samples in this study (99.3%) were sequenced from whole blood-derived DNA, we used the Watterson estimator^31^ to project a mean mutation rate of 0.29 *de novo* SVs per generation in regions of the genome accessible to short-read WGS (95% confidence interval: 0.13-0.44), or roughly one new SV every 2-8 live births, with mutation rates varying markedly by SV class (**Figure 3a**). While this imperfect method approximates mutation rates from aggregated genetic data across unrelated individuals, we previously demonstrated comparable rates from molecularly validated observations in WGS analyses of 519 quartet families.^25^ Like mutation rates, the distribution of SVs throughout the genome was non-uniform, significantly correlated with numerous repetitive sequence contexts, and particularly enriched near centromeres and telomeres (**Supplementary Figure 16**).^32^ These trends were strongly dependent on SV class. For instance, biallelic deletions and duplications were predominantly enriched at telomeres, whereas MCNVs were preferentially enriched in centromeric segmental duplications (**Figure 3b-d**). Given the reduced sensitivity of short-read WGS in repetitive and low-complexity sequences, gnomAD-SV certainly underestimates the true mutation rates and distributions of SVs, which are likely to be refined by population-scale applications of long-read genome assembly methods.^33,34^ Nevertheless, these analyses clearly implicate multiple aspects of chromosomal context and SV class in driving SV mutation rates throughout the genome.

**Figure 3.**
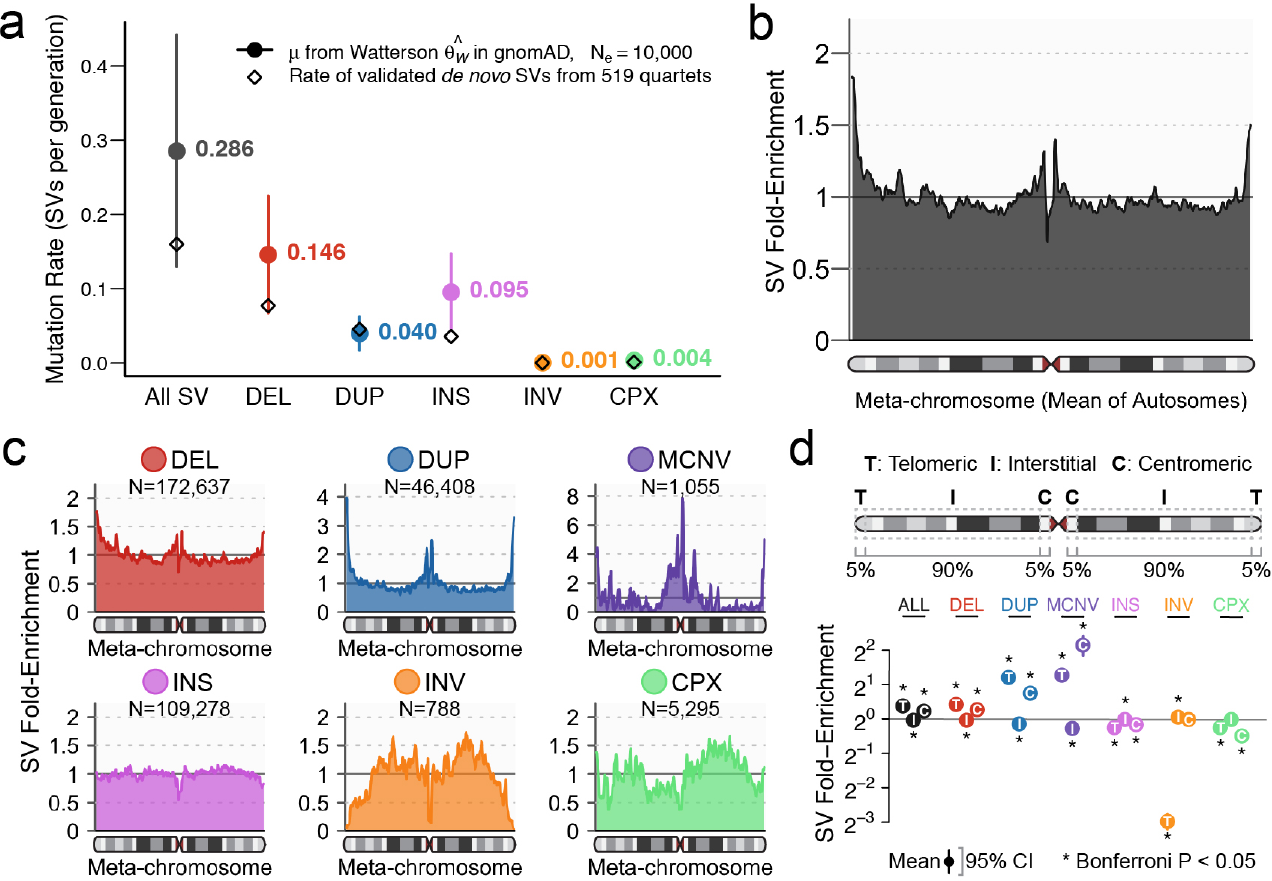
Genome-wide mutational patterns of SVs. (**a**) Mutation rates (μ) from the Watterson estimator for each SV class.^32^ Bars represent 95% confidence intervals. Rates of molecularly validated de novo SVs from 519 quartet families are provided for comparison.^25^ (**b**) Smoothed enrichment of SVs per 100 kb window across the average of all autosomes normalized by chromosome arm length (a “meta-chromosome”; also see **Supplementary Figure 16**). (**c**) The distribution of SVs along the meta-chromosome was dependent on variant class. (**d**) Biallelic CNVs were predominantly enriched at telomeres, MCNVs were predominantly enriched at centromeres, and canonical and complex inversions were depleted near telomeres. P-values computed using a two-sided t-test; bars correspond to 95% confidence intervals (CIs).

### Dosage sensitivity of protein-coding genes and noncoding regulatory elements

Due to their size and mutational diversity, SVs can have varied consequences on protein-coding genes (**Figure 4a** and **Supplementary Figure 17**).^7^ All classes of SVs can result in pLoF, either by deletion of coding nucleotides or alteration of open-reading frames. Coding duplications can result in copy-gain (CG) of entire genes, or duplication of a subset of exons contained within a gene, referred to here as intragenic exonic duplication (IED). The average genome in gnomAD-SV harbored a mean of 179.8 genes altered by biallelic SVs (144.3 pLoF, 24.3 CG, and 11.2 IED), of which 11.6 were predicted to be completely inactivated by homozygous biallelic pLoF SVs (**Figure 4b** and **Extended Data Figure 4e-h**). When restricted to rare (AF < 1%) SVs, the mean genome had 10.2 altered genes (5.5 pLoF, 3.4 CG, and 1.3 IED), all effectively heterozygous. By comparison, SNV and indel analyses in gnomAD estimated 122.4 pLoF SNVs/indels per genome, of which 16.3 were rare,^22^ suggesting that between 25-29% of all rare pLoF events per genome are contributed by SVs, although this fraction is likely to be upwardly revised as the sensitivity of SV detection improves with emerging technologies.

**Figure 4.**
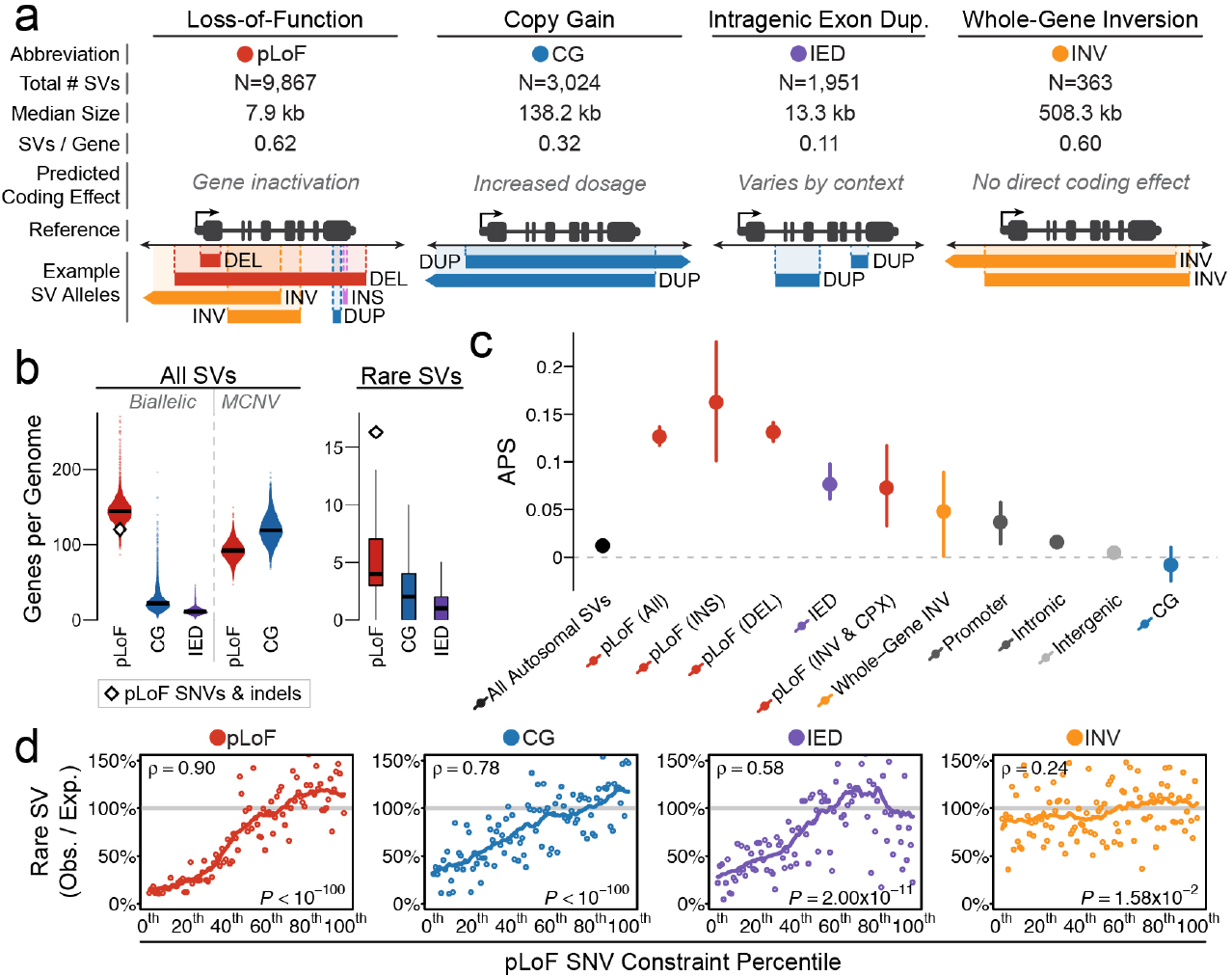
Pervasive selection against SVs in genes mirrors patterns observed from coding point mutations. (**a**) Four categories of gene-overlapping SVs, with counts of SVs in gnomAD-SV. (**b**) Distributions of genes altered by SVs per genome. (**c**) Autosomal SVs that overlap genes were enriched for singleton variants (a proxy for the strength of selection^20^) above baseline of all SVs genome-wide, and explicitly intergenic SVs (also see **Extended Data Figure 5c-d**). Bars indicate 100-fold bootstrapped 95% confidence intervals. (**d**) We evaluated the relationship of constraint against pLoF SNVs versus the four categories of gene-overlapping SVs from (a).^20^ Each point represents the total of ~175 genes, which have been ranked by SNV constraint. Correlations were assessed with a Spearman test. Solid lines represent 21-bin rolling means. See **Supplementary Figure 10** for comparisons to missense SNV constraint.

The degree to which selection acts globally on SVs within and outside of coding regions of the genome remains a fundamental question in human genetics. One approach to quantifying selection relies on inference from site-frequency spectra. Specifically, the proportion of singleton variants has been established as a proxy for strength of selection.^21,22^ However, given the strong correlation between SV size and AF (**Extended Data Figure 5a**), direct comparisons of raw singleton proportions between groups of SVs are inherently confounded by SV size and other factors. Therefore, we developed a metric to account for SV class, size, genomic context, and other technical covariates (referred to herein as <u>A</u>djusted <u>P</u>roportion of <u>S</u>ingletons [APS]; **Extended Data Figure 5b** and **Supplementary Figure 14**). Under this normalized APS metric, a value of zero for a group of SVs corresponds to a singleton proportion comparable to intergenic SVs, whereas values greater than zero reflect increased proportions of singleton variants—and therefore increased selection—similar to the “MAPS” metric used for SNVs.^21,22^ Using this APS model, we found signals of pervasive selection against nearly all classes of SVs that overlap genes, including intronic SVs and pLoF SVs as small as partial-exon deletions (**Figure 4c**, **Extended Data Figure 6a-d**, and **Supplementary Figure 18**). The exception was CG duplications, which showed no additional negative selection beyond what could already be explained by sizes vastly larger than seen for duplications that did not encompass entire genes (median CG duplication size = 134.8kb versus median non-CG duplication size = 2.7kb; P < 10^−100^, one-tailed Wilcoxon test). This result may be indicative of possible overcorrection for SV size in our APS model, but it is also consistent with the diverse evolutionary roles of gene duplication events, including positive selection acting on a subset of CGs in humans.^35,36^ While further methods development will continue to refine such predictions, these data show that SVs represent a substantial fraction of all gene-altering variants per genome, and widespread selection acts to remove most gene-altering SVs from the population.

Beyond the global impact of selection against coding variation, methods have recently been developed to quantitate selection on functional variation on a per-gene basis, such as the probability of LoF intolerance (pLI). These scores have become core resources in human genetics.^21–23^ Existing metrics like pLI are reliant on SNVs, and while previous studies have attempted to compute similar scores using large CNVs detected by microarray or to correlate deletions with pLI,^37,38^ no gene-level metrics comparable to pLI exist for SVs at WGS resolution. To gain insight into this problem, we estimated the number of rare SVs expected per gene while adjusting for gene length, exon-intron structure, and genomic context. This model is imperfect, as expectations can be influenced by many known and unknown covariates, and SVs are too sparse at current sample sizes to derive precise gene-level estimates of SV constraint. Nevertheless, we found strong concordance between pLoF constraint metrics from gnomAD SNV analyses and the depletion of rare pLoFs in gnomAD-SV (**Figure 4d**; Spearman’s rho = 0.90).^22^ This result was also true of missense constraint (**Supplementary Figure 19**), as expected given the strong correspondence between missense and pLoF constraint. Notably, a comparable positive correlation was also observed between CG from rare SVs and pLoF constraint from SNVs (rho = 0.78), while a weaker yet significant correlation was also detected for IED (rho = 0.58). When we cross-examined these relationships using APS, we found an inverse correlation between the proportion of singleton SVs and SNV constraint across all functional categories of SVs (**Extended Data Figure 6f**). These comparisons confirm that selection against most classes of gene-altering SVs is consistent with patterns observed for SNVs and indels. They further suggest that constraint metrics like pLI, which are derived from pLoF point mutations, in fact capture a general correspondence between haploinsufficiency and triplosensitivity—*on average*—for a large fraction of genes in the genome. It therefore appears that many highly pLoF-constrained genes are not only sensitive to pLoF, but also likely intolerant to increased dosage and other functional alterations more broadly.

In contrast to the well-established effects of coding variation, the impact of noncoding SVs on regulatory elements is mostly unknown. Examples of SVs with strong noncoding effects are scarce in humans and model organisms,^39–41^ though recent studies have shown that noncoding SVs are relevant for gene regulation and disease.^42,43^ We explored noncoding dosage sensitivity across a spectrum of 14 regulatory element classes, ranging from high-confidence experimentally validated enhancers to large databases of computationally predicted elements. We found that non-coding CNVs overlapping most element classes had elevated singleton proportions (i.e., APS), though no SV class matched the APS observed for protein-coding pLoFs (**Figure 5a**). Conversely, noncoding CNVs that did not overlap any annotated elements featured an APS not significantly different from zero, reflecting relatively neutral variation. In general, the effects from noncoding deletions were stronger than noncoding duplications, and CNVs predicted to delete or duplicate entire elements were under stronger selection than CNVs with only partial element overlap (**Figure 5b**). We also observed that primary sequence conservation was correlated with singleton proportion across all noncoding CNVs (**Figure 5c-d**), which lays the groundwork for functionally predictive models and interpretation frameworks for noncoding SVs. Collectively, these results mirrored trends we observed for protein-coding SVs, and can be interpreted to imply weak, widespread selection against CNVs altering most classes of annotated regulatory elements.

**Figure 5.**
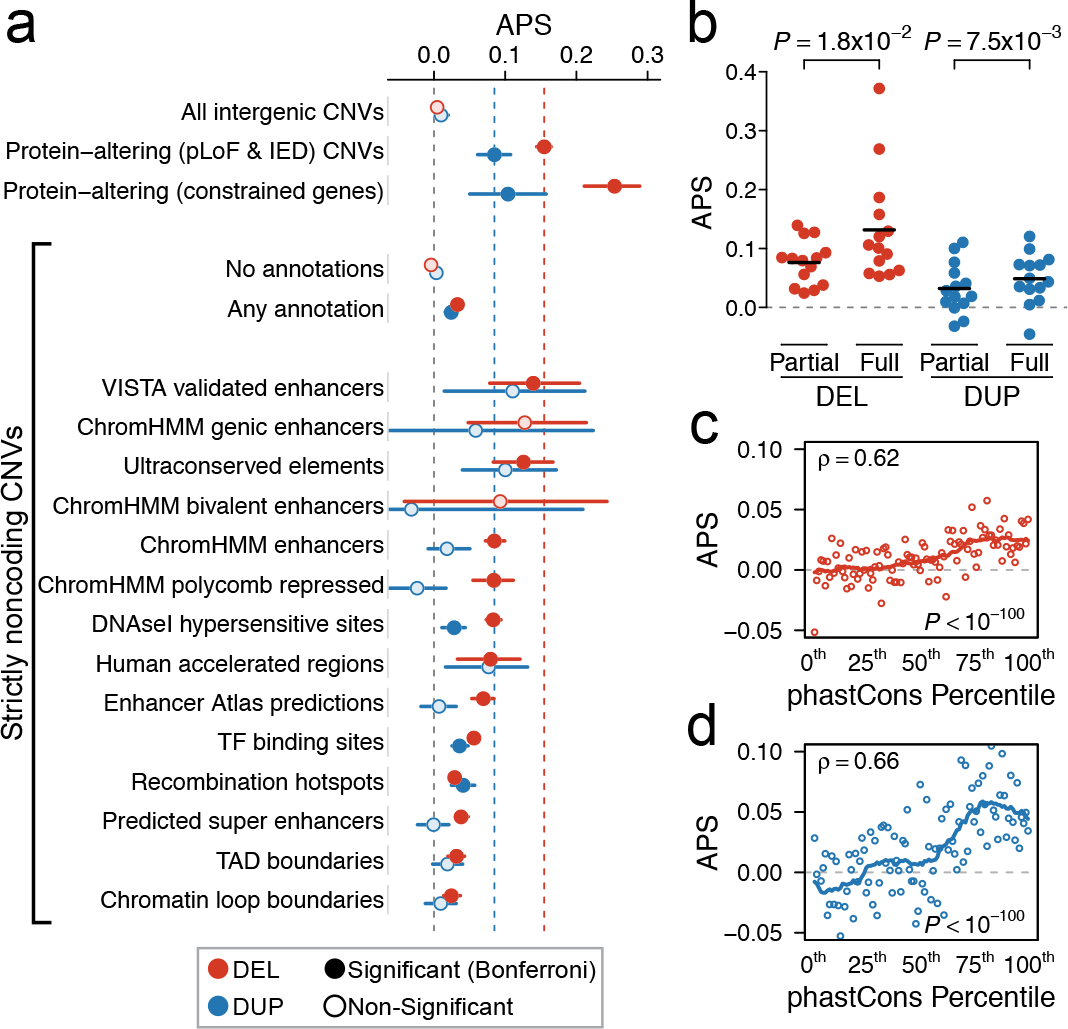
Dosage sensitivity in the noncoding genome. (**a**) Estimated strength of selection (APS) for noncoding CNVs intersected with 14 categories of noncoding elements. Bars reflect 95% confidence intervals from 100-fold bootstrapping. Protein-altering (pLoF & IED) CNVs provided for reference. Each category was compared to the expected APS of 0 for neutral variation using a one-tailed t-test. Categories surpassing Bonferroni-corrected significance for the 32 noncoding comparison performed here are shown with dark shaded points. (**b**) Across all 14 annotation classes evaluated in (a), CNVs that completely covered elements (“full”) had significantly higher average APS values than CNVs that only partially covered elements (“partial”). P-values calculated using a two-tailed paired two-sample t-test. (**c-d**) Correlations between primary sequence conservation and APS for noncoding (c) deletions and (d) duplications. Here, noncoding CNVs were divided into percentile bins based on the sum of the phastCons scores overlapped by the CNV, and the relationships between APS and phastCons percentile were evaluated with a Spearman rank correlation test. Solid lines represent 21-bin rolling means.

### Relevance to disease association and clinical genetics

Most large-scale disease association efforts have focused on genotyping common SNVs in genome-wide association studies (GWAS).^44^ Taking advantage of the sample size and resolution of gnomAD-SV, we evaluated whether SNVs associated with human traits from GWAS might be in LD with functional SVs not directly genotyped during GWAS.^42^ We identified 15,634 common SVs (AF > 1%) in strong LD (R^2^ ≥ 0.8) with at least one common SNV or indel (**Supplementary Figure 7** and **Supplementary Table 6**), 14.8% (2,307/15,634) of which matched a reported association from the NHGRI-EBI GWAS catalog or a recent analysis of 4,203 phenotypes in the UK BioBank.^45–47^ Common SVs in LD with GWAS associations were enriched for genic SVs across multiple functional categories, and included intriguing candidate SVs such as a 336 bp *Alu* deletion of a thyroid enhancer in the first intron of *ATP6V0D1* at a hypothyroidism-associated locus (**Extended Data Figure 7**).^46^ We also found matches for several previously proposed causal SVs tagged by common SNVs, including pLoF deletions of *CFHR3/CFHR1* in nephropathies and of *LCE3B/ LCE3C* in psoriasis.^48,49^ These results support the value of imputing SVs from WGS into future studies of human phenotypes, and for the eventual unification of SNVs, indels, and SVs in all trait association studies.

As genomic medicine advances toward diagnostic screening at sequence resolution, WGS-based methods for SV detection and publicly accessible WGS references will be indispensable for variant discovery and interpretation. In the context of variant discovery, one subset of disease-associated SVs is particularly important: genomic disorders (GDs). GDs are recurrent CNVs mediated by flanking homologous segmental duplications, and collectively represent one of the most common genetic causes of developmental disorders.^50^ Accordingly, a chromosomal microarray (CMA) to detect large CNVs is currently recommended as the first-tier genetic diagnostic screen for developmental disorders of unknown etiology.^51^ Therefore, the ability of WGS to reliably discover these repeat-mediated CNVs is critical. Using gnomAD-SV, we evaluated our ability to detect GD CNVs is critical. Using gnomAD-SV, we evaluated our ability to detect GD CNVs in WGS data by calculating CNV carrier frequencies from gnomAD-SV for 49 GDs across 10,047 unrelated samples with no known neuropsychiatric disease. We found that CNV carrier frequencies from WGS in gnomAD-SV were consistent with those reported from CMA in the UK BioBank^52^ (UKBB; R^2^ = 0.669; P = 7.38 × 10^−13^; Pearson correlation test; **Figure 6a**, **Supplementary Table 7**, and **Supplementary Figure 20**). GD carrier frequencies did not vary significantly among populations in gnomAD-SV, with the exception of a single CNV—duplications of *NPHP1* at 2q13—for which carrier frequencies in East Asian samples were 2.5-to-4.6-fold higher than in other populations, further highlighting the potential for disease risk interpretations to be confounded by the limited diversity of existing reference datasets (**Supplementary Figure 21**).

**Figure 6.**
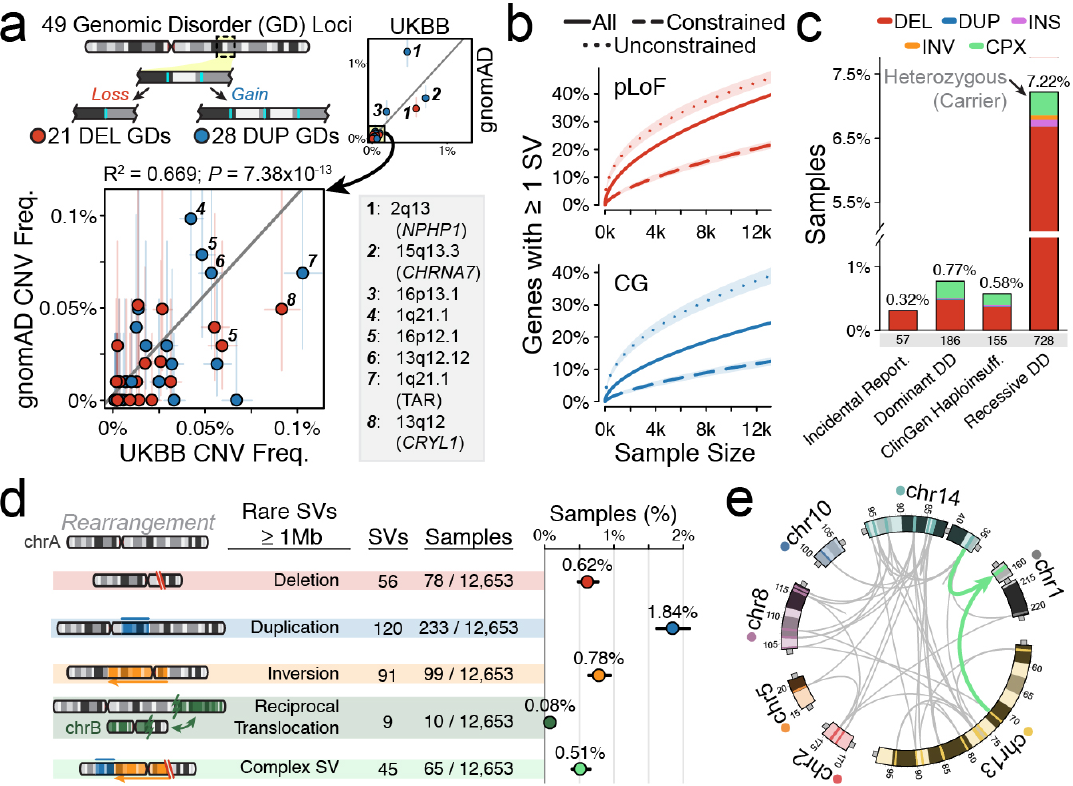
gnomAD-SV as a resource for clinical WGS interpretation. (**a**) Filtering SVs against gnomAD-SV reduces individual genomes to ~13 singleton variants at current sample sizes. (**b**) At least one pLoF or CG SV was detected in 40.4% and 23.5% of all autosomal genes, respectively. “Constrained” and “unconstrained” includes the least and most constrained 15% of all genes based on pLoF SNV observed:expected ratios, respectively.^20^ (**c**) Up to 1.3% of genomes in gnomAD-SV harbored a very rare (AF<0.1%) pLoF SV in a medically relevant gene across several gene lists.^38–40^ Manual review of all very rare pLoF SVs indicated that 0.24% of genomes carry a pathogenic or likely pathogenic variant in a clinically reportable gene for incidental findings.^38^ We also found that 9.5% of genomes carried pLoF SVs of recessive DD genes in the heterozygous state.^39^ (**d**) We found 308 rare autosomal SVs ≥ 1Mb, revealing that ~3.1% of genomes carry a large, rare chromosomal abnormality. Bars represent binomial 95% confidence intervals. (**e**) An extremely complex SV involving at least 49 breakpoints that localized in clusters across seven chromosomes in a single individual, yielding largely balanced derivatives, reminiscent of chromothripsis (see also **Extended Data Figure 8**). Chromosome coordinates provided as Mb.

In the context of variant interpretation, the current gnomAD-SV resource will permit a screening threshold of AF < 0.1% when matching on ancestry to the populations sampled here, and AF < 0.004% globally. gno-mAD-SV can also aid in gene-level interpretation: we catalogued at least one SV resulting in pLoF or CG for 36.9% and 23.7% of all autosomal genes, respectively, and 490 genes with at least one homozygous pLoF SV (**Figure 6b**, **Extended Data Figure 6e**, and **Supplementary Figure 22**). However, these data are still sparse compared to data for SNVs and indels, where analyses of the 140,000 gnomAD exomes documented at least one pLoF SNV for 95.8% of all genes.^22^ Despite this relative sparsity, we benchmarked carrier rates for several categories of medically relevant variants in gnomAD-SV. First, 0.32% of samples in gnomAD-SV carried a very rare (AF < 0.1%) SV resulting in pLoF of a gene for which incidental findings are clinically reportable, roughly half of which (*i.e.*, 0.18% of all samples) would meet criteria for classification as pathogenic or likely pathogenic (**Figure 6c**).^53^ Second, we observed that 7.22% of individuals were heterozygous carriers of rare pLoF SVs in known recessive developmental disorder genes.^54^ Third, we estimated that 3.8% of the general population (95% CI: 3.2-4.6%) carries at least one very large (≥1Mb) rare autosomal SV, roughly half of which (45.2%) are balanced or complex (**Figure 6d**). Among these was an example of highly complex localized chromosome shattering involving at least 49 breakpoints yet resulting in largely balanced products reminiscent of chromothripsis.^14–16,55^ The variant was identified in an adult from the general population with no indication of severe disease and no known DNA repair defect (**Figure 6e** and **Extended Data Figure 8**). Collectively, these analyses demonstrate multiple avenues by which gnomAD-SV can augment future disease association studies, population genetic screens, and clinical interpretation of SVs across a broad spectrum of variant classes and study designs.

## DISCUSSION

Human genetic research and clinical diagnostics are becoming increasingly invested in defining the complete landscape of variation in individual genomes. Ambitious international initiatives to generate short-read WGS in hundreds of thousands of individuals from complex disease cohorts have underwritten this goal,^56–59^ and millions of genomes from unselected individuals will be sequenced in the coming years from national biobanks.^60,61^ A central challenge to these efforts will be the uniform analysis and interpretation of all variation accessible to short-read WGS, particularly SVs, which are frequently cited as a source of added value offered by WGS over conventional technologies.^62^ Indeed, early efforts to deploy WGS in cardiovascular disease, autism, and type 2 diabetes were largely consistent in their analyses of SNVs using GATK, but all studies have differed in their analyses of SVs.^25,37,50,57–59,63,64^ Thus, while ExAC and gnomAD have catalyzed remarkable advances in medical and population genetics, the same opportunities for new discovery and translational impact have not yet been realized for SVs. Although gnomAD-SV is by no means comprehensive, the nearly half-million SVs it contains were derived from WGS methods and a reference genome that match those currently used in many research and clinical settings.

Most foundational assumptions about human genetic variation were consistent between SVs and the SNVs/indels from the gnomAD exome study,^22^ most notably that SVs segregate stably on haplotypes in the population and experience selection commensurate with their predicted biological consequences. This study also spotlights unique aspects of SVs, such as their remarkable mutational diversity, their varied functional impacts on coding sequence, and the strong selection against large and complex SVs. We provide resolved structures for over 5,000 complex SVs, and predict that SVs comprise at least 25-29% of all rare protein-truncating variation in each genome. These analyses demonstrate that gene-altering effects of SVs beyond pLoF parallel mutational constraints derived from analyses of SNVs, and that existing SNV constraint metrics are not specific to haploinsufficiency but underlie a more generalizable intolerance to alterations of both gene dosage and structure. Beyond the coding genome, we uncovered widespread but modest selection against noncoding dosage alterations of *cis*-regulatory elements, such as enhancers, ultraconserved elements, and chromatin domain boundaries. This finding represents one of the largest empirical assessments of noncoding dosage sensitivity in humans to date, but underscores two important corollaries: (1) that few—if any—classes of noncoding *cis*-regulatory elements are likely to experience selection as strong as for protein-truncating variants, and (2) that current WGS sample sizes are vastly underpowered to robustly identify individual constrained elements in the noncoding genome.

Technical barriers associated with short-read WGS preclude the establishment of a complete catalogue of SVs in gnomAD-SV. While the number of fully resolved SVs per genome in gnomAD-SV using the integration of multiple algorithms here (n = 7,439 SVs per genome from ~32× coverage WGS) is roughly twice that of existing references from short-read WGS, such as the 1000 Genomes Project (3,441 SVs from ~7× WGS) and the GTEx project (3,658 SVs from ~50× WGS),^1,42^ it is lower than estimates from recent long-read WGS analyses (24,825 SVs from ~40× long-read WGS).^19^ The technology and methods used here are thus blind to a disproportionate fraction of repeat-mediated SVs, and underestimate the true mutation rates within these hypermutable regions. Similarly, high copy state MCNVs often require specialized algorithms and manual curation to fully delineate their complicated haplotype structures,^12,65,66^ suggesting that the 1,055 MCNVs reported here are an incomplete portrait of extreme copy-number polymorphisms. We expect that emerging technologies, *de novo* assemblies, and graph-based genome representations are likely to expand our knowledge of SVs in repetitive sequences.^66,67^ Nevertheless, based on current estimates, 92.7% of known autosomal protein-coding nucleotides are not localized to simple- and low-copy repeats. Thus, catalogues of SVs accessible to short-read WGS across large populations—such as those presented here and in future releases of gnomAD-SV—will likely capture a majority of the most interpretable gene-disrupting SVs in humans.

The oncoming deluge of short-read WGS datasets has magnified the need for publicly available large-scale resources of SVs. In this study, we aimed to begin to bridge the gap between the existence of such references for SNVs/indels and those for SVs. While the dataset provided here significantly exceeds current references in terms of sample size and sensitivity, these data remain insufficient to derive accurate estimates of gene-level constraint, sequence-specific mutation rates, and intolerance to noncoding SVs. Nonetheless, these data provide an initial step toward these goals, and demonstrate the value of a commitment to open data sharing and joint analyses of aggregated datasets by the field. The gnomAD-SV resource has been made available without restrictions on reuse, and has been integrated into the widely adopted gnomAD Browser (https://gnomad.broadinstitute.org), where users can freely view, sort, and filter the SV dataset described here, as well as access future gno-mAD-SV releases. This resource will facilitate the continued development of methods for functional prediction of SVs, catalyze new discoveries in basic research, and provide immediate clinical utility for the interpretation of rare structural rearrangements in the human genome.

## METHODS & SUPPLEMENTARY INFO

There is supplementary information associated with this study, which includes detailed methods. These materials have been provided in a separate document, which will be linked directly from *bioRxiv*.

## ACKNOWLEDGEMENTS

This research and contributing authors were supported by resources from the Broad Institute, the National Institutes of Health (R01MH115957, R01HD081256, P01GM061354, R01HD091797, R01HD096326, R01MH111776 to MET; U01MH105669 to MJD, BN, and MET; P50HD028138 to BN, MET, and HB) and the Simons Foundation for Autism Research Initiative (SFARI #573206 to MET). We are grateful to all of the families at the participating Simons Simplex Collection (SSC) sites, as well as the SSC principal investigators. RLC was supported by NHGRI T32HG002295 and NSF GRFP #2017240332. HB was supported by NIDCR K99DE026824. MESA and the MESA SHARe project are conducted and supported by the National Heart, Lung, and Blood Institute (NHLBI) in collaboration with MESA investigators. Support for MESA is provided by contracts HHSN268201500003I, N01-HC-95159, N01-HC-95160, N01-HC-95161, N01-HC-95162, N01-HC-95163, N01-HC-95164, N01-HC-95165, N01-HC-95166, N01-HC-95167, N01-HC-95168, N01-HC-95169, UL1-TR-000040, UL1-TR-001079, and UL1-TR-001420. MESA Family is conducted and supported by the National Heart, Lung, and Blood Institute (NHLBI) in collaboration with MESA investigators. Support is provided by grants and contracts R01HL071051, R01HL071205, R01HL071250, R01HL071251, R01HL071258, R01HL071259, by the National Center for Research Resources, Grant UL1RR033176, and the National Center for Advancing Translational Sciences ULTR001881, and the National Institute of Diabetes and Digestive and Kidney Disease Diabetes Research Center (DRC) grant DK063491 to the Southern California Diabetes Endocrinology Research Center. We thank Tim Hefferon of the NIH National Center for Biotechnology Information for his help hosting gnomAD-SV on dbVar.

---

**Extended Data Figure 1.**
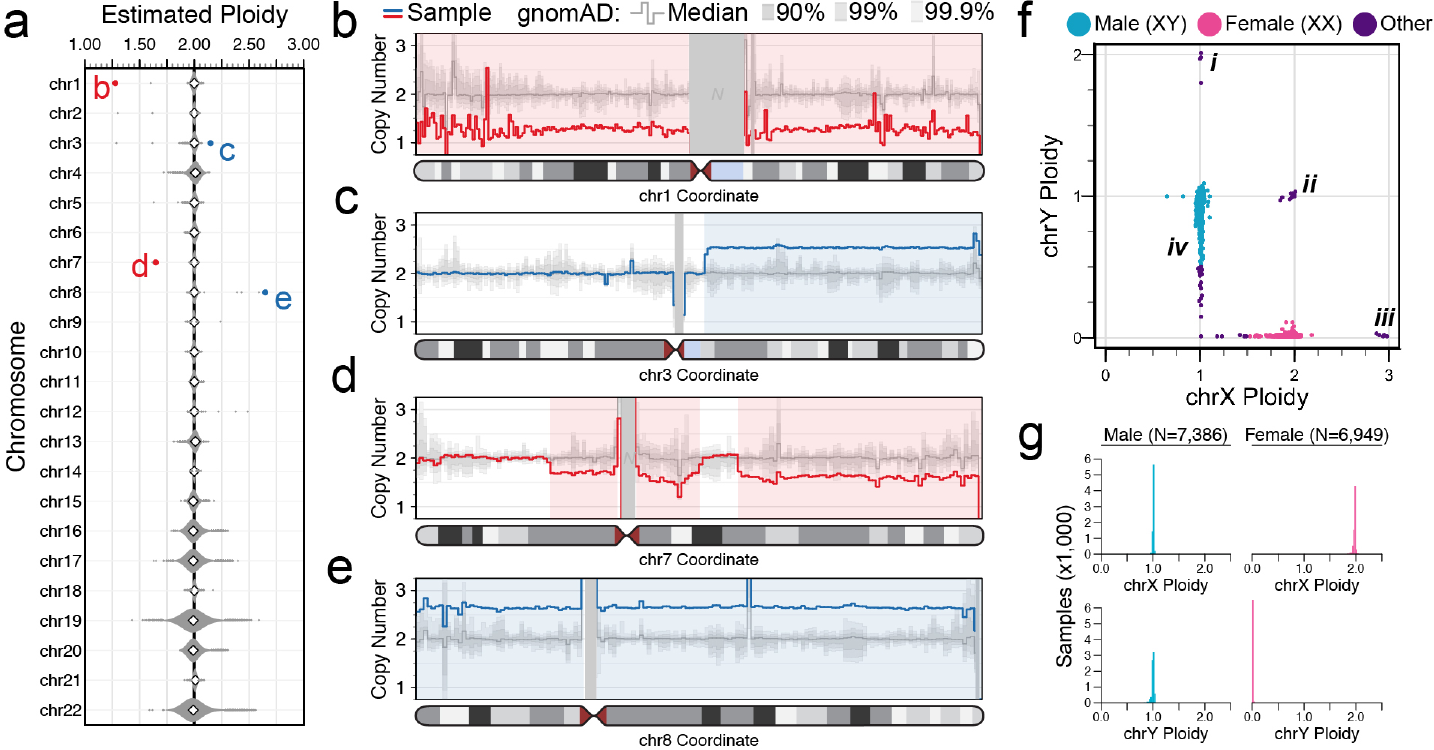
Detection of chromosome-scale dosage alterations. We estimated ploidy (i.e. whole-chromosome copy number) for all 24 chromosomes per sample. (**a**) Distribution of autosome ploidy estimates across 14,378 samples passing initial data quality thresholds. The outlier points marked in red and blue correspond to the samples highlighted in panels (b-e). (**b-e**) Samples with outlier autosome ploidy estimates typically harbored somatic or mosaic chromosomal abnormalities, such as somatic aneuploidy of chr1 (b) or chr8 (e), or large focal somatic or mosaic CNVs on chr3 (c) and chr7 (d). Each panel depicts copy-number estimates in 1Mb bins for each rearranged sample in red or blue. Dark, medium, and light grey background shading indicates the range of copy number estimates for 90%, 99%, and 99.9% of all gnomAD-SV samples, respectively, and the medium grey line indicates the median copy number estimate across all samples. Regions of unalignable N-masked bases >1Mb in the reference genome are masked with grey rectangles. (**f**) Sex chromosome ploidy estimates for all samples from (a). We inferred karyotypic sex by clustering samples to their nearest integer ploidy for sex chromosomes. Several abnormal sex chromosome ploidies are marked, including XYY (i), XXY (ii), XXX (iii), and mosaic loss-of-Y (iv). (**g**) The overwhelming majority of samples conformed to canonical sex chromosome ploidies.

**Extended Data Figure 2.**
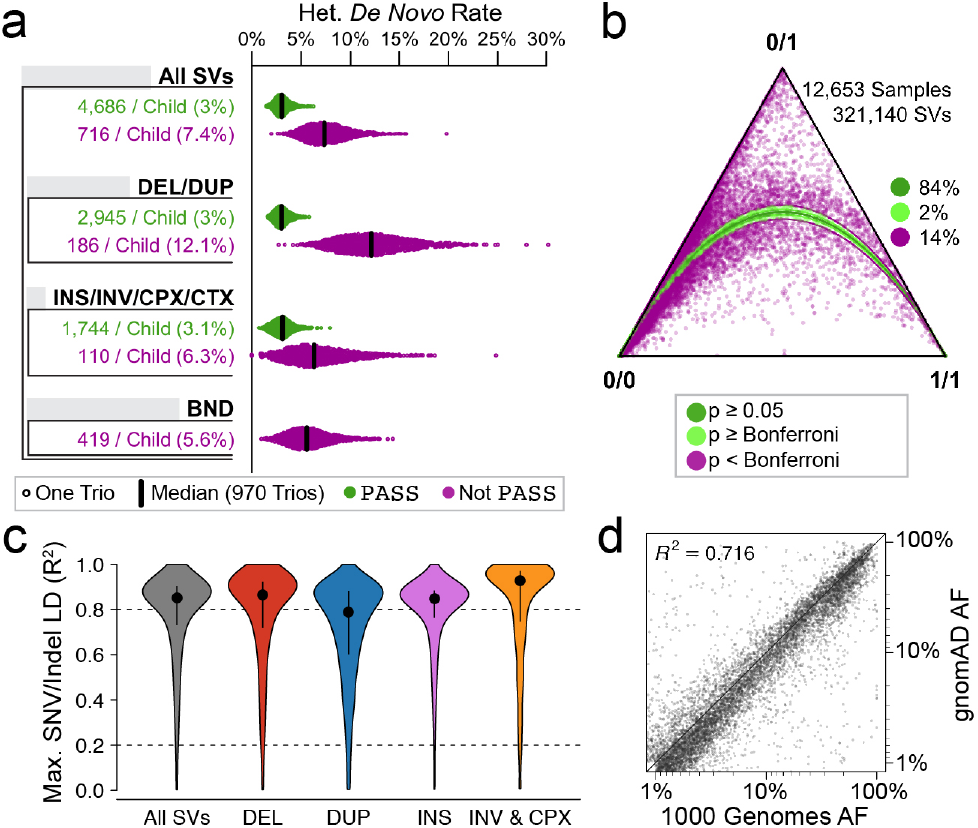
Benchmarking the technical qualities of the gnomAD-SV callset. We evaluated the quality of gnomAD-SV with seven orthogonal analyses detailed in **Supplementary Table 4** and **Supplementary Figures 6-9**. Four core analyses are presented here. (**a**) Apparent rates of de novo (i.e., spontaneous) heterozygous SVs per child across 970 parent-child trios. Given the expected mutation rate of SVs accessible to short-read WGS (<1 true de novo SV per trio; see also **Figure 3a**),^1,25^ effectively all de novo SVs represented a combination of false-positive genotypes in children and/or false-negative genotypes in parents. SVs passing all filters and included in the final gnomAD-SV callset (“PASS”) are shown in green. For comparison, variants that did not pass post hoc site-level filters (“Not PASS”) are also shown in purple. (**b**) Hardy-Weinberg Equilibrium (HWE) metrics for all biallelic SVs localized to autosomes. Vertex labels reflect genotypes: 0/0=homozygous reference; 0/1=heterozygous; 1/1=homozygous alternate, with all sites shaded by HWE p-value. (**c**) Linkage disequilibrium between SVs and SNVs/indels represented as cross-population maximum R^2^ after excluding repetitive and low-complexity regions (see **Supplementary Figure 7**). (**d**) AF correlation for common (AF>1%) SVs captured by both the 1000 Genomes Project and gnomAD-SV.^1^ A Pearson correlation coefficient is provided here.

**Extended Data Figure 3.**
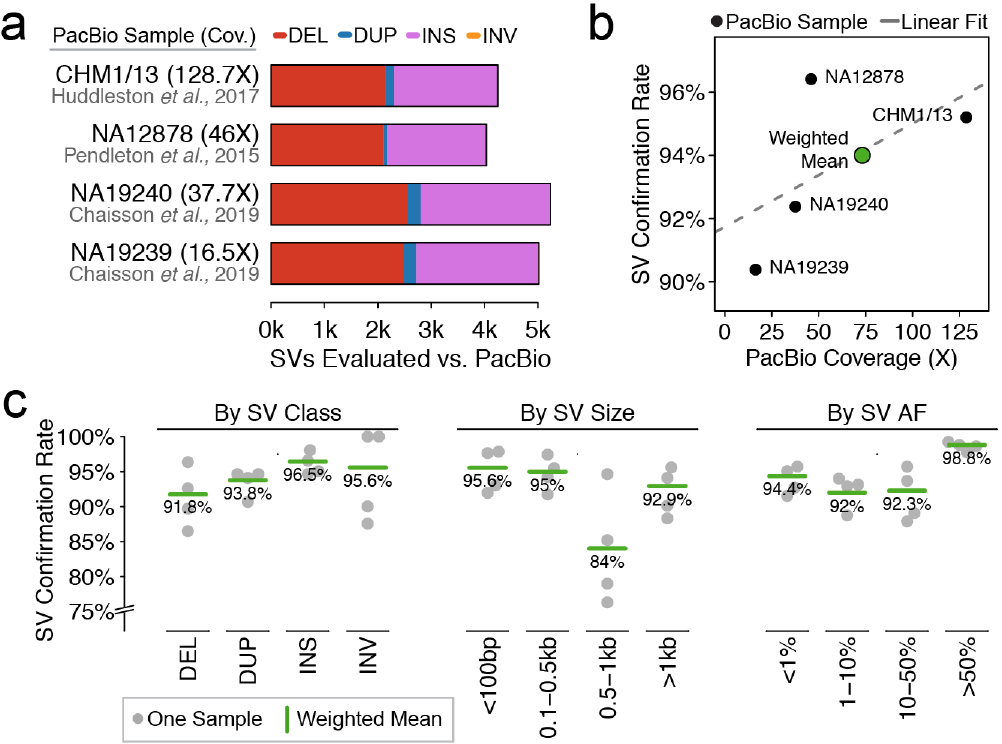
Evaluating the positive predictive value of gnomAD-SV with long-read WGS. We used Pacific Biosciences (PacBio) long-read WGS data available for four samples in this study to perform in silico confirmation to estimate positive predictive value (PPV) and breakpoint accuracy for SVs in gnomAD-SV (**Supplementary Figure 10**).^19,27,28^ (**a**) Counts of SVs evaluated per sample in this analysis. SVs were restricted to those with breakpoint-level read support (i.e., “split-read” evidence, 92.8% of all SVs) and also did not have breakpoints localized to annotated simple repeats or segmental duplications. (b) An iterative local long-read WGS realignment algorithm, VaPoR,^69^ was used to perform in silico confirmation of SVs predicted from short-read WGS in gnomAD-SV. As noted by the VaPoR developers,^69^ the performance of this approach was sensitive to the sequencing depth of long-read WGS data. Therefore, the weighted mean of the four samples was used as a study-wide long-read WGS confirmation rate, weighting each sample’s confirmation rate based on the square root of its long-read WGS sequencing depth. (**c**) Confirmation rates stratified by SV class, size, and AF.

**Extended Data Figure 4.**
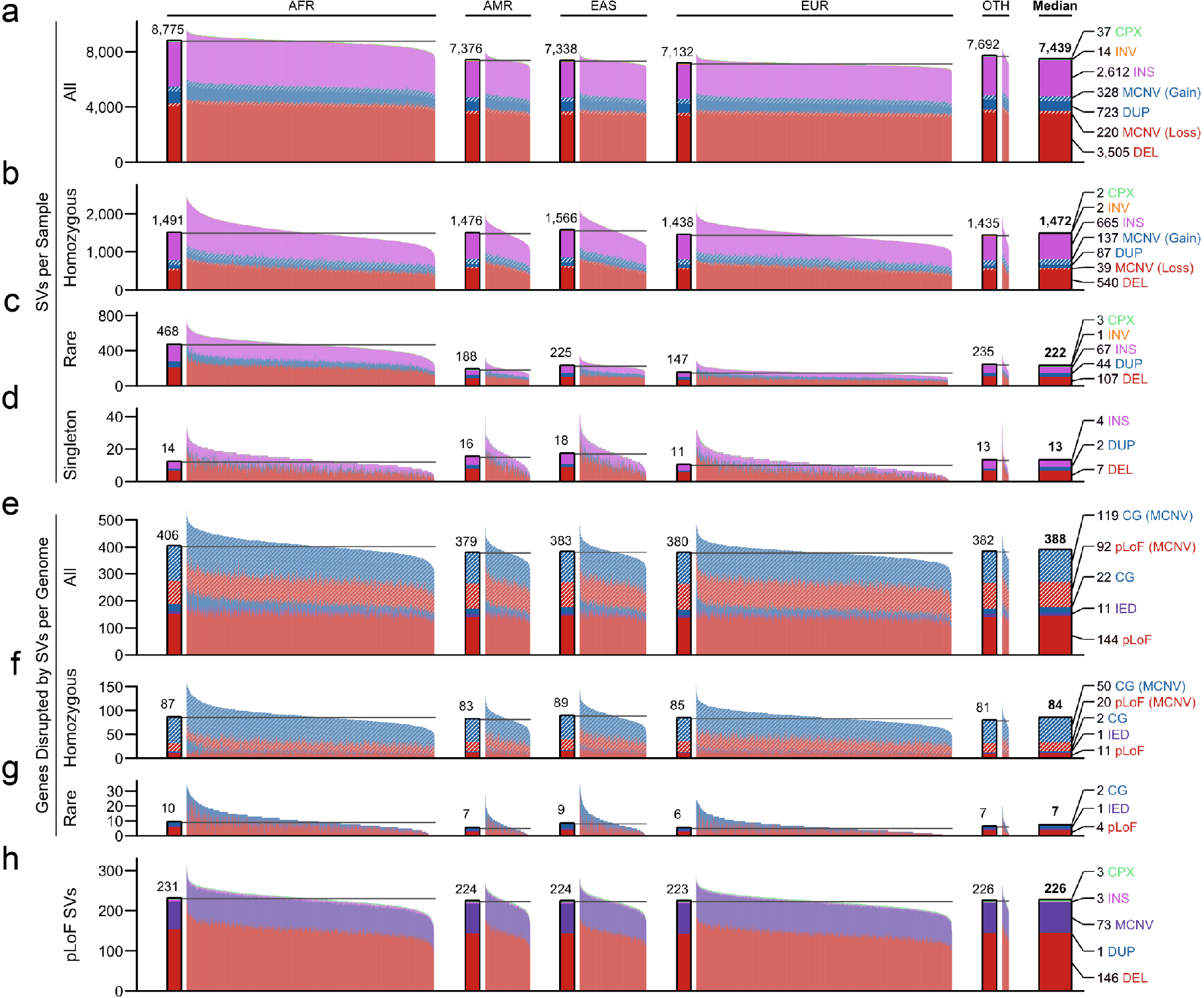
SVs contribute a substantial burden of rare, homozygous, and coding mutations per genome. (**a**-**d**) Counts of SVs per genome across a variety of parameters, corresponding to median counts of (a) total SVs, (b) homozygous SVs, (c) rare SVs, and (d) singleton SVs. Samples are grouped by population and colored by SV types. The solid bar to the left of each population indicates the population median. (**e**-**g**) Median counts of genes disrupted by SVs per genome when considering (e) all SVs (including MCNVs), (f) homozygous SVs (including MCNVs), and (g) rare SVs. Colors correspond to predicted functional consequence. (**h**) Counts of pLoF SVs per genome. For certain categories, such as genes disrupted by rare SVs per genome, a subset of samples (<5%) were enriched above the population average, as expected for individuals carrying large, rare CNVs predicted to cause the disruption of dozens or hundreds of genes (see **Extended Data Figure 1**); for the purposes of visualization, the y-axis for all panels presented here has been restricted to a maximum of three interquartile ranges above the third quartile across all samples for each category.

**Extended Data Figure 5.**
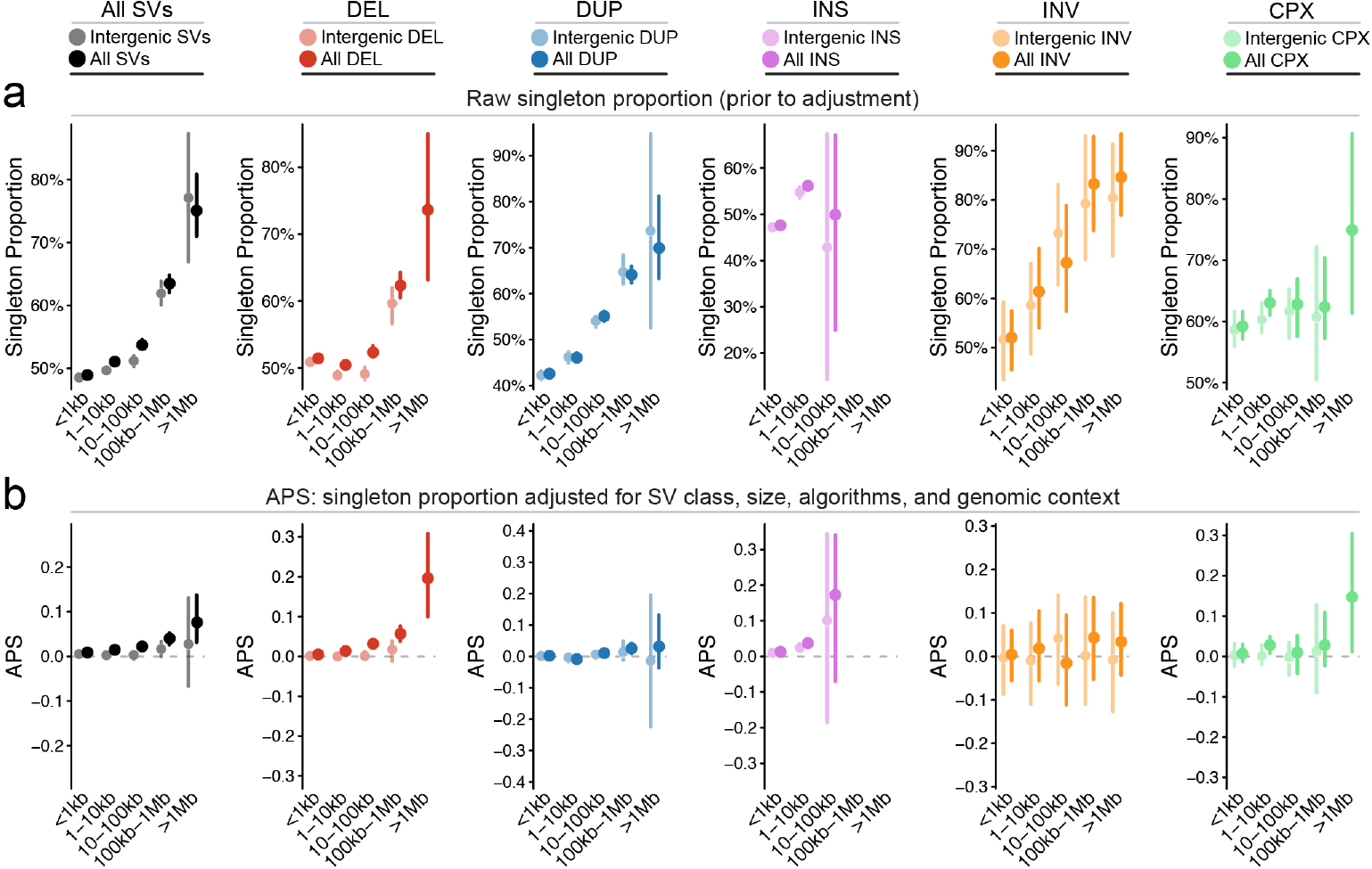
Rearrangement size is a primary determinant of allele frequency for most classes of SVs. (**a**) Proportion of singleton SVs in five SV size bins for each class of biallelic SVs considered in this study. Intergenic SVs (light colors) exhibited reduced singleton proportions when compared to all SVs (dark colors) of the same size and class. Bars reflect 95% confidence intervals from 100-fold bootstrapping. (**b**) To account for the strong dependency of singleton proportion on SV size and class, we developed a metric dubbed the “<u>A</u>djusted <u>P</u>roportion of <u>S</u>ingletons” (APS), which normalizes all values to zero to permit comparisons of the frequency spectra across SV classes (see **Supplementary Figure 14**). Shown here is the same data from (a) transformed onto the APS scale, which shows effectively no dependency on SV size for intergenic SVs. Bars reflect 95% confidence intervals from 100-fold bootstrapping. Residual deviation from APS=0 is maintained when considering all SVs, due to APS being intentionally calibrated to intergenic SVs as a proxy for neutral variation. Since larger SVs are more likely to be gene-disruptive, they upwardly bias the APS point estimates due to residual negative selection not captured by SV size alone.

**Extended Data Figure 6.**
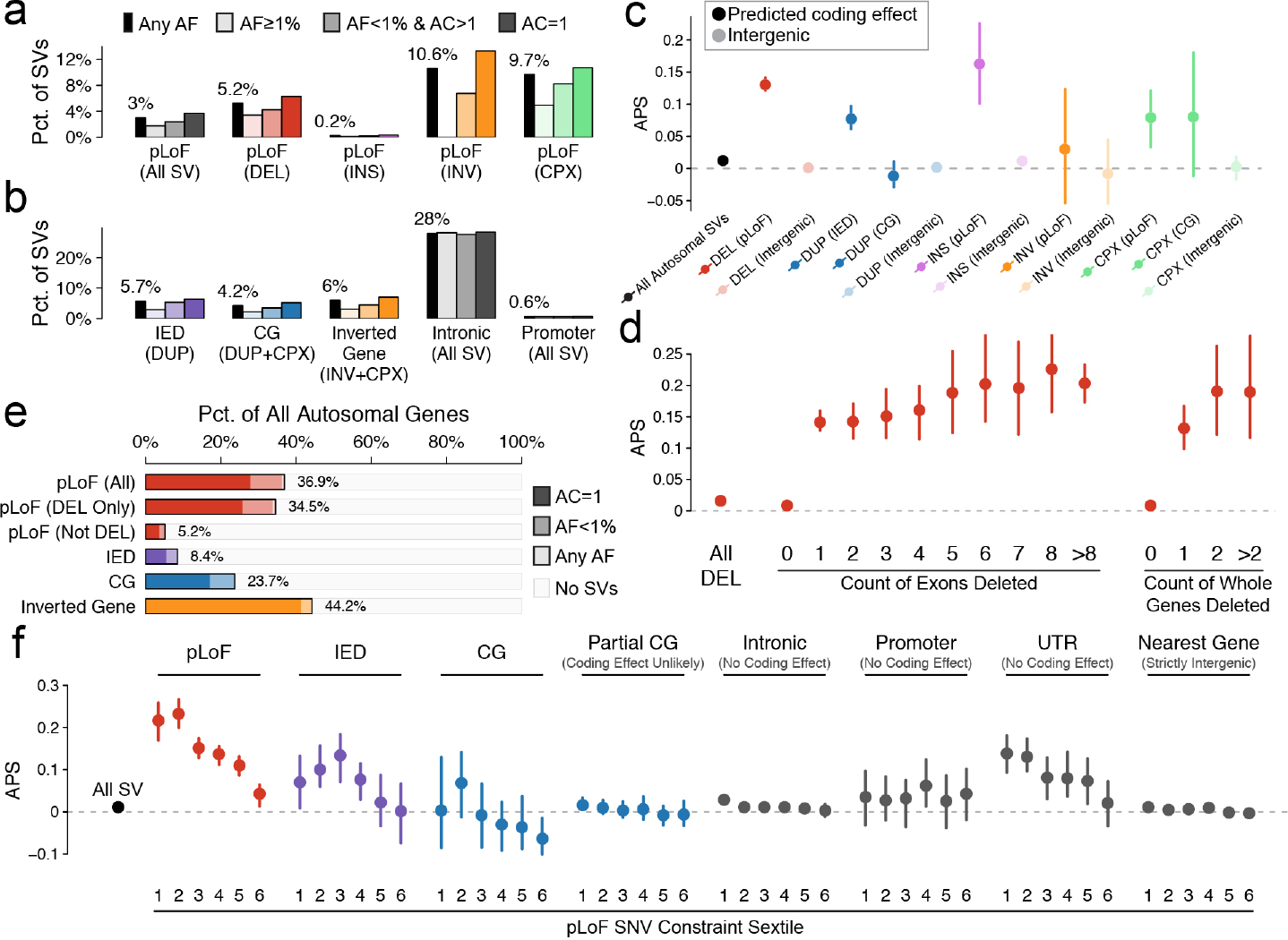
Most SVs within genes appear under negative selection. (**a**) Enrichments for pLoF consequences among rare and singleton SVs across SV classes. (**b**) Enrichments for non-pLoF functional consequences among rare and singleton SVs across SV classes. (**c**) Proportion of singletons, represented by our APS metric, across SV types and functional consequences. For panels (c), (d), and (f), bars represent 95% confidence intervals from 100-fold bootstrapping. (**d**) APS among deletions relative to count of exons and whole-genes deleted. For panels (d) and (f), deletions in highly repetitive or low-complexity sequence (≥30% coverage by annotated segmental duplications or simple repeats) were excluded. (**e**) Fractions of all autosomal protein-coding genes with at least one SV across a variety of functional consequences. (**f**) Relationship of APS and constraint against pLoF SNVs.^22^ For this analysis, intronic, promoter, and UTR SVs were required to have precise breakpoints (i.e., have “split-read” support) to protect against any cryptic overlap with coding sequence unable to be annotated due to imprecise breakpoints.

**Extended Data Figure 7.**
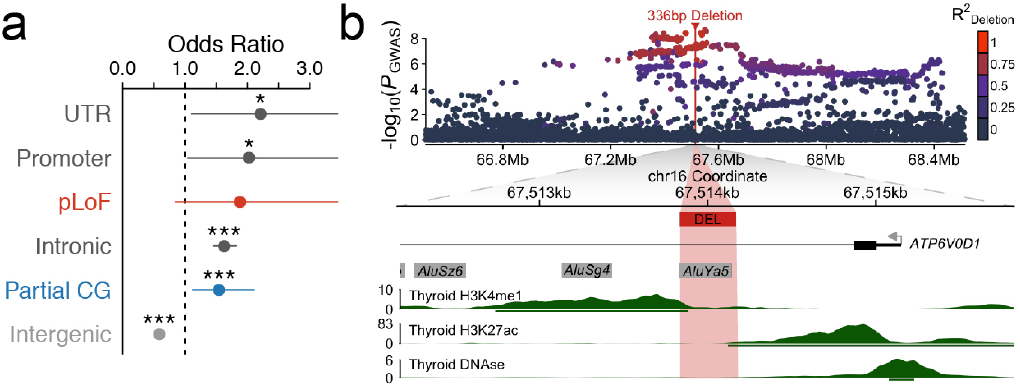
gnomAD-SV can augment disease association studies. (**a**) Functional enrichments of 2,307 common SVs in strong LD (R^2^≥0.8) with an SNV associated with a trait or disease in the GWAS Catalog or the UK BioBank.^45,46^ Points represent odds ratios of SVs being in strong LD with at least one GWAS-significant SNV among all SVs in strong LD with at least one SNV (total N=15,634 SVs). Single and triple asterisks correspond to nominal (P<0.05) and Bonferroni-corrected significance thresholds from a two-sided Fisher’s Exact Test, respectively. Bars represent 95% confidence intervals. (**b**) Example locus on 16q22.1, where we identified a 336bp deletion in strong LD with SNVs significantly associated with hypothyroidism in the UK BioBank.^46^ The top panel depicts the GWAS signal among genotyped SNVs in the UK BioBank, colored by strength of LD with the 336bp deletion identified in gnomAD-SV. The bottom panel depicts the local genomic context of this deletion, which overlaps an annotated intronic Alu element near (<1kb) the first exon of a highly constrained, thyroid-expressed gene, ATP6V0D1. The deletion lies amidst histone mark peaks commonly found at active enhancers (H3K27ac & H3K4me1) based on publicly available chromatin data from adult thyroid samples, a phenotype-relevant tissue.^70^ Human Alu elements are known to frequently act as enhancers,^71^ and the sentinel hypothyroidism SNV from the UK BioBank GWAS is a significant expression-modifying variant (i.e., eQTL) for ATP6V0D1 and other nearby genes across many tissues, implying that the hypothyroidism risk haplotype modifies expression of ATP6V0D1 and/or other genes, potentially through the deletion of an intronic enhancer.^22,72^

**Extended Data Figure 8.**
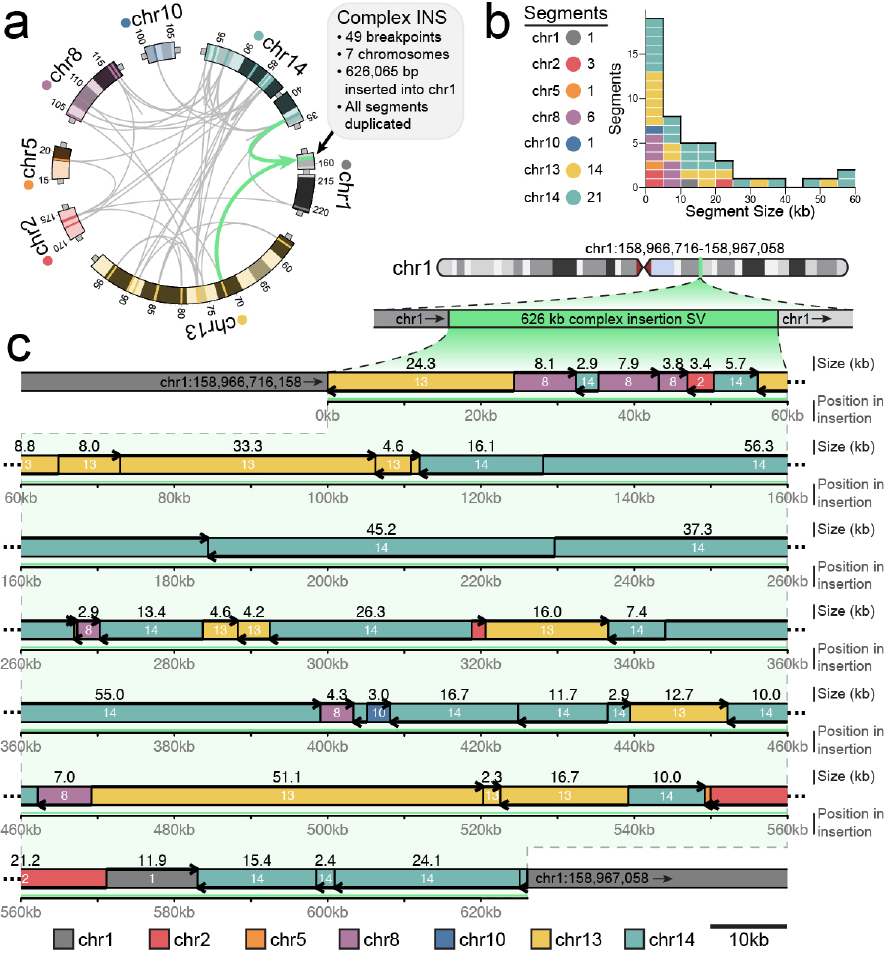
An extremely complex SV involving 49 breakpoints and seven chromosomes. A highly complex insertion rearrangement from gnomAD-SV where 47 segments from six different chromosomes were duplicated and inserted into a single locus on chromosome 1, forming a 626,065bp stretch of contiguous inserted sequence composed of shattered fragments. Given the involvement of multiple chromosomes, the signature of localized shattering, and the clustered breakpoints, we note that this rearrangement has several hallmarks of germline chromothripsis,^14,55^ which has been observed in healthy adults previously, albeit rarely.^55^ However, unlike previous reports of germline chromothripsis, there are no apparent whole-chromosome translocations, and all segments were duplicated before being inserted in a compound manner into chromosome 1, potentially suggesting a replication-based repair mechanism. The exact origin of this rearrangement is unclear. (**a**) Circos representation of all 49 breakpoints and seven chromosomes involved in this SV. Teal arrows indication insertion point into chromosome (**b**) The median segment size was 8.4kb. (**c**) Linear representation of the rearranged inserted sequence. Colors correspond to chromosome of origin, and arrows indicate strandedness of inserted sequence, relative to the GRCh37 reference.

## GROUP AUTHORS

### Genome Aggregation Database Production Team

Jessica Alföldi^1,2^, Irina M. Armean^3,1,2^, Eric Banks^4^, Louis Bergelson^4^, Kristian Cibulskis^4^, Ryan L Collins^1,5,6^, Kristen M. Connolly^7^, Miguel Covarrubias^4^, Beryl Cummings^1,2,8^, Mark J. Daly^1,2,9^, Stacey Donnelly^1^, Yossi Farjoun^4^, Steven Ferriera^10^, Laurent Francioli^1,2^, Stacey Gabriel^10^, Laura D. Gauthier^4^, Jeff Gentry^4^, Namrata Gupta^10,1^, Thibault Jeandet^4^, Diane Kaplan^4^, Konrad J. Karczewski^1,2^, Kristen M. Laricchia^1,2^, Christopher Llanwarne^4^, Eric V. Minikel^1^, Ruchi Munshi^4^, Benjamin M Neale^1,2^, Sam Novod^4^, Anne H. O’Donnell-Luria^1,11,12^, Nikelle Petrillo^4^, Timothy Poterba^9,2,1^, David Roazen^4^, Valentin Ruano-Rubio^4^, Andrea Saltzman^1^, Kaitlin E. Samocha^13^, Molly Schleicher^1^, Cotton Seed^9,2^, Matthew Solomonson^1,2^, Jose Soto^4^, Grace Tiao^1,2^, Kathleen Tibbetts^4^, Charlotte Tolonen^4^, Christopher Vittal^9,2^, Gordon Wade^4^, Arcturus Wang^9,2,1^, Qingbo Wang^1,2,6^, James S Ware^14,15,1^, Nicholas A Watts^1,2^, Ben Weisburd^4^, Nicola Whiffin^14,15,1^

1. Program in Medical and Population Genetics, Broad Institute of MIT and Harvard, Cambridge, Massachusetts 02142, USA

2. Analytic and Translational Genetics Unit, Massachusetts General Hospital, Boston, Massachusetts 02114, USA

3. European Molecular Biology Laboratory, European Bioinformatics Institute, Wellcome Genome Campus, Hinxton, Cambridge, CB10 1SD, United Kingdom

4. Data Sciences Platform, Broad Institute of MIT and Harvard, Cambridge, Massachusetts 02142, USA

5. Center for Genomic Medicine, Massachusetts General Hospital, Boston, MA 02114, USA

6. Program in Bioinformatics and Integrative Genomics, Harvard Medical School, Boston, MA 02115, USA

7. Genomics Platform, Broad Institute of MIT and Harvard, Cambridge, Massachusetts 02142, USA

8. Program in Biological and Biomedical Sciences, Harvard Medical School, Boston, MA, 02115, USA

9. Stanley Center for Psychiatric Research, Broad Institute of MIT and Harvard, Cambridge, Massachusetts 02142, USA

10. Broad Genomics, Broad Institute of MIT and Harvard, Cambridge, Massachusetts 02142, USA

11. Division of Genetics and Genomics, Boston Children’s Hospital, Boston, Massachusetts 02115, USA

12. Department of Pediatrics, Harvard Medical School, Boston, Massachusetts 02115, USA

13. Wellcome Sanger Institute, Wellcome Genome Campus, Hinxton, Cambridge CB10 1SA, UK

14. National Heart & Lung Institute and MRC London Institute of Medical Sciences, Imperial College London, London UK

15. Cardiovascular Research Centre, Royal Brompton & Harefield Hospitals NHS Trust, London UK

### Genome Aggregation Database Consortium

Carlos A Aguilar Salinas^1^, Tariq Ahmad^2^, Christine M. Albert^3,4^, Diego Ardissino^5^, Gil Atzmon^6,7^, John Barnard^8^, Laurent Beaugerie^9^, Emelia J. Benjamin^10,11,12^, Michael Boehnke^13^, Lori L. Bonnycastle^14^, Erwin P. Bottinger^15^, Donald W Bowden^16,17,18^, Matthew J Bown^19,20^, John C Chambers^21,22,23^, Juliana C. Chan^24^, Daniel Chasman^3,25^, Judy Cho^15^, Mina K. Chung^26^, Bruce Cohen^27,25^, Adolfo Correa^28^, Dana Dabelea^29^, Mark J. Daly^30,31,32^, Dawood Darbar^33^, Ravindranath Duggirala^34^, Josée Dupuis^35,36^, Patrick T. Ellinor^30,37^, Roberto Elosua^38,39,40^, Jeanette Erdmann^41,42,43^, Tõnu Esko^30,44^, Martti Färkkilä^45^, Jose Florez^46^, Andre Franke^47^, Gad Getz^48,49,25^, Benjamin Glaser^50^, Stephen J. Glatt^51^, David Goldstein^52,53^, Clicerio Gonzalez^54^, Leif Groop^55,56^, Christopher Haiman^57^, Craig Hanis^58^, Matthew Harms^59,60^, Mikko Hiltunen^61^, Matti M. Holi^62^, Christina M. Hultman^63,64^, Mikko Kallela^65^, Jaakko Kaprio^56,66^, Sekar Kathiresan^67,68,25^, Bong-Jo Kim^69^, Young Jin Kim^69^, George Kirov^70^, Jaspal Kooner^23,22,71^, Seppo Koskinen^72^, Harlan M. Krumholz^73^, Subra Kugathasan^74^, Soo Heon Kwak^75^, Markku Laakso^76,77^, Terho Lehtimäki^78^, Ruth J.F. Loos^15,79^, Steven A. Lubitz^30,37^, Ronald C.W. Ma^24,80,81^, Daniel G. MacArthur^31,30^, Jaume Marrugat^82,39^, Kari M. Mattila^78^, Steven McCarroll^32,83^, Mark I McCarthy^84,85,86^, Dermot McGovern^87^, Ruth McPherson^88^, James B. Meigs^89,25,90^, Olle Melander^91^, Andres Metspalu^44^, Benjamin M Neale^30,31^, Peter M Nilsson^92^, Michael C O’Donovan^70^, Dost Ongur^27,25^, Lorena Orozco^93^, Michael J Owen^70^, Colin N.A. Palmer^94^, Aarno Palotie^56,32,31^, Kyong Soo Park^75,95^, Carlos Pato^96^, Ann E. Pulver^97^, Nazneen Rahman^98^, Anne M. Remes^99^, John D. Rioux^100,101^, Samuli Ripatti^56,66,102^, Dan M. Roden^103,104^, Danish Saleheen^105,106,107^, Veikko Salomaa^108^, Nilesh J. Samani^19,20^, Jeremiah Scharf^30,32,67^, Heribert Schunkert^109,110^, Moore B. Shoemaker^111^, Pamela Sklar^*112,113,114^, Hilkka Soininen^115^, Harry Sokol^9^, Tim Spector^116^, Patrick F. Sullivan^63,117^, Jaana Suvisaari^108^, E Shyong Tai^118,119,120^, Yik Ying Teo^118,121,122^, Tuomi Tiinamaija^56,123,124^, Ming Tsuang^125,126^, Dan Turner^127^, Teresa Tusie-Luna^128,129^, Erkki Vartiainen^66^, James S Ware^130,131,30^, Hugh Watkins^132^, Rinse K Weersma^133^, Maija Wessman^123,56^, James G. Wilson^134^, Ramnik J. Xavier^135,136^

1. Unidad de Investigacion de Enfermedades Metabolicas. Instituto Nacional de Ciencias Medicas y Nutricion. Mexico City

2. Peninsula College of Medicine and Dentistry, Exeter, UK

3. Division of Preventive Medicine, Brigham and Women’s Hospital, Boston, Massachusetts, USA.

4. Division of Cardiovascular Medicine, Brigham and Women’s Hospital and Harvard Medical School, Boston, Massachusetts, USA.

5. Department of Cardiology, University Hospital, 43100 Parma, Italy

6. Department of Biology, Faculty of Natural Sciences, University of Haifa, Haifa, Israel

7. Departments of Medicine and Genetics, Albert Einstein College of Medicine, Bronx, NY, USA, 10461

8. Department of Quantitative Health Sciences, Lerner Research Institute, Cleveland Clinic, Cleveland, OH 44122, USA

9. Sorbonne Université, APHP, Gastroenterology Department, Saint Antoine Hospital, Paris, France

10. NHLBI and Boston University’s Framingham Heart Study, Framingham, Massachusetts, USA.

11. Department of Medicine, Boston University School of Medicine, Boston, Massachusetts, USA.

12. Department of Epidemiology, Boston University School of Public Health, Boston, Massachusetts, USA.

13. Department of Biostatistics and Center for Statistical Genetics, University of Michigan, Ann Arbor, Michigan 48109

14. National Human Genome Research Institute, National Institutes of Health, Bethesda, MD, USA

15. The Charles Bronfman Institute for Personalized Medicine, Icahn School of Medicine at Mount Sinai, New York, NY

16. Department of Biochemistry, Wake Forest School of Medicine, Winston-Salem, NC, USA

17. Center for Genomics and Personalized Medicine Research, Wake Forest School of Medicine, Winston-Salem, NC, USA

18. Center for Diabetes Research, Wake Forest School of Medicine, Winston-Salem, NC, USA

19. Department of Cardiovascular Sciences, University of Leicester, Leicester, UK

20. NIHR Leicester Biomedical Research Centre, Glenfield Hospital, Leicester, UK

21. Department of Epidemiology and Biostatistics, Imperial College London, London, UK

22. Department of Cardiology, Ealing Hospital NHS Trust, Southall, UK

23. Imperial College Healthcare NHS Trust, Imperial College London, London, UK

24. Department of Medicine and Therapeutics, The Chinese University of Hong Kong, Hong Kong, China.

25. Department of Medicine, Harvard Medical School, Boston, MA

26. Departments of Cardiovascular Medicine, Cellular and Molecular Medicine, Molecular Cardiology, and Quantitative Health Sciences, Cleveland Clinic, Cleveland, Ohio, USA.

27. McLean Hospital, Belmont, MA

28. Department of Medicine, University of Mississippi Medical Center, Jackson, Mississippi, USA

29. Department of Epidemiology, Colorado School of Public Health, Aurora, Colorado, USA.

30. Program in Medical and Population Genetics, Broad Institute of MIT and Harvard, Cambridge, MA, USA

31. Analytic and Translational Genetics Unit, Massachusetts General Hospital, Boston, Massachusetts 02114, USA

32. Stanley Center for Psychiatric Research, Broad Institute of MIT and Harvard, Cambridge, MA, USA

33. Department of Medicine and Pharmacology, University of Illinois at Chicago

34. Department of Genetics, Texas Biomedical Research Institute, San Antonio, TX, USA

35. Department of Biostatistics, Boston University School of Public Health, Boston, MA 02118, USA

36. National Heart, Lung, and Blood Institute’s Framingham Heart Study, Framingham, MA 01702, USA

37. Cardiac Arrhythmia Service and Cardiovascular Research Center, Massachusetts General Hospital, Boston, MA

38. Cardiovascular Epidemiology and Genetics, Hospital del Mar Medical Research Institute (IMIM). Barcelona, Catalonia, Spain

39. CIBER CV, Barcelona, Catalonia, Spain

40. Department of Medicine, Medical School, University of Vic-Central University of Catalonia. Vic, Catalonia, Spain

41. Institute for Cardiogenetics, University of Lübeck, Lübeck, Germany

42.1. DZHK (German Research Centre for Cardiovascular Research), partner site Hamburg/Lübeck/Kiel, 23562 Lübeck, Germany

43. University Heart Center Lübeck, 23562 Lübeck, Germany

44. Estonian Genome Center, Institute of Genomics, University of Tartu, Tartu, Estonia

45. Helsinki University and Helsinki University Hospital, Clinic of Gastroenterology, Helsinki, Finland.

46. Diabetes Unit and Center for Genomic Medicine, Massachusetts General Hospital; Programs in Metabolism and Medical & Population Genetics, Broad Institute; Department of Medicine, Harvard Medical School

47. Institute of Clinical Molecular Biology (IKMB), Christian-Albrechts-University of Kiel, Kiel, Germany

48. Bioinformatics Program, MGH Cancer Center and Department of Pathology

49. Cancer Genome Computational Analysis, Broad Institute.

50. Endocrinology and Metabolism Department, Hadassah-Hebrew University Medical Center, Jerusalem, Israel

51. Department of Psychiatry and Behavioral Sciences; SUNY Upstate Medical University

52. Institute for Genomic Medicine, Columbia University Medical Center, Hammer Health Sciences, 1408, 701 West 168th Street, New York, New York 10032, USA.

53. Department of Genetics & Development, Columbia University Medical Center, Hammer Health Sciences, 1602, 701 West 168th Street, New York, New York 10032, USA.

54. Centro de Investigacion en Salud Poblacional. Instituto Nacional de Salud Publica MEXICO

55. Lund University, Sweden

56. Institute for Molecular Medicine Finland (FIMM), HiLIFE, University of Helsinki, Helsinki, Finland

57. Lund University Diabetes Centre

58. Human Genetics Center, University of Texas Health Science Center at Houston, Houston, TX 77030

59. Department of Neurology, Columbia University

60. Institute of Genomic Medicine, Columbia University

61. Institute of Biomedicine, University of Eastern Finland, Kuopio, Finland

62. Department of Psychiatry, PL 320, Helsinki University Central Hospital, Lapinlahdentie, 00 180 Helsinki, Finland

63. Department of Medical Epidemiology and Biostatistics, Karolinska Institutet, Stockholm, Sweden

64. Icahn School of Medicine at Mount Sinai, New York, NY, USA

65. Department of Neurology, Helsinki University Central Hospital, Helsinki, Finland.

66. Department of Public Health, Faculty of Medicine, University of Helsinki, Finland

67. Center for Genomic Medicine, Massachusetts General Hospital, Boston, Massachusetts 02114, USA

68. Cardiovascular Disease Initiative and Program in Medical and Population Genetics, Broad Institute of MIT and Harvard, Cambridge, Massachusetts 02142, USA

69. Center for Genome Science, Korea National Institute of Health, Chungcheongbuk-do, Republic of Korea.

70. MRC Centre for Neuropsychiatric Genetics & Genomics, Cardiff University School of Medicine, Hadyn Ellis Building, Maindy Road, Cardiff CF24 4HQ

71. National Heart and Lung Institute, Cardiovascular Sciences, Hammersmith Campus, Imperial College London, London, UK.

72. Department of Health, THL-National Institute for Health and Welfare, 00271 Helsinki, Finland.

73. Section of Cardiovascular Medicine, Department of Internal Medicine, Yale School of Medicine, New Haven, Connecticut Center for Outcomes Research and Evaluation, Yale-New Haven Hospital, New Haven, Connecticut.

74. Division of Pediatric Gastroenterology, Emory University School of Medicine, Atlanta, Georgia, USA.

75. Department of Internal Medicine, Seoul National University Hospital, Seoul, Republic of Korea

76. The University of Eastern Finland, Institute of Clinical Medicine, Kuopio, Finland

77. Kuopio University Hospital, Kuopio, Finland

78. Department of Clinical Chemistry, Fimlab Laboratories and Finnish Cardiovascular Research Center-Tampere, Faculty of Medicine and Health Technology, Tampere University, Finland

79. The Mindich Child Health and Development Institute, Icahn School of Medicine at Mount Sinai, New York, NY

80. Li Ka Shing Institute of Health Sciences, The Chinese University of Hong Kong, Hong Kong, China.

81. Hong Kong Institute of Diabetes and Obesity, The Chinese University of Hong Kong, Hong Kong, China.

82. Cardiovascular Research REGICOR Group, Hospital del Mar Medical Research Institute (IMIM). Barcelona, Catalonia.

83. Department of Genetics, Harvard Medical School, Boston, MA, USA

84. Oxford Centre for Diabetes, Endocrinology and Metabolism, University of Oxford, Churchill Hospital, Old Road, Headington, Oxford, OX3 7LJ UK

85. Wellcome Centre for Human Genetics, University of Oxford, Roosevelt Drive, Oxford OX3 7BN, UK

86. Oxford NIHR Biomedical Research Centre, Oxford University Hospitals NHS Foundation Trust, John Radcliffe Hospital, Oxford OX3 9DU, UK

87. F Widjaja Foundation Inflammatory Bowel and Immunobiology Research Institute, Cedars-Sinai Medical Center, Los Angeles, CA, USA.

88. Atherogenomics Laboratory, University of Ottawa Heart Institute, Ottawa, Canada

89. Division of General Internal Medicine, Massachusetts General Hospital, Boston, MA, 02114

90. Program in Population and Medical Genetics, Broad Institute, Cambridge, MA

91. Department of Clinical Sciences, University Hospital Malmo Clinical Research Center, Lund University, Malmo, Sweden.

92. Lund University, Dept. Clinical Sciences, Skane University Hospital, Malmo, Sweden

93. Instituto Nacional de Medicina Genómica (INMEGEN), Mexico City, 14610, Mexico

94. Medical Research Institute, Ninewells Hospital and Medical School, University of Dundee, Dundee, UK.

95. Department of Molecular Medicine and Biopharmaceutical Sciences, Graduate School of Convergence Science and Technology, Seoul National University, Seoul, Republic of Korea

96. Department of Psychiatry, Keck School of Medicine at the University of Southern California, Los Angeles, California, USA.

97. Department of Psychiatry and Behavioral Sciences, Johns Hopkins University School of Medicine, Baltimore, Maryland, USA

98. Division of Genetics and Epidemiology, Institute of Cancer Research, London SM2 5NG

99. Medical Research Center, Oulu University Hospital, Oulu, Finland and Research Unit of Clinical Neuroscience, Neurology, University of Oulu, Oulu, Finland.

100. Research Center, Montreal Heart Institute, Montreal, Quebec, Canada, H1T 1C8

101. Department of Medicine, Faculty of Medicine, Université de Montréal, Québec, Canada

102. Broad Institute of MIT and Harvard, Cambridge MA, USA

103. Department of Biomedical Informatics, Vanderbilt University Medical Center, Nashville, Tennessee, USA.

104. Department of Medicine, Vanderbilt University Medical Center, Nashville, Tennessee, USA.

105. Department of Biostatistics and Epidemiology, Perelman School of Medicine at the University of Pennsylvania, Philadelphia, PA, USA

106. Department of Medicine, Perelman School of Medicine at the University of Pennsylvania, Philadelphia, PA, USA

107. Center for Non-Communicable Diseases, Karachi, Pakistan

108. National Institute for Health and Welfare, Helsinki, Finland

109. Deutsches Herzzentrum München, Germany

110. Technische Universität München

111. Division of Cardiovascular Medicine, Nashville VA Medical Center and Vanderbilt University, School of Medicine, Nashville, TN 37232-8802, USA

112. Department of Psychiatry, Icahn School of Medicine at Mount Sinai, New York, NY, USA

113. Department of Genetics and Genomic Sciences, Icahn School of Medicine at Mount Sinai, New York, NY, USA

114. Institute for Genomics and Multiscale Biology, Icahn School of Medicine at Mount Sinai, New York, NY, USA

115. Institute of Clinical Medicine, neurology, University of Eastern Finland, Kuopio, Finland

116. Department of Twin Research and Genetic Epidemiology, King’s College London, London UK

117. Departments of Genetics and Psychiatry, University of North Carolina, Chapel Hill, NC, USA

118. Saw Swee Hock School of Public Health, National University of Singapore, National University Health System, Singapore

119. Department of Medicine, Yong Loo Lin School of Medicine, National University of Singapore, Singapore

120. Duke-NUS Graduate Medical School, Singapore

121. Life Sciences Institute, National University of Singapore, Singapore

122. Department of Statistics and Applied Probability, National University of Singapore, Singapore

123. Folkhälsan Institute of Genetics, Folkhälsan Research Center, Helsinki, Finland

124. HUCH Abdominal Center, Helsinki University Hospital, Helsinki, Finland

125. Center for Behavioral Genomics, Department of Psychiatry, University of California, San Diego

126. Institute of Genomic Medicine, University of California, San Diego

127. Juliet Keidan Institute of Pediatric Gastroenterology, Shaare Zedek Medical Center, The Hebrew University of Jerusalem, Israel

128. Instituto de Investigaciones Biomédicas UNAM Mexico City

129. Instituto Nacional de Ciencias Médicas y Nutrición Salvador Zubirán Mexico City

130. National Heart & Lung Institute & MRC London Institute of Medical Sciences, Imperial College London, London UK

131. Cardiovascular Research Centre, Royal Brompton & Harefield Hospitals NHS Trust, London UK

132. Radcliffe Department of Medicine, University of Oxford, Oxford UK

133. Department of Gastroenterology and Hepatology, University of Groningen and University Medical Center Groningen, Groningen, the Netherlands

134. Department of Physiology and Biophysics, University of Mississippi Medical Center, Jackson, MS 39216, USA

135. Program in Infectious Disease and Microbiome, Broad Institute of MIT and Harvard, Cambridge, MA, USA

136. Center for Computational and Integrative Biology, Massachusetts General Hospital

