## Supplementary Information for "An open resource of structural variation for medical and population genetics"

#### Table of Contents

**Supplementary Figures**

1. Sample QC & exclusion prior to SV discovery 4
2. Distribution of sample ages 6
3. Overview of gnomAD-SV discovery pipeline 7
4. Whole-genome dosage bias quantification 8
5. Sample batching strategy 10
6. Hardy-Weinberg Equilibria of SVs across major continental populations 12
7. Patterns of linkage disequilibrium between SVs and SNVs/indels 13
8. Site-level comparison of SVs to the 1000 Genomes Project 14
9. Allele frequency comparisons to SVs from the 1000 Genomes Project 15
10. Breakpoint accuracy estimates from long-read WGS 16
11. Distribution of variant quality scores 17
12. Validation of variant quality score calibration 18
13. Principal components analysis of common SVs 20
14. Development of the adjusted proportion of singletons (APS) metric 21
15. Comparison of allele frequencies per SV class by genic context 22
16. Genomic patterns and correlates of SV density 23
17. Summary of SV annotations in coding sequences 24
18. Raw singleton proportions for selected findings 25
19. Comparisons of SV depletion vs SNV pLoF and missense constraint 26
20. Evidence for genomic disorder CNV carriers in gnomAD-SV 27
21. Properties of genomic disorders evaluated in gnomAD-SV 29
22. Homozygous pLoF SVs 30
23. Overview of *post hoc* filtering & final callsets 31
24. Evaluation of callset filtering on key results 32

**Supplementary Tables**

1. Sample QC thresholds & filtering 33
2. Sample overlap with gnomAD SNV/indel analyses 34
3. SV callset summary 35

4. SV callset benchmarking 36
5. Common SVs in strong LD with SNV/indels 37
6. Carrier frequencies of genomic disorder CNVs in gnomAD-SV 38

**Supplementary Notes**

1. Technical benchmarking and quality assessment of the gnomAD-SV callset 39

**Methods**

WGS data aggregation 40
Computational platform 40
SV discovery 40
*Module 00: Preprocessing* 41
*Module 01: Clustering* 45
*Module 02: Evidence Collection* 45
*Module 03: Variant Filtering* 46
*Module 04: Genotyping* 47
*Module 05: Batch Integration* 48
*Module 06: VCF Refinement* 50
*Module 07: Gene annotation* 51
Sample and variant QC after SV discovery 53
*Optimizing per-sample genotype quality filters* 53
*Outlier sample exclusion* 55
*Assessment of batch effects* 55
*Assignment of final VCF filter labels* 56
*Variant quality score recalibration* 57
*Population assignments* 57
*Relatedness inference* 58
*Final variant modifications and callset curation* 58
Callset benchmarking 61
*Assessment of Mendelian violation rate from parent-child trios* 61
*Comparison to chromosomal microarray data on matched samples* 62
*Analysis of Hardy-Weinberg equilibrium across populations* 62
*Comparison to long-read WGS data on matched samples* 63
*Comparison to SVs from the 1000 Genomes Project* 63
*Cross-population genotype analysis for doubleton SVs* 64
*Analysis of linkage disequilibrium between SNVs/indels and SVs* 64
Calculating adjusted proportion of singletons (APS) 65
Chromosome-level analyses of SV density 65
Annotated repetitive element correlations 66
Estimating SV mutation rates 66
Comparisons of SVs to metrics of genic constraint against point mutations 67
Estimating noncoding selection against cis-regulatory annotation classes 67
*Definition of noncoding CNVs* 68
*Curation of functional annotation classes* 68
*Calculation of APS for noncoding CNVs overlapping functional annotation classes* 69
Intersection of gnomAD-SV and published genome-wide association studies 70
Analysis of genomic disorder loci 71

gnomAD-SV callset downsampling analyses 72
Predicting the rate of clinically reportable incidental findings from SVs 72
Evaluation of callset filtering on key results 72
Compliance with ethical regulations 72
Data availability 73
Code availability 73

**Supplementary References** 74

### SUPPLEMENTARY FIGURES

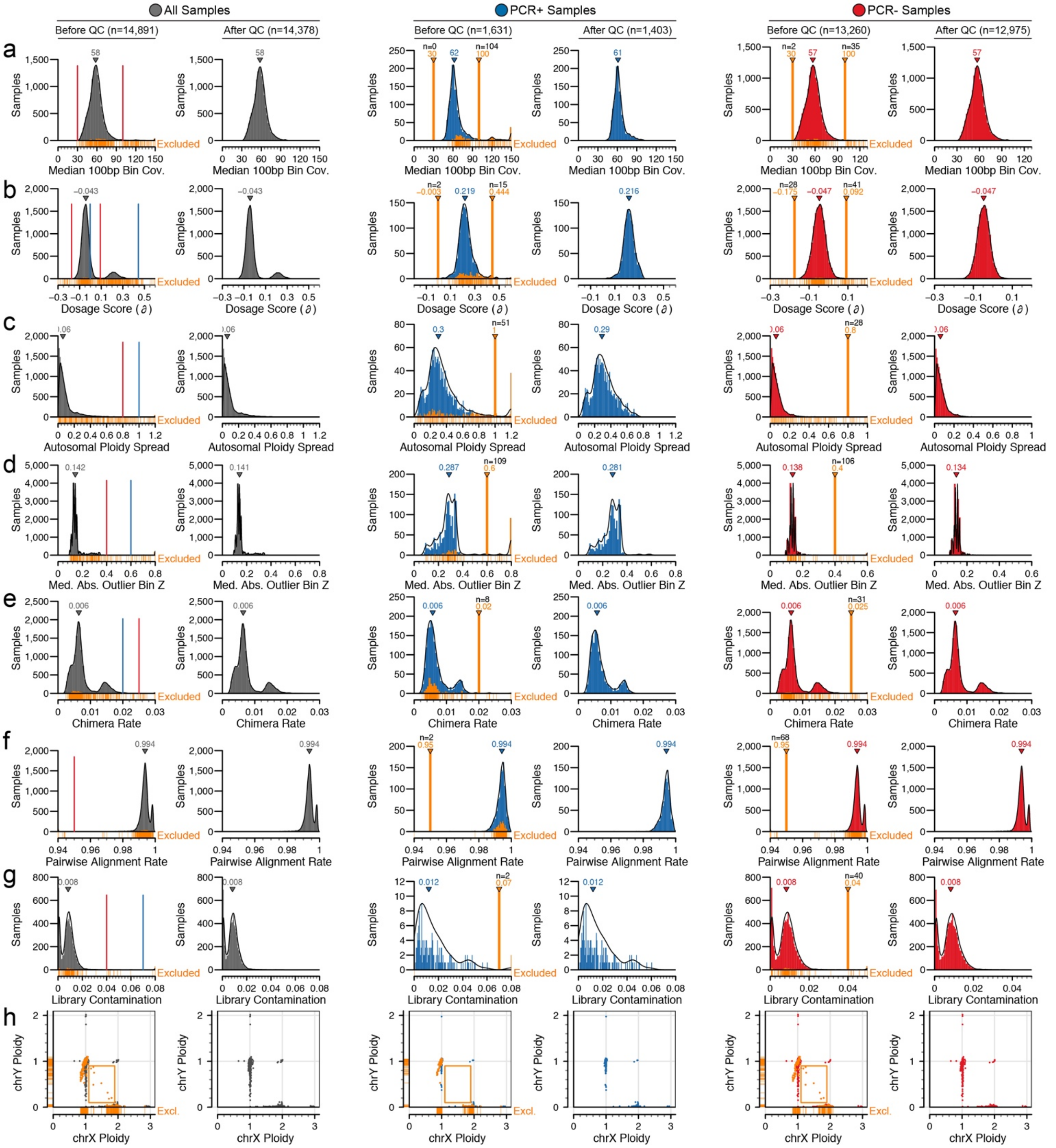


**Supplementary Figure 1 | Sample QC & exclusion prior to SV discovery.** We performed quality control (QC) on all 14,891 samples in the gnomAD-SV cohort prior to structural variant (SV) discovery, excluding a total of 513 samples (3.45% of the cohort) that failed to meet baseline sample or WGS quality thresholds for at least one of 10 features considered. A description of this filtering process and the definitions of the 10 filtered features is provided in **Methods**. The distributions, filter thresholds, and sample exclusion statistics for 8/10 of these features are depicted here. Rows correspond to one filtered feature. Pairs of columns (those with distributions of the same color) correspond to all samples (left), PCR+ samples (center), and PCR- samples (right). Within each pair of columns, the left and right panels represent the distribution of the feature before and after sample exclusion, respectively. Orange lines indicate filter exclusion thresholds, and the orange portions of each distribution mark the fraction of samples that failed at least 1/10 filters applied. Labels above each vertical orange line indicate the exact value of filter threshold (orange text) and the number of samples failing this filter (black text). For the left pairs of columns, blue and red vertical lines correspond to the filter thresholds applied for PCR+ and PCR- samples, respectively. Features are ordered as follows: (**a**) median coverage per sample in 100bp bins; (**b**) dosage bias score, $\partial$; (**c**) absolute difference between smallest and largest estimated copy number for across all 22 autosomes; (**d**) median absolute Z-score of number of 1Mb bins with estimated copy number < 1.5 or > 2.5; (**e**) fraction of chimeric read pairs; (**f**) pairwise read alignment rate; (**g**) percent of library estimated to be contaminant DNA; (**h**) inferred sex chromosome ploidy. Two filtered features (mean read length & inferred-reported sex concordance) are not pictured here.

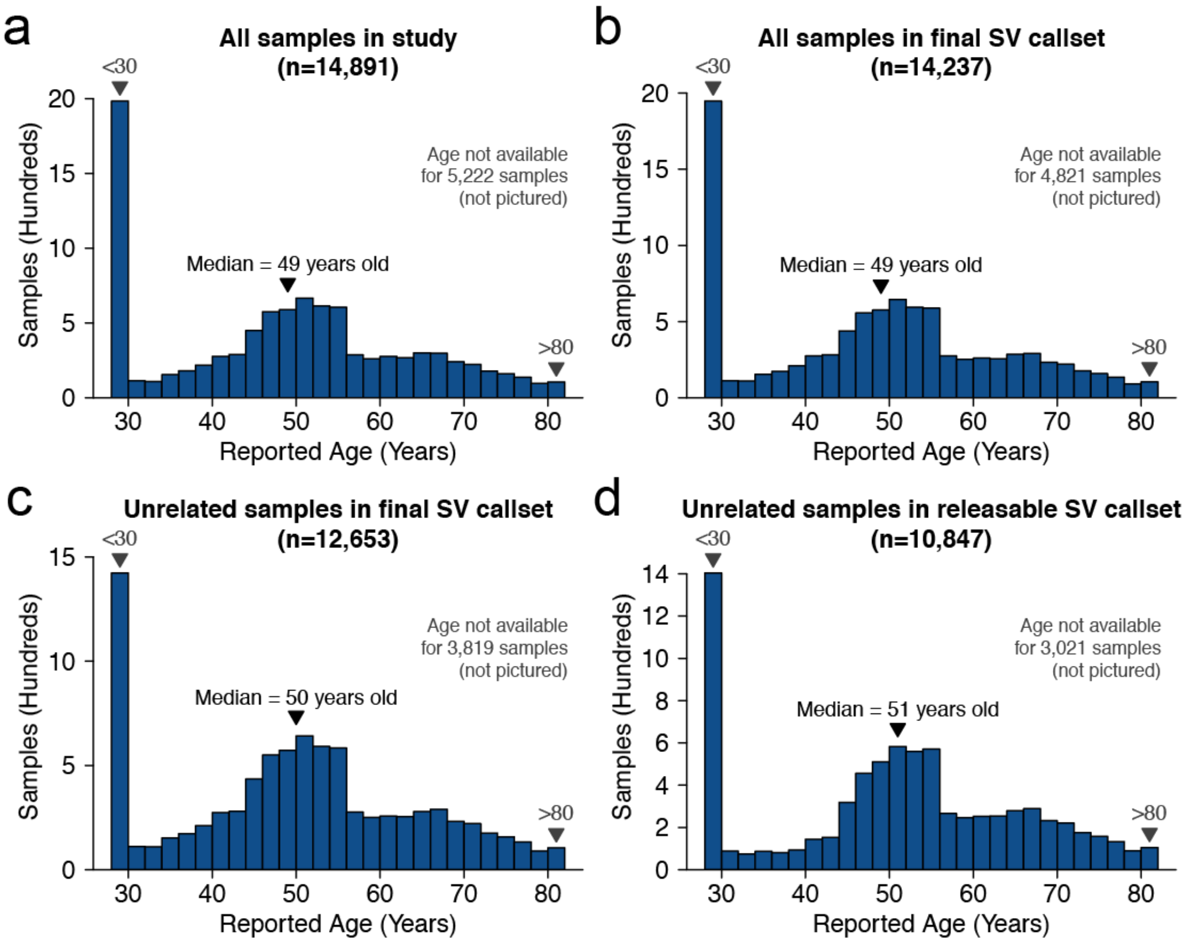


**Supplementary Figure 2 | Distribution of sample ages.** We collated reported age metadata for all samples where available, rounded to the nearest whole year, for various subsets of the gnomAD-SV dataset, including (**a**) all samples included in this study, (**b**) all samples that passed all QC measures and were included in the final gnomAD-SV callset, (**c**) all unrelated samples in the final gnomAD-SV callset used for all analyses presented in this study, and (**d**) the unrelated subset of samples with appropriate permissions to release site-frequency data on the online gnomAD Browser (https://gnomad.broadinstitute.org).

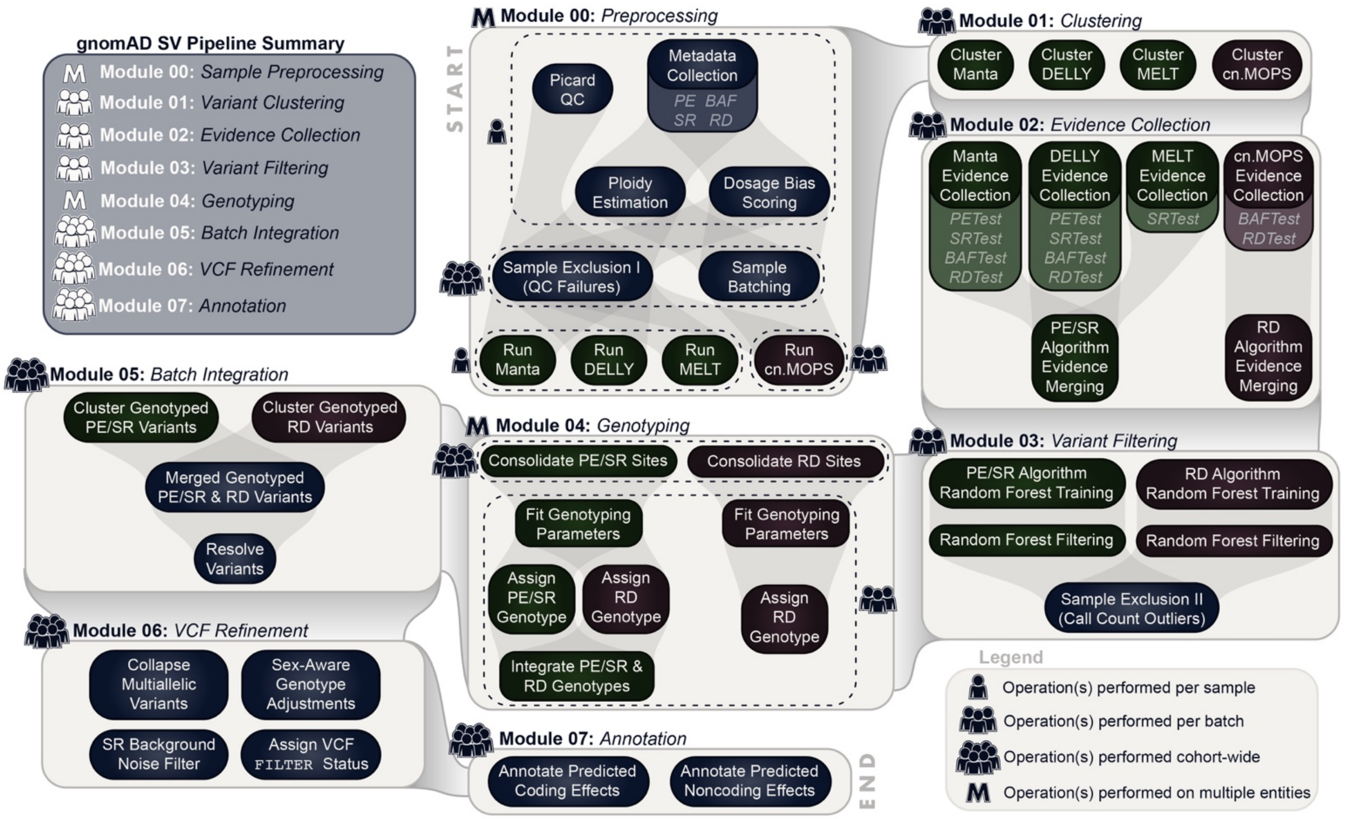


**Supplementary Figure 3 | Overview of gnomAD-SV discovery pipeline.** We extended our previously described modular SV pipeline for multi-sample joint SV discovery & genotyping in the gnomAD-SV dataset.[1] An overview of the pipeline is summarized here, but is outlined in detail in **Methods**. The gnomAD-SV discovery pipeline contains seven sequential modules (light beige boxes). The sequence of modules is listed in the top left panel and is also indicated by connections between light beige boxes. Each module contains multiple sub-modules (smaller, dark boxes) that operate on the per-sample (N=1), per-batch (N~400; see **Methods** for a description of sample batching scheme), or cohort-wide (N=14,891) level, as listed in the legend. This pipeline has been made available as a series of publicly accessible methods on FireCloud/Terra to permit cloud-based analyses of SVs across WGS studies.[2]

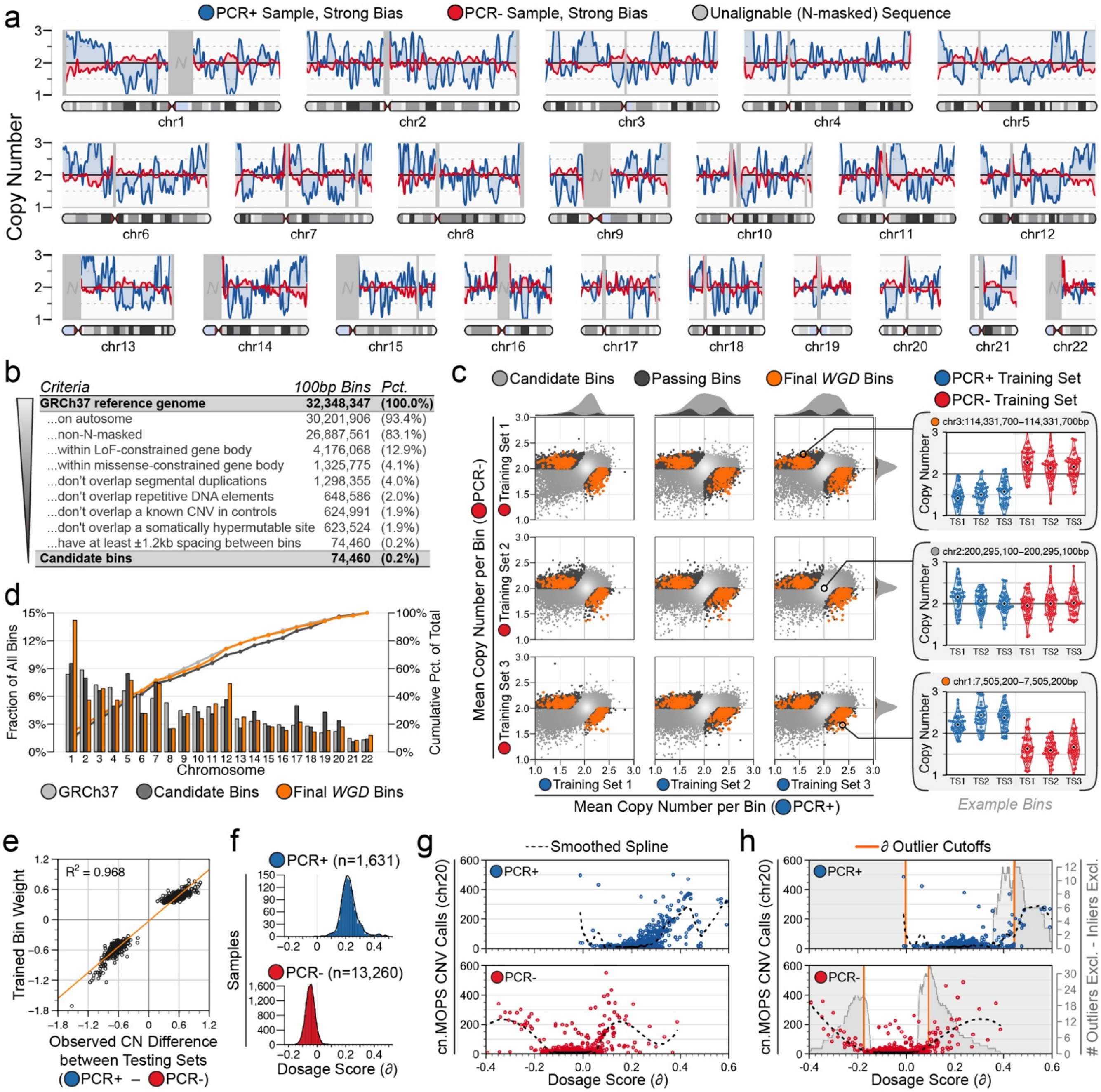


**Supplementary Figure 4 | Whole-genome dosage bias quantification.** We developed a model ("*WGD*") to quantify non-uniformity of sequencing coverage (*i.e.,* "dosage bias") per sample, and used this model to perform sample-level QC prior to SV discovery (see **Methods**). (**a**) We observed antipodal patterns of genome-wide normalized coverage throughout the gnomAD-SV dataset, consistent with our previous observations from WGS analyses of other datasets.[1,3] These patterns corresponded predominantly to PCR-amplified (PCR+) and PCR-free (PCR-) library protocols. All WGS samples in this dataset featured some degree of dosage bias, although the magnitude varied per sample. Two samples with strong dosage biases were arbitrarily selected to be shown here as examples. (**b**) To construct our model, we segmented the GRCh37 reference assembly into contiguous 100bp bins and filtered these bins to a small subset of bins depleted for technical short-read mapping artifacts and with strong prior likelihoods on being diploid in the average healthy individual. (**c**) We identified statistically informative bins by evaluating the difference in mean copy number for three training sets of 50 samples each for PCR+ and PCR- (total n=300 training samples). Per-sample copy numbers for all training samples are shown at right for three example bins. (**d**) The distribution of bins per chromosome for candidate bins passing our filters from (b) and bins in our final *WGD* model were approximately consistent with each chromosome's relative length. (**e**) Per-bin weights assigned during model training were strongly correlated with observed copy number (CN) differences between an independent pair of PCR+ and PCR- training sets (total n=100 samples). (**f**) Distribution of *WGD* scores (denoted *∂*) for all 14,891 samples in the gnomAD-SV cohort. (**g-h**) We generated raw cn.MOPS calls for chromosome 20 across all 14,891 samples split by PCR status then batched randomly (g) and ranked by *∂* then batched sequentially (h). Ranking samples by *∂* prior to batching for read depth-based CNV discovery (g) improved the uniformity of raw CNV calls per sample, and better controlled outlier samples. From these data, we also learned minimum and maximum *∂* thresholds for sample-level QC prior to SV discovery (g) that maximized the number of cn.MOPS outlier samples excluded while minimizing the number of well-behaved samples lost.

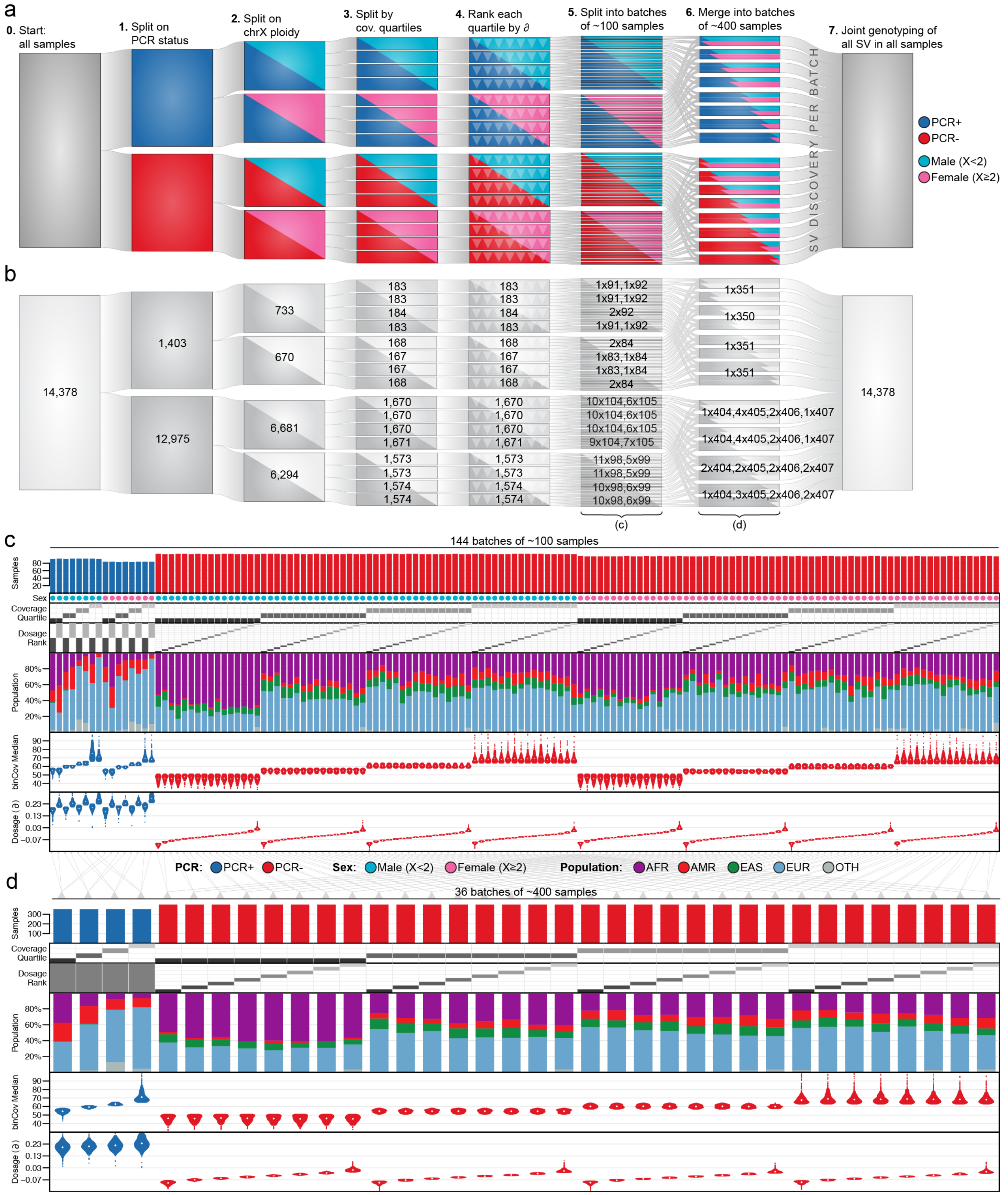


**Supplementary Figure 5 | Sample batching strategy.** We devised a strategy for subdividing samples into smaller batches for joint discovery of SVs to control for technical variability between samples (*e.g.,* dosage biases, PCR status, depth of coverage) and to leverage increased parallel computation in the cloud. (**a**) Overview of the batching procedure, which is described in detail in **Methods**. (**b**) Annotation of number of samples per split point in the batching procedure as applied to the full gnomAD-SV cohort after preliminary sample QC (see **Supplementary Figure 1**). (**c-d**) Distributions of metadata & sequencing metrics per (c) cn.MOPS batch and (d) gnomAD-SV pipeline batch. Populations correspond to population assignments inferred from the final gnomAD-SV callset (see **Methods**).

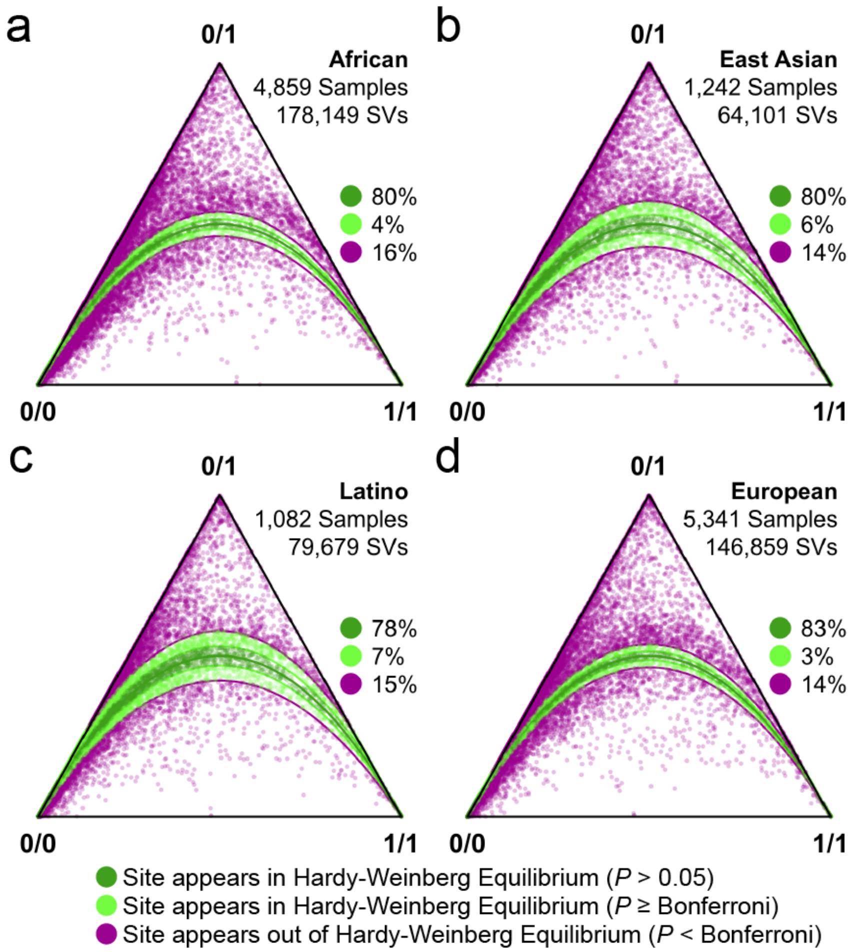


**Supplementary Figure 6 | Hardy-Weinberg Equilibria of SVs across major continental populations.** Despite being an imperfect measurement to assess genotyping accuracy, we computed Hardy-Weinberg Equilibrium (HWE) statistics for all biallelic autosomal SVs documented in this study for each of the four major populations considered: (**a**) African/African-American, (**b**) East Asian, (**c**) Latino, and (**d**) European. These data are presented here as De Finetti diagrams,[4] where each point is a single biallelic autosomal SV projected onto HWE ternary axes corresponding to its ratio of homozygous reference (0/0), heterozygous (0/1), and homozygous alternate (1/1) genotypes across all samples in the indicated population. The distance of a point to a vertex indicates the fraction of samples with that genotype. Points are colored based on their adjusted p-value compared to HWE expectations ($1 = p^2 + 2pq + q^2$). Green points are SVs within bounds defined for HWE based on the number of sites documented in each population, and purple points are SVs outside of these p-value bounds. The proportion of SVs corresponding to each p-value cutoff is provided at the right of each panel. Plots were generated using the "HardyWeinberg" package in R.[5] See **Extended Data Figure 2b** for a combined HWE ternary plot across all samples.

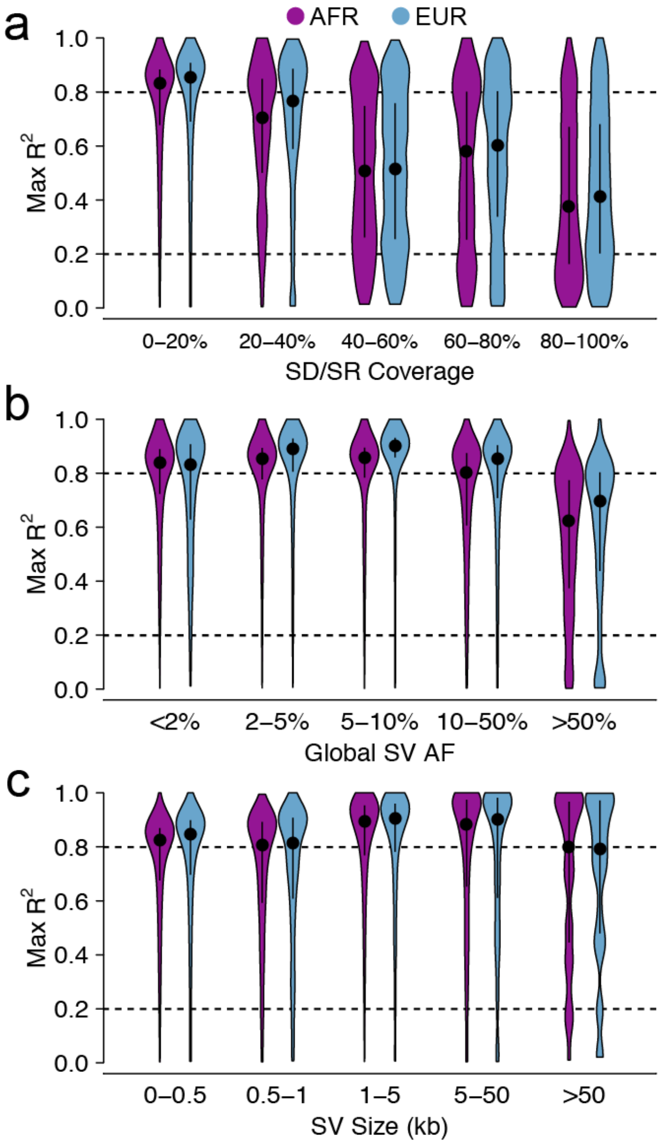


**Supplementary Figure 7 | Patterns of linkage disequilibrium between SVs and SNVs/indels.** We computed linkage disequilibrium (LD) between common (allele frequency [AF] ≥1%) autosomal SVs and all SNVs/indels within ±1Mb from a subset of 5,353 African/African-American (AFR; n=3,470) and European (EUR; n=1,883) samples overlapping between this study and a sister study.[6] (**a**) Maximum $R^2$ values for SVs and nearby SNVs/indels, stratified by population and repeat coverage. Points reflect medians and vertical black bars reflect interquartile range. Repetitive primary sequence contexts, including segmental duplications (SD) and simple repeats (SR), had a strong influence on LD between SVs and nearby SNVs and indels, which was likely due to a combination of the complex haplotype structures in these regions as well as the increased technical difficulty of genotyping variants in repetitive sequences from short reads. To account for this, we restricted all subsequent analyses of LD between SVs and SNVs/indels to SVs that were <30% covered by annotated SD and SR elements. (**b-c**) Maximum $R^2$ between common SVs and SNVs/indels, stratified by population and (b) global SV AF or (c) SV size, after restricting on SD/SR coverage as described above.

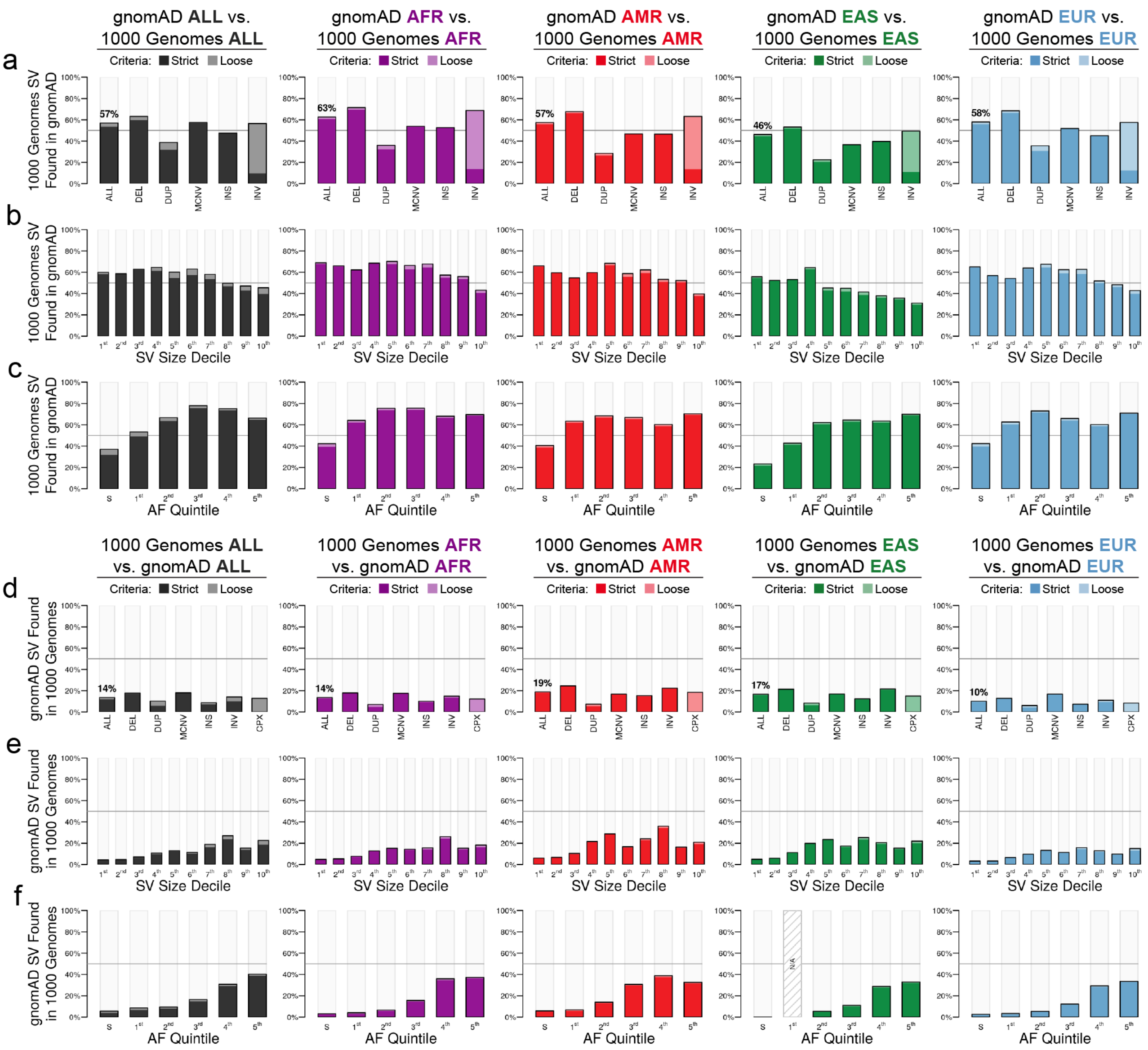


**Supplementary Figure 8 | Site-level comparison of SVs to the 1000 Genomes Project.** We compared the gnomAD-SV callset to the SVs from the 1000 Genomes Project phase III release.[7] We considered two ‘directions’ of comparison: (**a-c**) the fraction of SVs reported by the 1000 Genomes Project that were also observed in gnomAD, and (**d-f**) the fraction of SVs discovered in gnomAD-SV that were also reported by the 1000 Genomes Project. For each comparison, we further stratified across three dimensions: (**a & d**) SV class, (**b & e**) SV size binned by decile, and (**c & f**) AF, binned into quintiles after holding out singletons as their own bin (marked with an “S”). We evaluated these comparisons across all samples (“ALL”), as well as when matching on four major populations considered in both studies (AFR, AMR, EAS, and EUR). Sites matching with strict criteria are marked with dark colors, whereas sites matching with looser criteria are marked with lighter colors (see **Methods** for details).

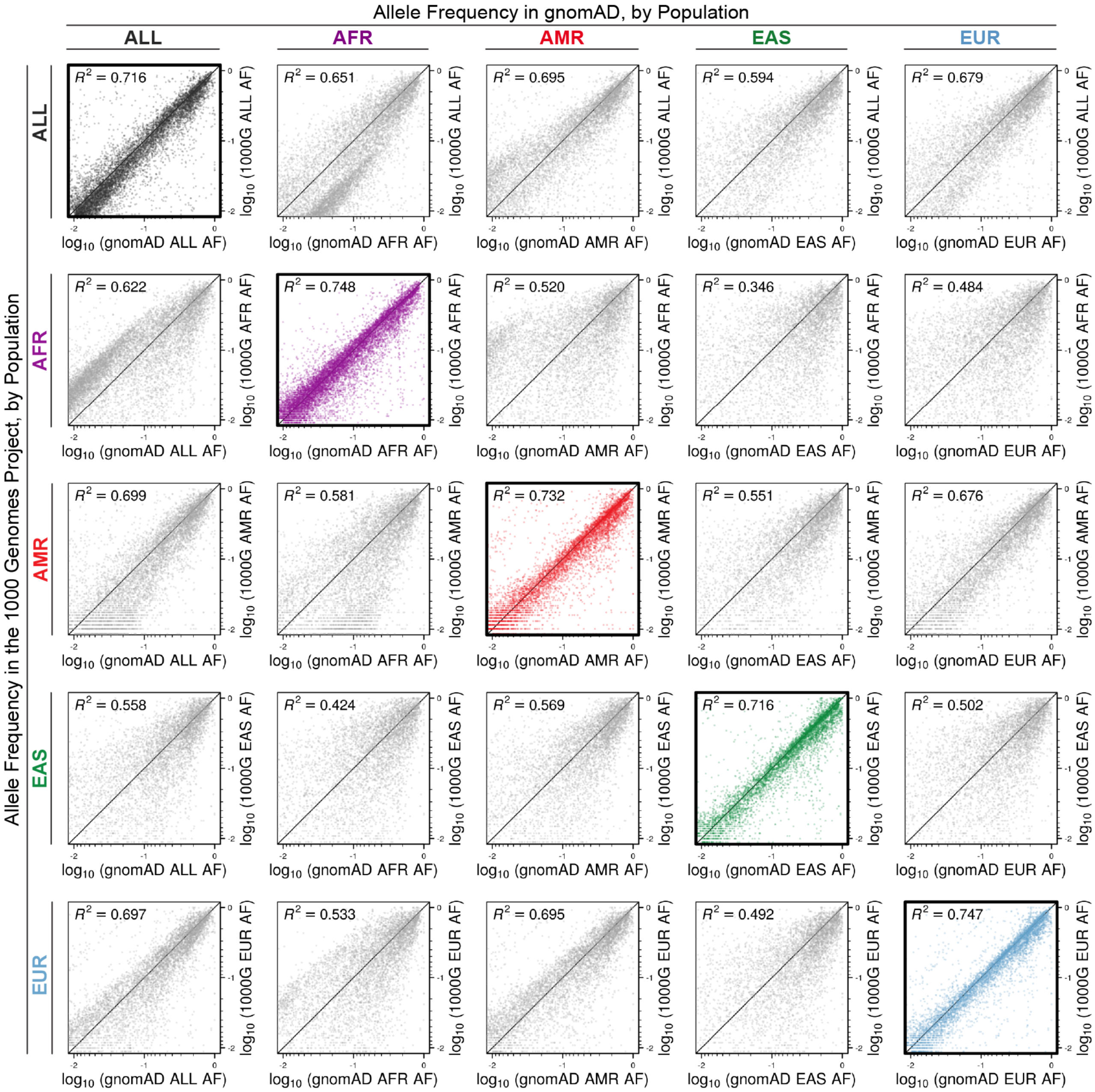


**Supplementary Figure 9 | Allele frequency comparisons to SVs from the 1000 Genomes Project.** In addition to site-level comparisons (see **Supplementary Figure 8**), we also compared allele frequencies (AFs) between common (AF > 1%), biallelic, autosomal SVs discovered in both gnomAD-SV and the 1000 Genomes Project phase III release.[7] We found a positive correlation between AFs of SVs discovered in both studies, which was strongest when matching on population. We compared all pairs across four major populations considered in both studies (AFR, AMR, EAS, and EUR), as well as all samples across all populations (“ALL”). Comparisons where populations were matched are marked with a thick border and colored points; all other comparisons represent inter-population comparisons. Correlations were assessed with a Pearson correlation test.

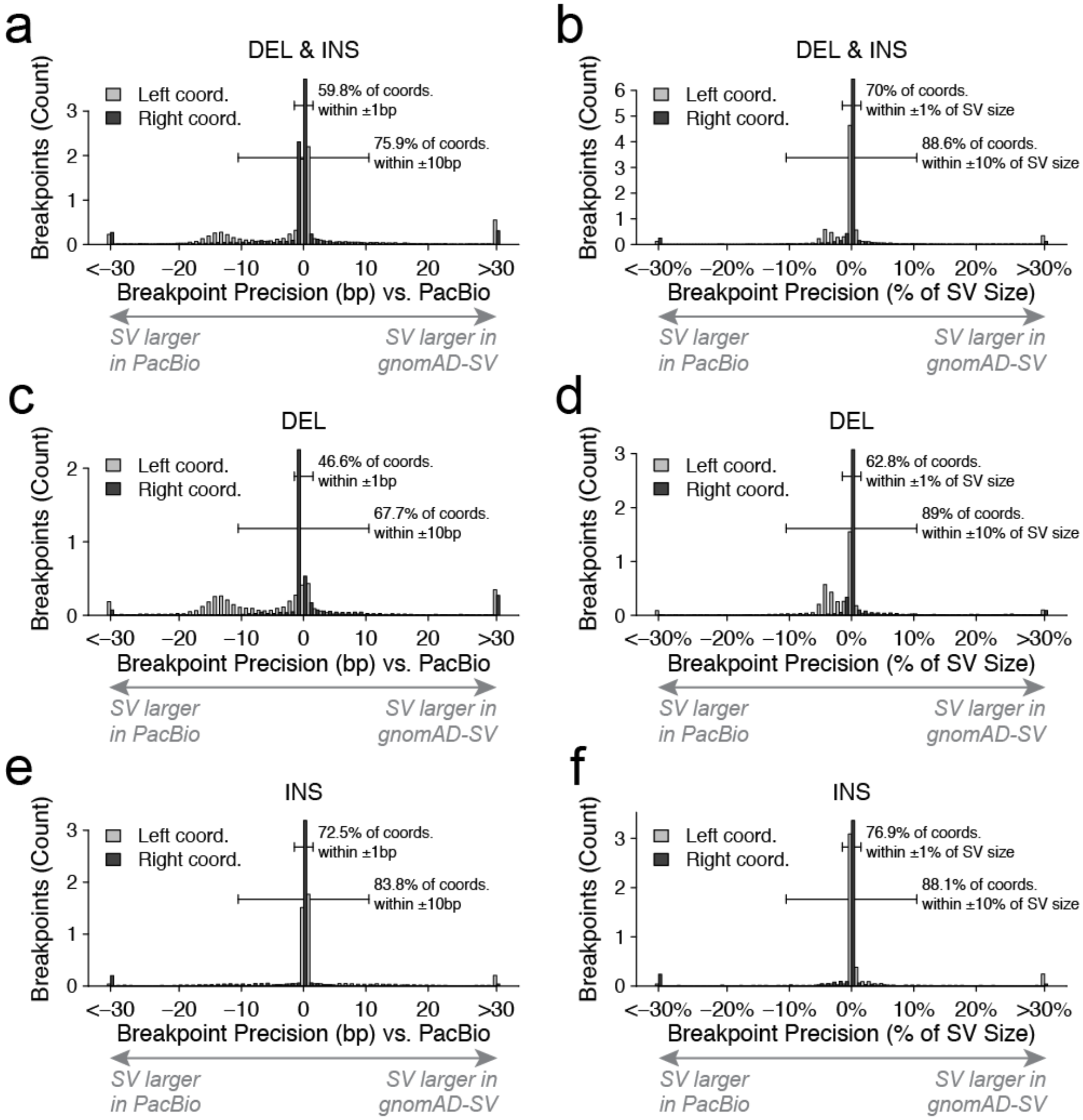


**Supplementary Figure 10 | Breakpoint accuracy estimates from long-read WGS.** We estimated breakpoint accuracy in gnomAD-SV by comparing to Pacific Biosciences ("PacBio") long-read WGS-derived SV callsets produced by the same assembly & SV calling algorithms for two samples also present in gnomAD-SV.[8,9] For insertion and deletion SVs with long-read support for their breakpoints (see **Extended Data Figure 3**), we calculated the distance between reported breakpoints from short-read and long-read WGS separately for the left (*i.e.*, lower) and right (*i.e.*, higher) coordinate (abbreviated "coord."). (**a**, **c**, **e**) Absolute breakpoint accuracy estimated vs long-read WGS for the left and right coordinates per breakpoint, split by SV class. (**b**, **d**, **f**) Breakpoint accuracy normalized to the overall size of each SV, split by SV class.

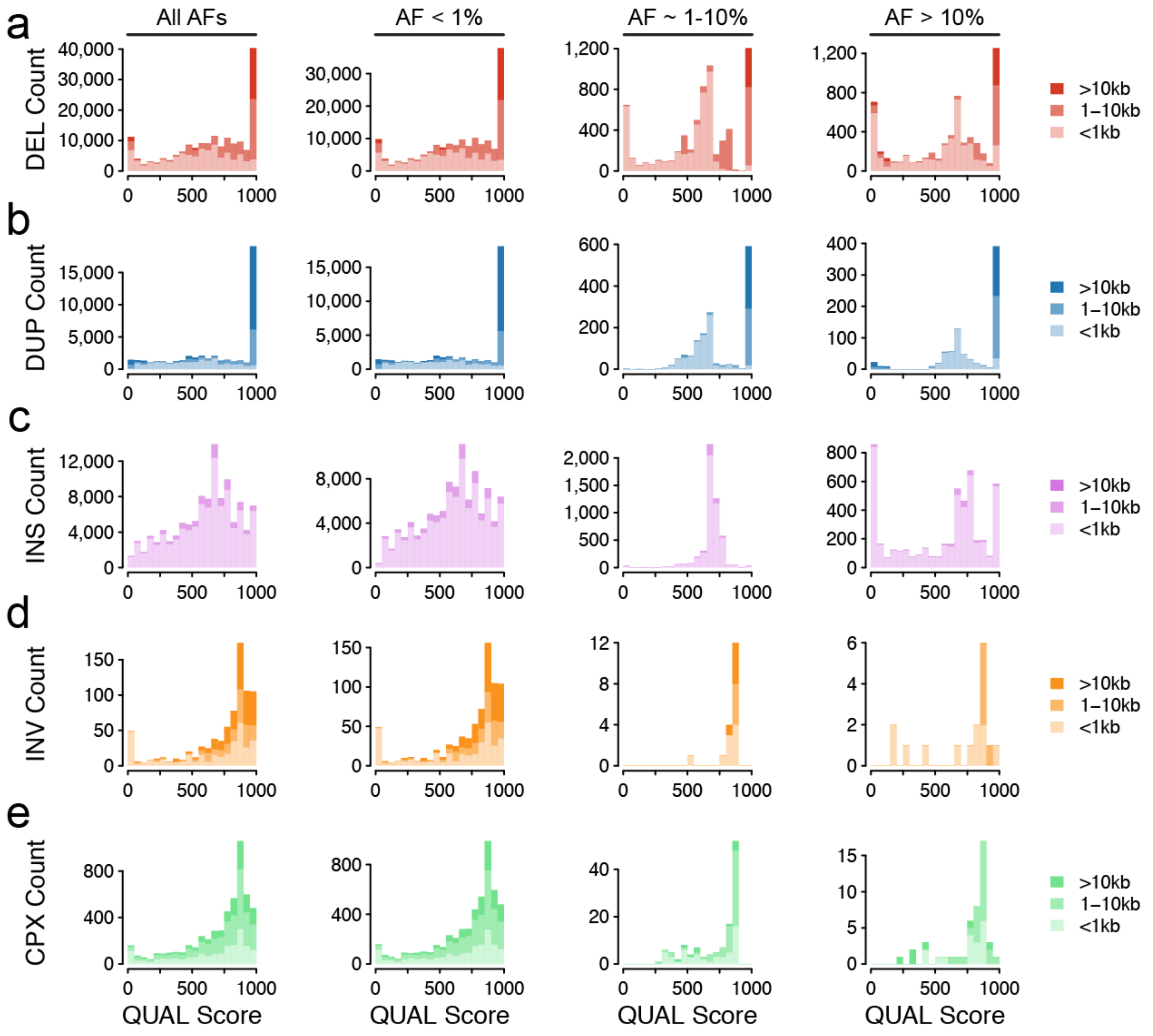


**Supplementary Figure 11 | Distribution of variant quality scores.** Distributions of quality scores ("QUAL" scores in VCF format) stratified by AF (columns) and variant size (shading) for all five major SV classes, including (**a**) deletions, (**b**) duplications, (**c**) insertions, (**d**) canonical inversions, and (**e**) complex SV. MCNVs and reciprocal translocations are not pictured here, as both variant classes are sparse and/or QUAL scores are uninformative.

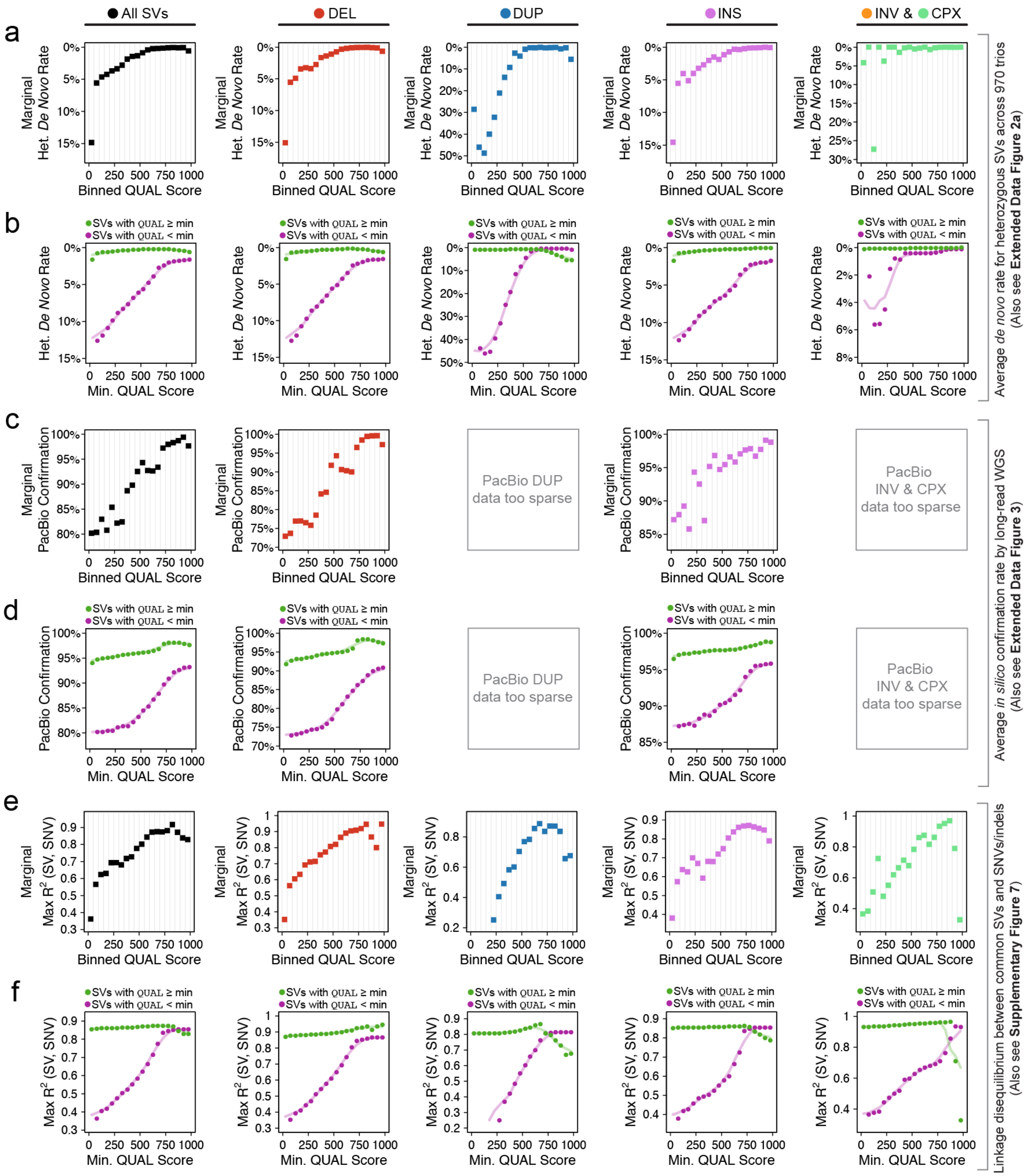


**Supplementary Figure 12 | Validation of variant quality score calibration.** We sought to assess whether our variant quality scores ("QUAL" scores in VCF format; see **Supplementary Figure 11**) were consistent with available orthogonal measurements of variant confidence. We considered three independent analyses: (**a-b**) apparent *de novo* rates for heterozygous SVs across 970 parent-child trios (also see **Extended Data Figure 2a**), (**c-d**) *in silico* variant confirmation by Pacific Biosciences (PacBio) long-read WGS on a subset of samples (also see **Extended Data Figure 3** and

**Supplementary Figure 10**), and (**e-f**) patterns of linkage disequilibrium (LD) between common SVs and nearby SNVs or indels (also see **Supplementary Figure 7**). For each analysis, we stratified variants by SV class, then partitioned each into sequential, non-overlapping bins by increments of +50 QUAL score from the global minimum score (QUAL=0) to the global maximum score (QUAL=1000). Given the requirements for *in silico* long-read WGS confirmation, we uniformly restricted all analyses presented here to SVs with breakpoint-level read support (*i.e.*, "split-read" evidence; includes ~93% of all SVs) and SVs with breakpoints not localized to annotated segmental duplications and/or simple repeats. After applying these filters, for each QUAL score bin, we either computed the (**a**-**b**) mean heterozygous *de novo* rate, (**c**-**d**) the fraction of SVs confirmed by long-read WGS, or (**e-f**) the median peak LD between SVs and SNVs/indels within ±1Mb. Panels (**a**), (**c**), and (**e**) represent these data as the marginal performance for each metric within each QUAL score bin; *i.e.*, the variants included in each bin in these panels are not included in any other bins within the same graph. Panels (**b**), (**d**), and (**f**) represent these data as the cumulative average of all SVs either above (green) or below (purple) a specified minimum QUAL threshold for theoretical *post hoc* filtering of the gnomAD-SV callset, and solid lines sliding window averages over 5 bin windows (corresponding to windows of 250 on the QUAL score scale, in steps of 50).

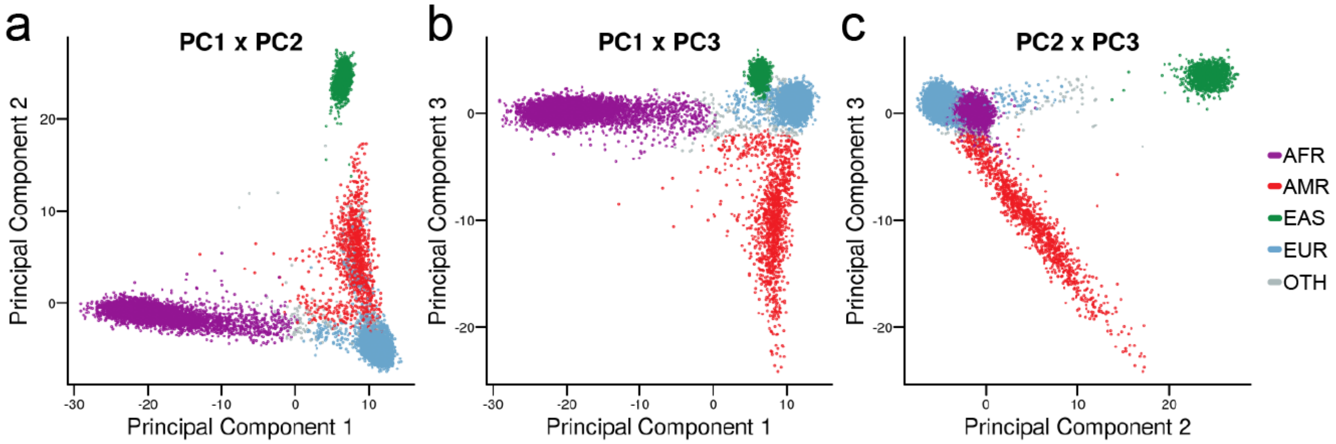


**Supplementary Figure 13 | Principal components analysis of common SVs.** We performed a principal component (PC) analysis of common (AF>1%), high-quality SVs across all samples in the final gnomAD-SV callset before pruning related individuals. The top three principal components from this analysis clearly stratified samples into continental ancestry groups. Show here are all three pairwise combinations of the top three principal components, colored by population assignment (also see **Figure 1d**).

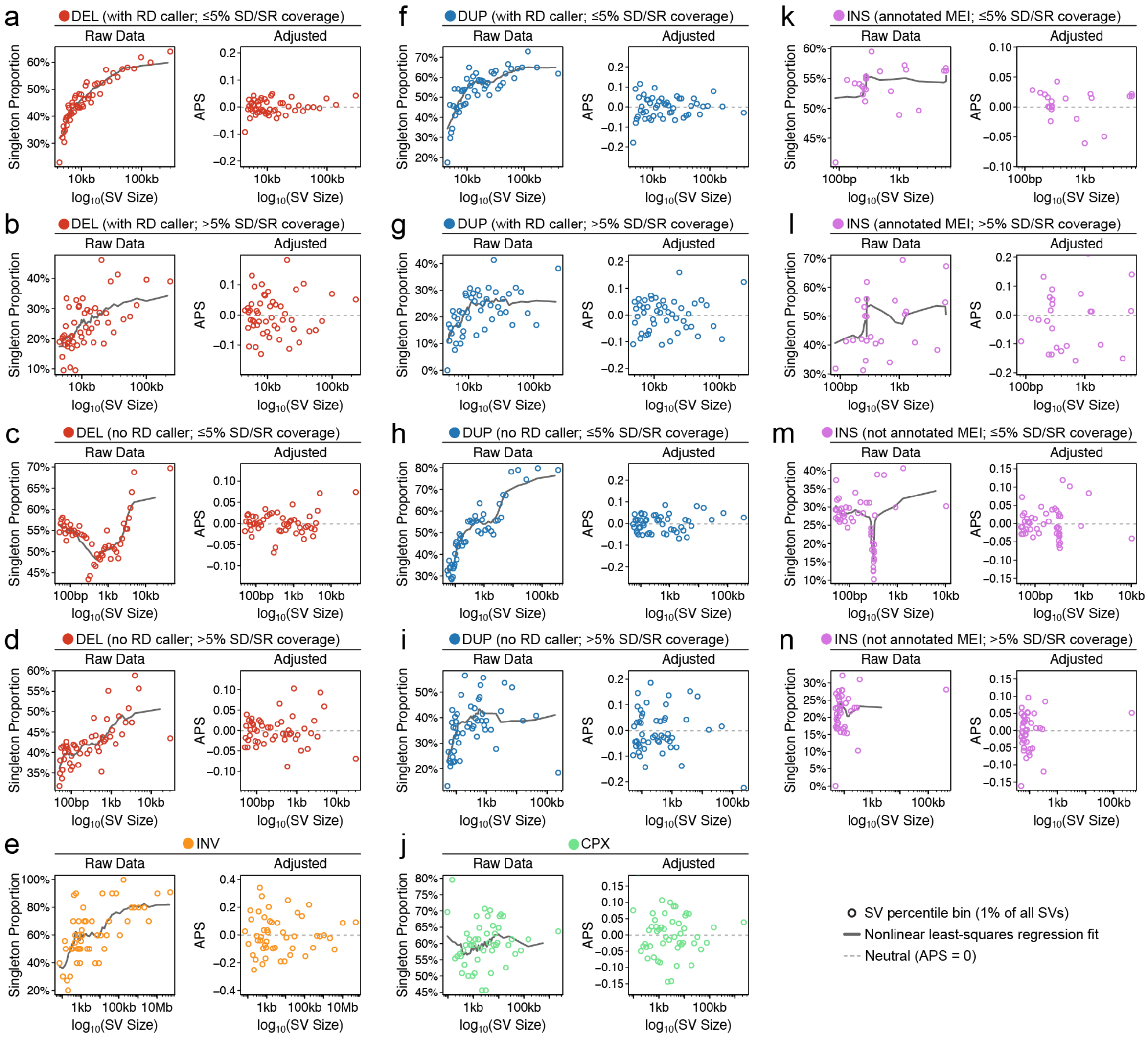


**Supplementary Figure 14 | Development of the adjusted proportion of singletons (APS) metric.** We observed that the singleton proportion for an arbitrary set of SVs was highly dependent on SV size, class, contributing evidence supporting event (*e.g.*, read depth, split reads), and genomic context. To account for these relationships in our analyses, and to permit comparisons of frequency spectra across SV classes, sizes, and contexts, we fit a nonlinear least-squares regression model separately to each of 14 SV categories, including: deletions contributed by RD callers with **(a)** ≤5% or **(b)** >5% coverage by annotated segmental duplications and simple repeats (SD/SR); deletions not contributed by RD callers with **(c)** ≤5% or **(d)** >5% SD/SR coverage; **(e)** inversions; duplications contributed by RD callers with **(f)** ≤5% or **(g)** >5% SD/SR coverage; duplications not contributed by RD callers with **(h)** ≤5% or **(i)** >5% SD/SR coverage; **(j)** complex SVs; insertions annotated as mobile element insertions (MEIs) with **(k)** ≤5% or **(l)** >5% SD/SR coverage; insertions not annotated as mobile elements with **(m)** ≤5% SD/SR coverage or **(n)** >5% SD/SR coverage. The left side of each panel displays the unadjusted relationship between SV size and singleton proportion after dividing all SVs into 100 uniform bins based on SV size, and the grey line indicates the nonlinear least-squares fit to those data. The right side of each panel displays the same data after adjustment.

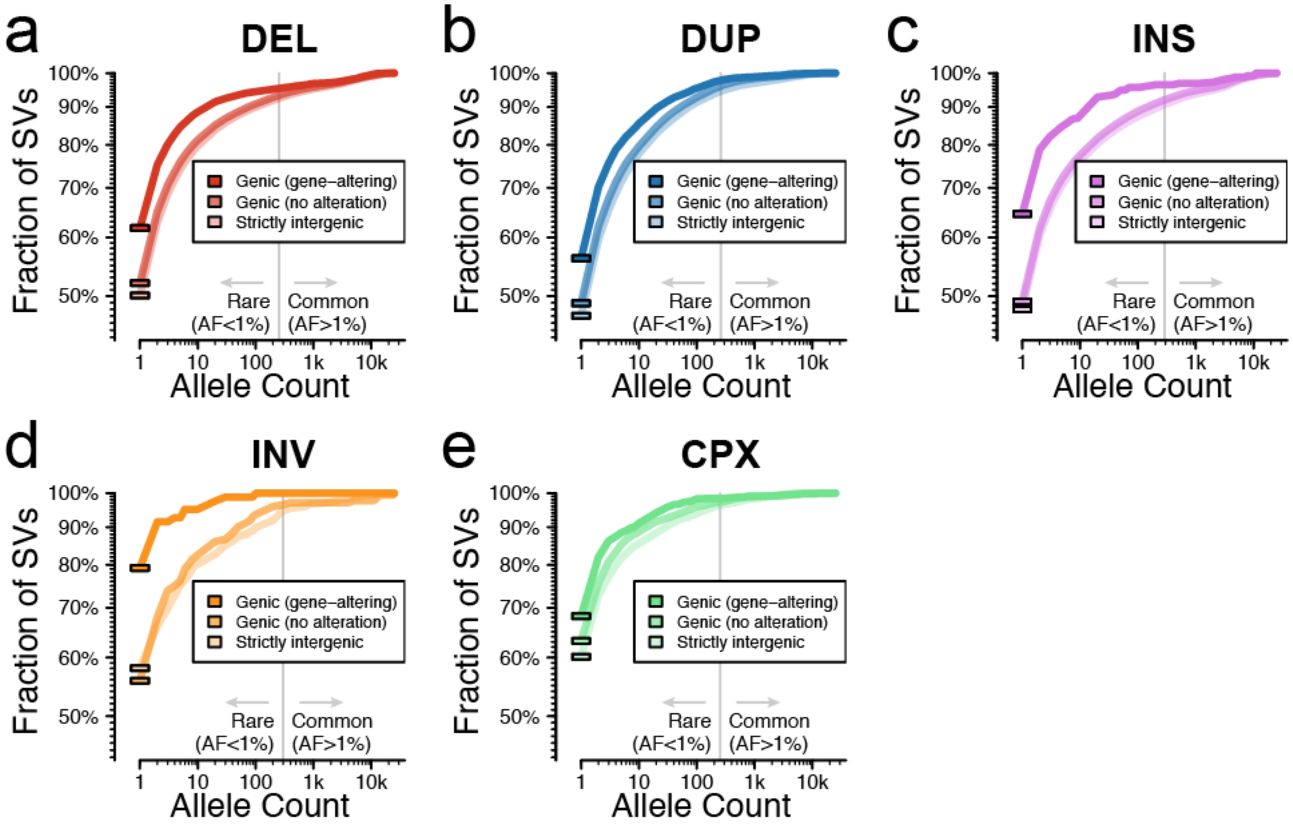


**Supplementary Figure 15 | Comparison of allele frequencies per SV class by genic context.** We compared the AF distributions across SV classes conditioned by their relationship to autosomal protein-coding genes. Each panel corresponds to a single SV class, which we further decomposed into three categories based on their genic context: (i) predicted gene-altering SVs, which included predicted loss-of-function (pLoF), whole-gene copy gain (CG), or intragenic exonic duplication (IED) SVs of at least one gene (see **Supplementary Figure 17**), (ii) SVs that overlapped genes but were not predicted to result in a disruptive functional consequence, which included intronic SV, gene-spanning inversions, or partial duplications not resulting in CG or IED, and (iii) SVs with no overlap with any protein-coding genes.

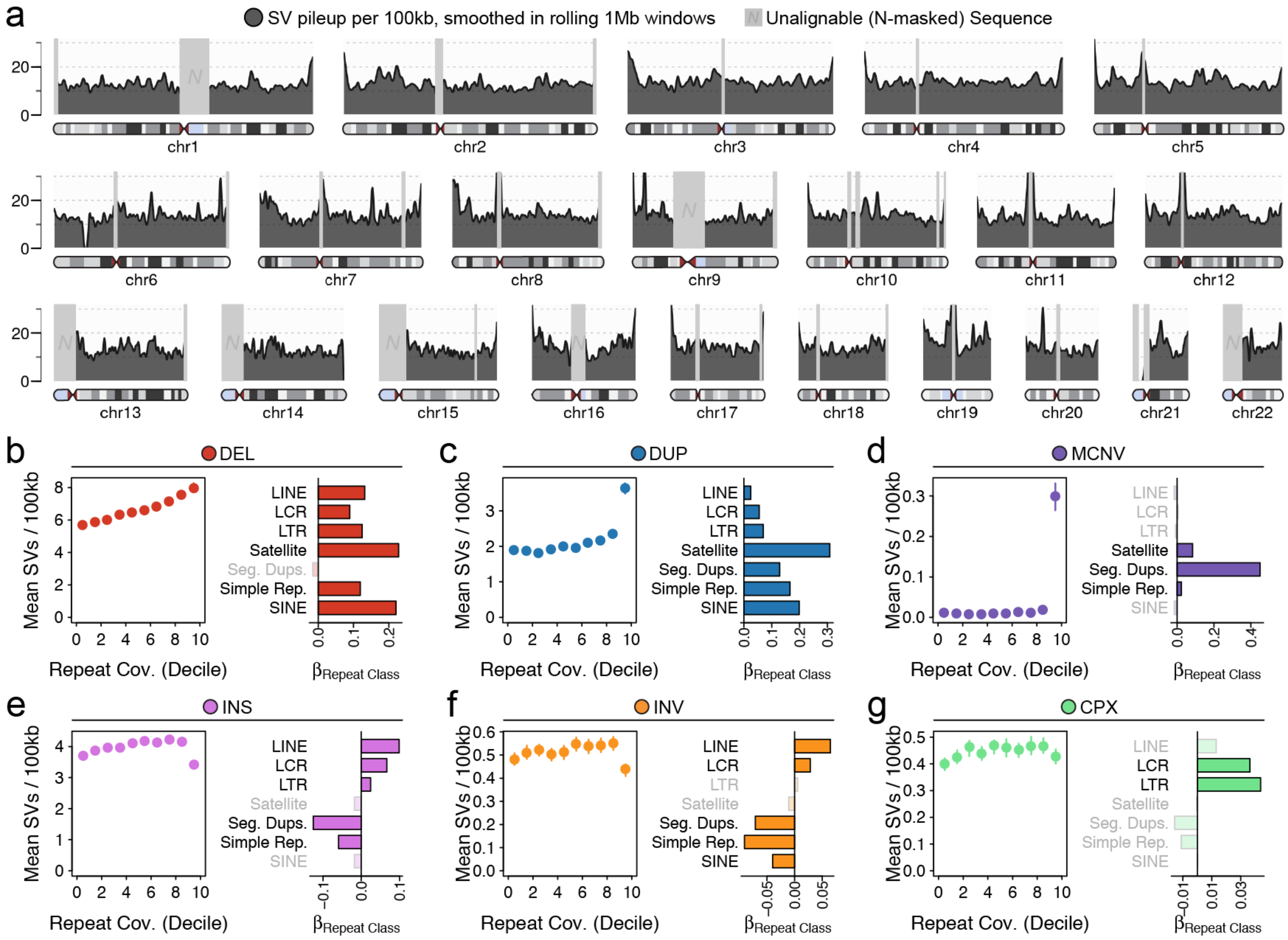


**Supplementary Figure 16 | Genomic patterns and correlates of SV density.** (**a**) We computed the density of SVs per autosome in 100kb sequential windows, represented here after being smoothed as a 1Mb rolling mean. SV density varied by chromosome and position, with centromeres and telomeres being particularly enriched for SVs (also see **Figure 3c-d**). Unalignable regions of the GRCh37 reference genome are masked with light grey. (**b-g**) Analyses of SV density versus classes of annotated repetitive elements. For each SV class, the left panel displays the mean number of SVs per 100kb window divided into deciles based on coverage by annotated repetitive elements. Bars represent 95% confidence intervals from 100-fold bootstrapping. The right panel displays regression coefficients from a multivariate linear model of SV density versus seven annotated repeat classes. Dark shaded bars indicate repeat classes that were significantly associated with SV density after Bonferroni correction, while light shaded bars are non-significant. LCR = low-complexity repeat; LTR = long terminal repeat; Seg. Dups. = segmental duplications. The relationships between SV density and repeat class inferred here are likely to be influenced in part by technical limitations of short-read WGS in low-complexity sequences.

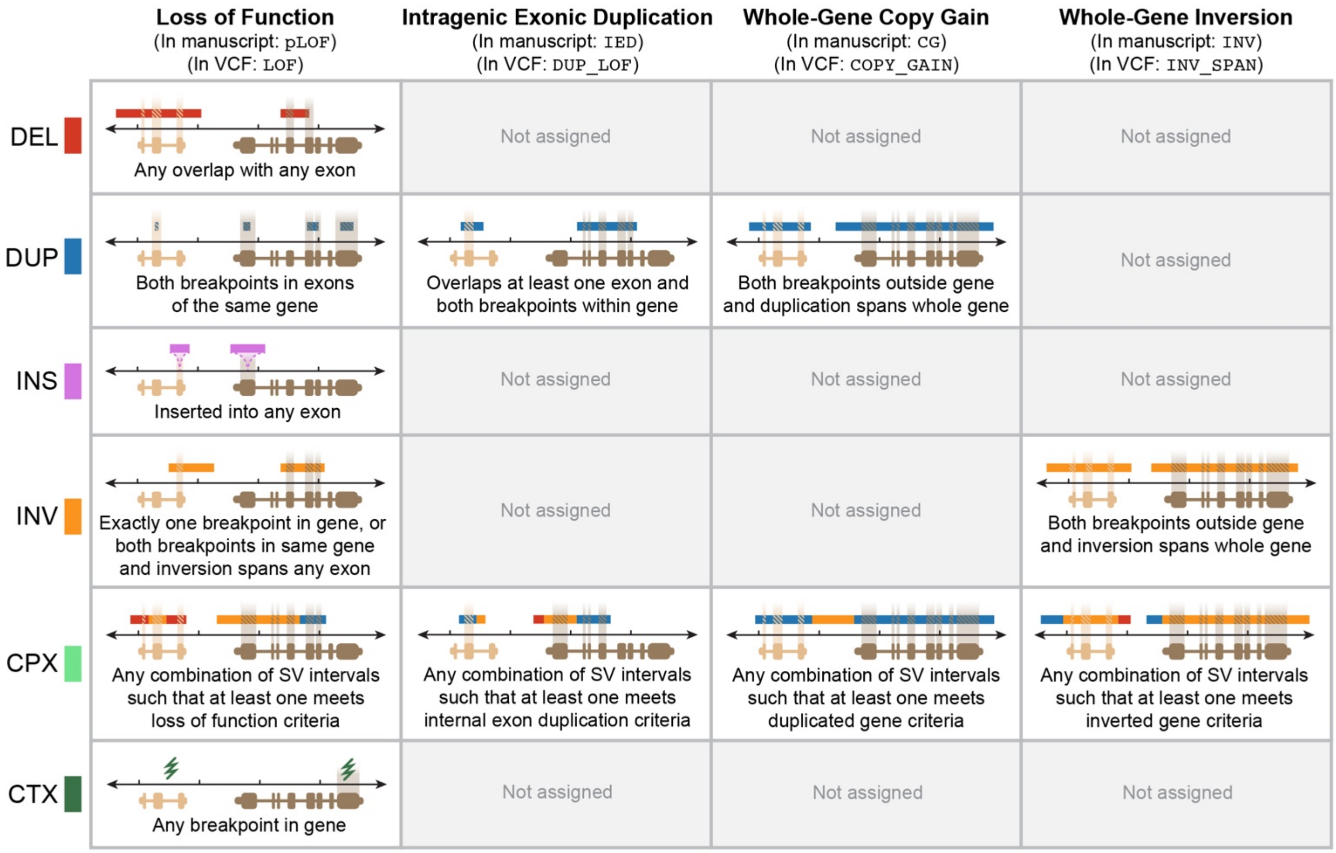


**Supplementary Figure 17 | Summary of SV annotations in coding sequences.** We annotated all SV for multiple possible functional effects on the canonical transcripts of protein-coding genes (see **Methods**). The possible effects assigned per SV class are illustrated here, with example schematics of qualifying variants for each category.

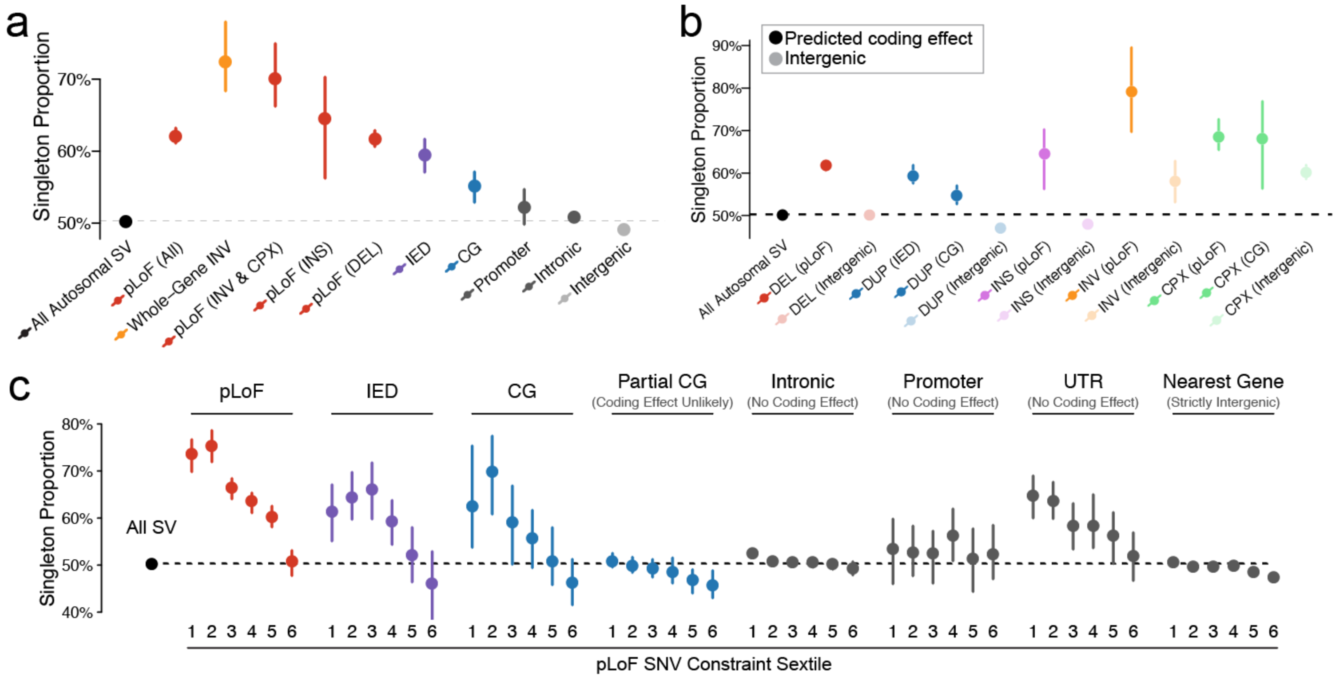


**Supplementary Figure 18 | Raw singleton proportions for selected findings.** Here, we present multiple findings from this study on the scale of raw (unadjusted) singleton proportion, rather than on the APS metric scale. (**a**) Equivalent of **Figure 4c**. (**b**) Equivalent of **Extended Data Figure 6c**. (**c**) Equivalent of **Extended Data Figure 6f**. Refer to the legend of each referenced figure for more information.

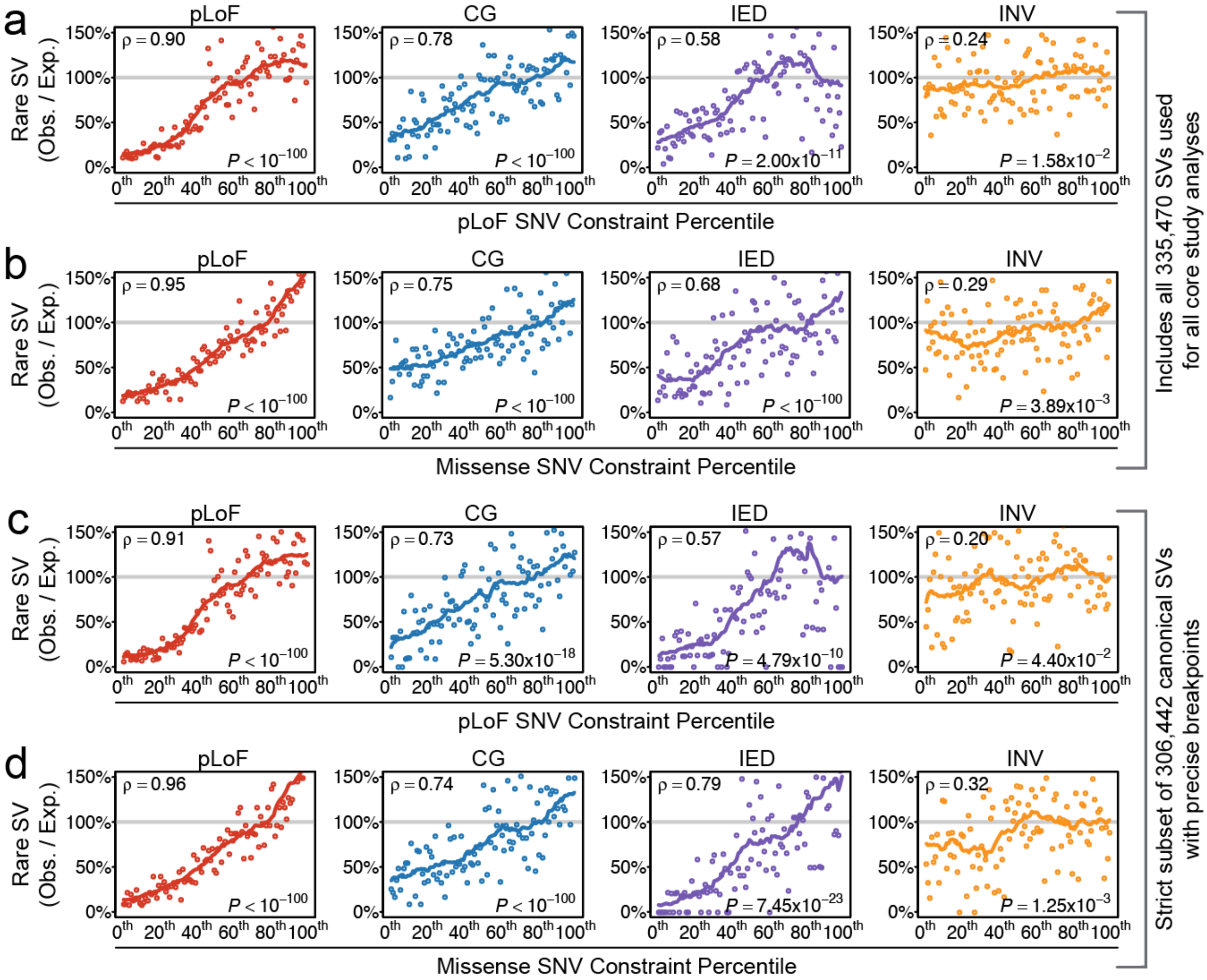


**Supplementary Figure 19 | Comparisons of SV depletion vs SNV pLoF and missense constraint.** As described in **Figure 4d**, we compared a relative measure of rare SV depletion within genes to (**a, c**) pLoF SNV constraint and (**b, d**) missense SNV constraint.[6] We performed this analysis separately for each of four possible SV-gene annotations: pLoF, CG, IED, and whole-gene inversion (INV), as described in **Supplementary Figure 17**. Panels are formatted as described in **Figure 4d**. Panels (**a-b**) include all SVs used in the main analyses presented in this study, whereas panels (**c-d**) restrict to canonical (i.e., non-complex) SVs with precise breakpoints (i.e., SVs with "split-read" evidence), with the second set of strict filters applied to exclude potential annotation errors either due to complex rearrangements or imprecise breakpoint coordinates. We found that the results were highly similar irrespective of filtering and conclude that these findings are relatively robust to the potential sources of technical confounding considered here.

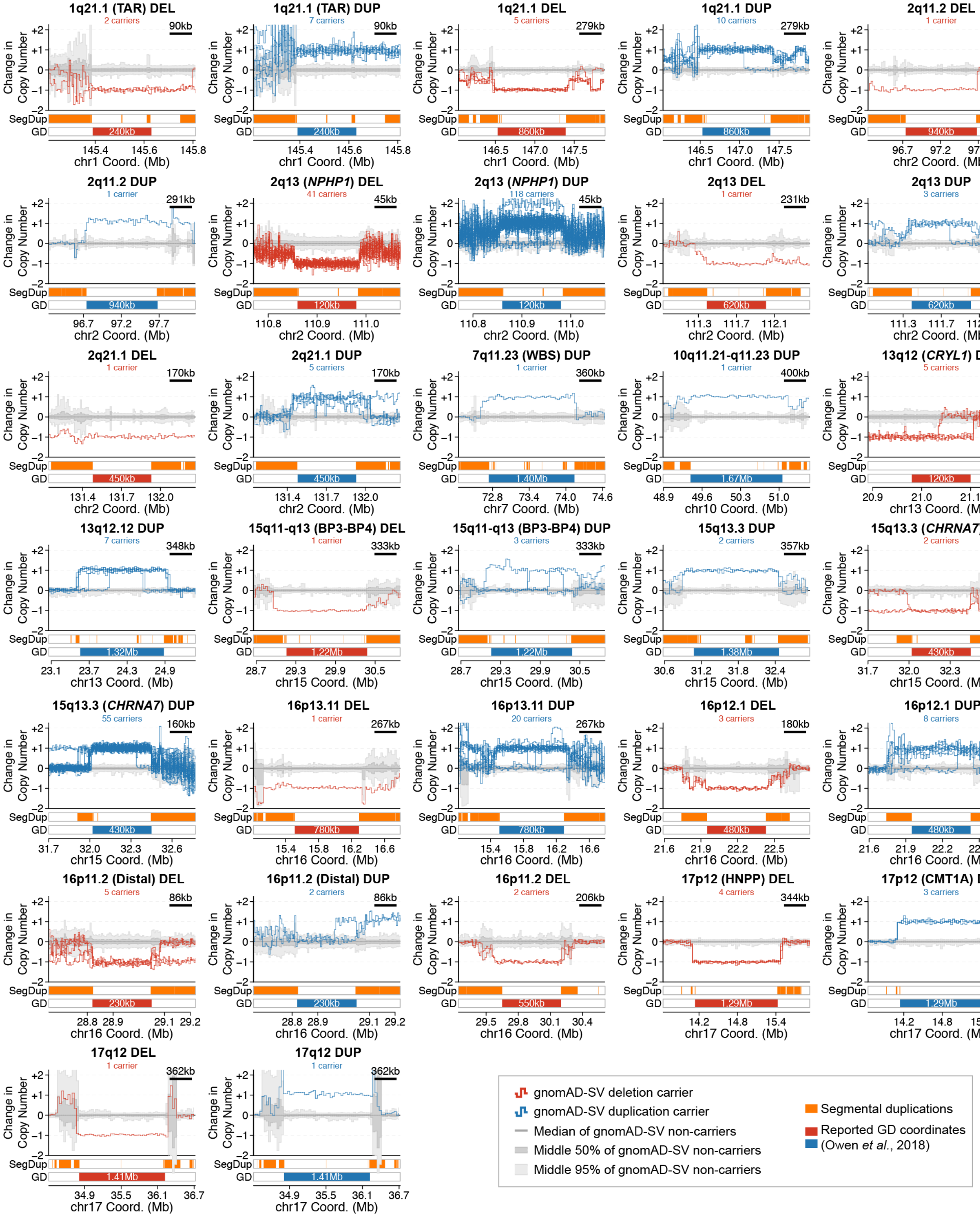


**Supplementary Figure 20 | Evidence for genomic disorder CNV carriers in gnomAD-SV.** Here we provide normalized copy-number estimates for the 32 genomic disorders (GDs) with at least one non-reference carrier predicted among the subset of gnomAD-SV samples after excluding cases of known neurological disease (also see **Supplementary Table 6**). Each panel represents the deviation from the median copy state across all samples at a single GD locus. Individual predicted CNV carriers are shown with red or blue lines. The distribution of all predicted non-carriers is shown with grey shading: the dark grey line indicates the median across all samples, the medium grey shading indicates the middle 50% of all non-carrier samples, and the light grey shading indicates the middle 95% of all non-carrier samples. Gain or loss of integer copy states are indicated with horizontal dashed grey lines, for reference. As many genomic disorders are mediated by non-allelic homologous recombination, segmental duplications are marked in orange below each plot.

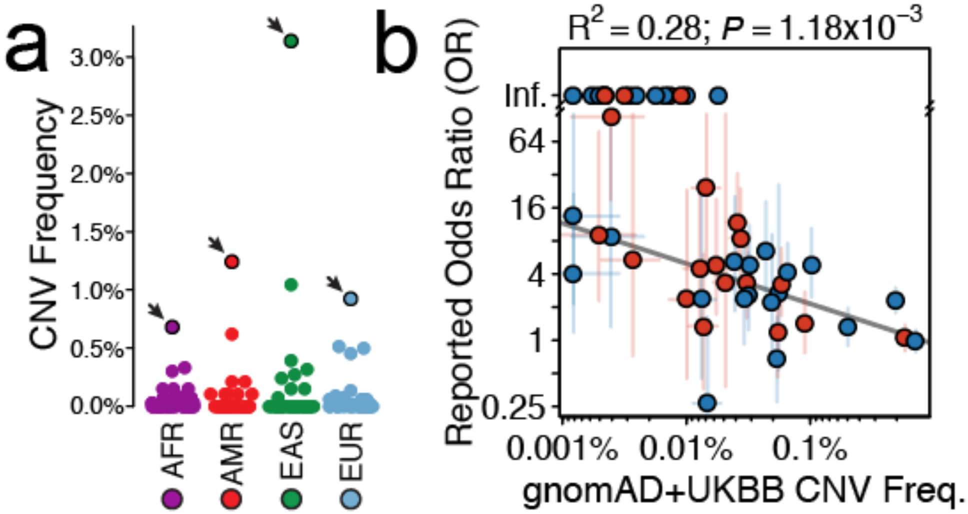


**Supplementary Figure 21 | Properties of genomic disorders evaluated in gnomAD-SV.** (**a**) GD CNV frequencies were comparable across populations in gnomAD-SV, except for duplications at 2q13 (NPHP1), where the frequency in East Asian samples was up to 5-fold greater than other populations (2q13 *NPHP1* duplications marked with solid black outlines and arrows). (**b**) The odds ratios (ORs) for these 49 GDs in developmental disorder (DD) patients from Coe et al.[10] were inversely correlated with the combined CNV frequencies in the gnomAD-SV and UKBB datasets ($R^2$=0.28; P=1.18x$10^{-3}$; Pearson correlation test). Solid grey line represents linear best fit.

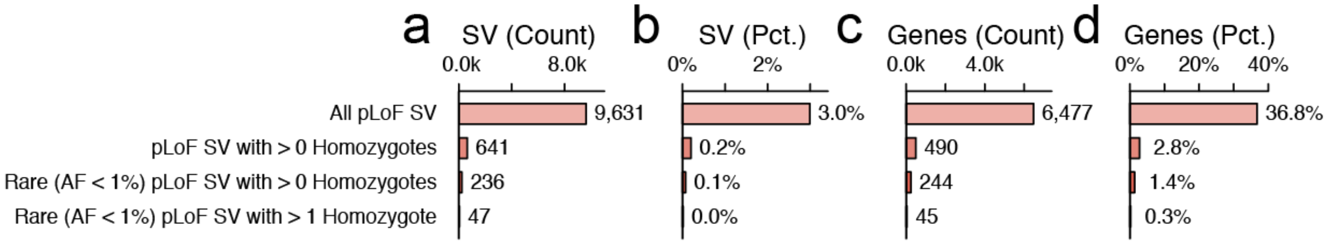


**Supplementary Figure 22 | Homozygous pLoF SVs.** We compiled a list of genes predicted to be completely inactivated in at least one individual due to a homozygous pLoF SV. (**a**) Counts of SVs resulting in pLoF, with three tiers of filters as listed. (**b**) Percent of total SVs meeting each criterion listed in (a). (**c**) Counts of unique genes with at least one SV meeting the criteria in (a). (**d**) Percent of all autosomal, protein-coding genes with at least one homozygous pLoF SV.

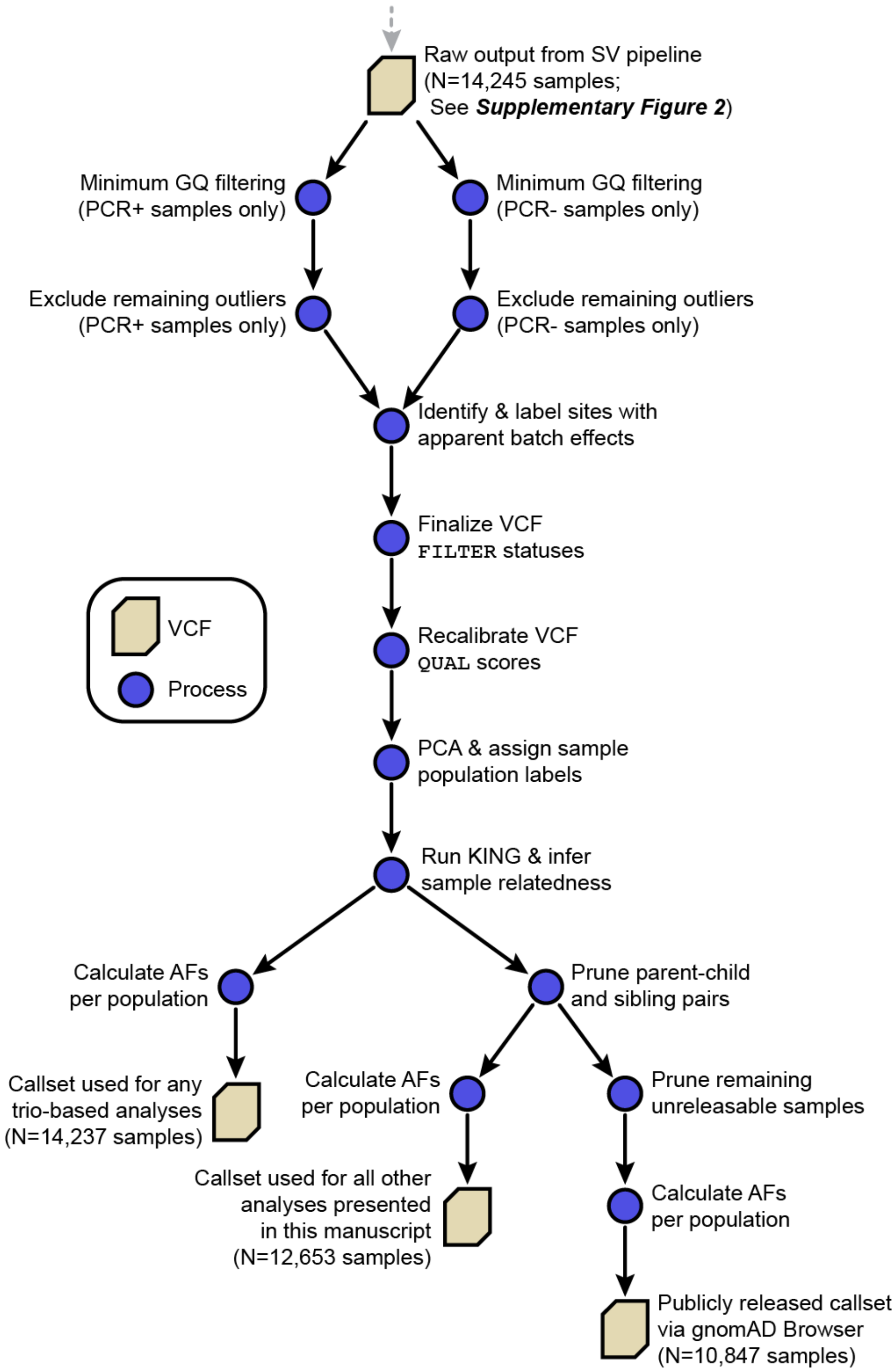


**Supplementary Figure 23 | Overview of *post hoc* filtering & final callsets.** As described in **Methods**, we performed a series of post hoc filters and post-processing steps to clean the final callsets used in the analyses for this study. These steps involved excluding outlier samples, detecting lingering batch effects, inferring relatedness, assigning population labels, and calculating allele frequencies.

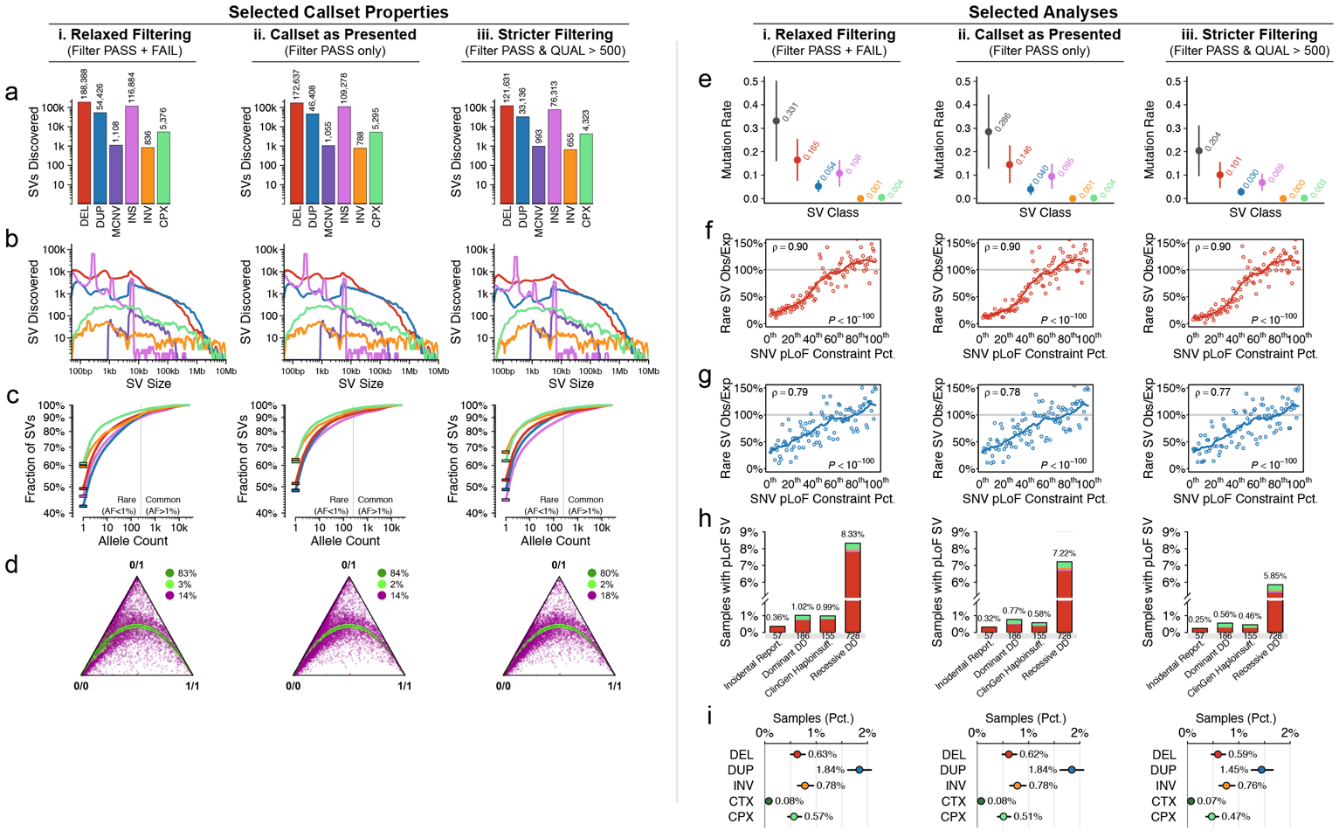


**Supplementary Figure 24 | Evaluation of callset filtering on key results.** We evaluated whether a variety of the principal findings in this study were sensitive to the callset filtering thresholds employed here. To examine this possibility, we reanalyzed the gnomAD-SV dataset at three quality thresholds, representing (**i**) relaxed filtering, where we included all SVs, even those that did not have a FILTER status of PASS, (**ii**) the same filtering thresholds as presented in this study, and (**iii**) stricter filtering, where we required all variants to have a QUAL score > 500 in addition to a FILTER status of PASS. For these three filtering thresholds, we assessed several callset properties, including (**a**) SV counts, (**b**) size distributions (see **Figure 1f**), (**c**) AF distributions (see **Figure 1g**), and (**d**) Hardy-Weinberg equilibrium rates (see **Extended Data Figure 2b**). Additionally, we reproduced multiple analyses presented in this study from these three callsets, including (**e**) mutation rate estimates (see **Figure 3a**), correlations between gene constraint and (**f**) pLoF SVs and (**g**) CG SVs (see **Figure 4d**), (**h**) carrier rates for rare pLoF SVs in medically relevant genes (see **Figure 6c**), and (**i**) carrier rates for large (≥1Mb) rare SVs (see **Figure 6d**). Across all analyses, we found that none of the principal conclusions of this study would have been meaningfully altered with either looser or stricter filtering, suggesting the findings as presented in this study are largely robust to the technical details of the gnomAD-SV dataset.

### SUPPLEMENTARY TABLES

#### Supplementary Table 1 | Sample QC thresholds & filtering

| | All Samples | | | PCR+ samples | | | | PCR- samples | | | |
|---|---|---|---|---|---|---|---|---|---|---|---|
| **Analysis stage** | Eligible samples | Sample failures | Fail rate | Threshold | Eligible samples | Sample failures | Fail rate | Threshold | Eligible samples | Sample failures | Fail rate |
| *Filter phase I: before SV discovery* | | | | | | | | | | | |
| **All phase I filters** | **14,891** | **513** | **3.45%** | | **1,631** | **228** | **13.98%** | | **13,260** | **285** | **2.15%** |
| Median absolute outlier bins Z-score | 14,891 | 215 | 1.44% | <0.6 | 1,631 | 109 | 6.68% | <0.4 | 13,260 | 106 | 0.80% |
| Median 100bp bin coverage | 14,891 | 141 | 0.95% | >30 & <100 | 1,631 | 104 | 6.38% | >30 & <100 | 13,260 | 37 | 0.28% |
| Dosage score (∂) | 14,891 | 86 | 0.58% | >-0.003 & <0.444 | 1,631 | 17 | 1.04% | >-0.175 & <0.092 | 13,260 | 69 | 0.52% |
| Autosomal ploidy spread | 14,891 | 79 | 0.53% | <1 | 1,631 | 51 | 3.13% | <0.8 | 13,260 | 28 | 0.21% |
| Pairwise alignment rate | 14,874 | 70 | 0.47% | >0.95 | 1,631 | 2 | 0.12% | >0.95 | 13,243 | 68 | 0.51% |
| Library contamination | 9,143 | 42 | 0.46% | <0.07 | 160 | 2 | 1.25% | <0.04 | 8,983 | 40 | 0.45% |
| Chimera rate | 14,874 | 39 | 0.26% | <0.02 | 1,631 | 8 | 0.49% | <0.025 | 13,243 | 31 | 0.23% |
| Read length | 14,874 | 10 | 0.07% | >125 | 1,631 | 10 | 0.61% | >125 | 13,243 | 0 | 0.00% |
| Ambiguous sex genotypes | 14,891 | 8 | 0.05% | . | 1,631 | 0 | 0.00% | . | 13,260 | 8 | 0.06% |
| Discordant reported & inferred sex | 14,863 | 5 | 0.03% | . | 1,631 | 0 | 0.00% | . | 13,232 | 5 | 0.04% |
| *Filter phase II: during SV discovery* | | | | | | | | | | | |
| **All phase II filters** | **14,378** | **141** | **0.98%** | | **1,403** | **69** | **4.92%** | | **12,975** | **72** | **0.55%** |
| SV call count outliers after module 03 | 14,378 | 133 | 0.93% | < Q3 + 6*IQR & > Q1 - 6*IQR | 1,403 | 61 | 4.35% | < Q3 + 6*IQR & > Q1 - 6*IQR | 12,975 | 72 | 0.55% |
| SV call count outliers after minimum GQ filtering | 14,245 | 8 | 0.06% | < Q3 + 6*IQR & > Q1 - 6*IQR | 1,342 | 8 | 0.60% | < Q3 + 6*IQR & > Q1 - 6*IQR | 12,903 | 0 | 0.00% |
| *Filter phase III: after SV discovery* | | | | | | | | | | | |
| **All phase III filters** | **14,237** | **1,584** | **11.13%** | | **1,334** | **279** | **20.91%** | | **12,903** | **1,305** | **10.11%** |
| Prune first-degree relatives | 14,237 | 1,584 | 11.13% | . | 1,334 | 279 | 20.91% | . | 12,903 | 1,305 | 10.11% |
| *Final analysis cohort* | | | | | | | | | | | |
| **Final analysis cohort** | **12,653** | **-** | **-** | | **1,055** | **-** | **-** | | **11,598** | **-** | **-** |

#### Supplementary Table 2 | Sample overlap with gnomAD SNV/indel analyses

| | All Samples in gnomAD-SV | | | Samples in Karczewski et al., 2019 | | | Samples not in Karczewski et al., 2019 | | |
|---|---|---|---|---|---|---|---|---|---|
| **Analysis stage** | All | PCR+ | PCR- | All | PCR+ | PCR- | All | PCR+ | PCR- |
| Initial cohort | 14,891 | 1,631 | 13,260 | 8,540 | 1,471 | 7,069 | 6,351 | 160 | 6,191 |
| Passed phase I filters (before SV discovery) | 14,378 | 1,403 | 12,975 | 8,176 | 1,246 | 6,930 | 6,202 | 157 | 6,045 |
| Passed phase II filters (during SV discovery) | 14,237 | 1,334 | 12,903 | 8,040 | 1,179 | 6,861 | 6,197 | 155 | 6,042 |
| Passed phase III filters (after SV discovery) | 12,653 | 1,055 | 11,598 | 7,626 | 979 | 6,647 | 5,027 | 76 | 4,951 |
| Included in public VCF release | 10,847 | 540 | 10,307 | 6,842 | 540 | 6,302 | 4,005 | 0 | 4,005 |

*Please refer to Supplementary Table 1 and Supplementary Figure 23 for a description of sample filtering phases*

#### Supplementary Table 3 | SV callset summary

| SV Class | Total SVs Discovered | Filter Pass Rate | SVs Used in Final Analyses* | Singleton (AC=1) | Rare (AC>1 & AF<1%) | Common (AF>1%) | Median Size (bp) |
|---|---|---|---|---|---|---|---|
| ALL | 433,371 | 78.7% | 335,470 | 166,937 | 142,744 | 24,734 | 331 |
| DEL | 191,527 | 91.6% | 172,637 | 88,331 | 72,017 | 12,289 | 756 |
| DUP | 55,380 | 85.4% | 46,408 | 21,995 | 22,118 | 2,295 | 4,283 |
| MCNV | 1,108 | 95.2% | 1,055 | 0 | 0 | 1,055 | 8,878 |
| INS | 118,700 | 93.5% | 109,278 | 52,860 | 46,469 | 9,949 | 280 |
| INV | 850 | 94.4% | 788 | 495 | 259 | 34 | 1,418 |
| CPX | 5,450 | 98.5% | 5,295 | 3,248 | 1,880 | 167 | 2,229 |
| CTX | 10 | 100.0% | 9 | 8 | 1 | 0 | NA |
| BND | 60,346 | 0.0% | 0 | 0 | 0 | 0 | NA |

**Final analyses restricted to filter-passing SV found in at least one unrelated individual (N=12,653 samples)*

#### Supplementary Table 4. | SV callset benchmarking

| Analysis | Details | Samples | SVs | Measurement | Value |
|---|---|---|---|---|---|
| **1. Trio analysis** | Rate of Mendelian violations per trio for autosomal SVs with complete trio genotypes at at least one non-reference allele present in the trio | 2,910 (970 trios) | 8,512 per trio (median) | Mendelian violation rate | 3.8% |
| | Rate of apparently *de novo* heterozygous autosomal SVs in children with complete trio genotypes | 2,910 (970 trios) | 4,686 per trio (median) | Heterozygous genotype error rate (mix of FDR in children, FNR in parents, and true *de novo* SV) | 3.0% |
| | Fraction of homozygous genotypes in children where at least one parent is reference for autosomal SVs with complete trio genotypes | 2,910 (970 trios) | 1,227 per trio (median) | Genotype error rate (mix of homozygous FDR in parents & heterozygous FNR in children) | 7.5% |
| | Number of untransmitted homozygous genotypes in parents divided by the sum of transmitted heterozygous genotypes in children and untransmitted homozygous genotypes for autosomal SVs with complete trio genotypes | 2,910 (970 trios) | 4,624 per trio (median) | Heterozygous genotype FNR | 1.9% |
| **2. CMA comparison** | Fraction of autosomal CNVs >40kb from CMA (Sanders *et al.,* 2015) with <30% coverage by simple repeats, segmental duplications, or somatic hypermutable sites that also have matching CNVs (≥50% coverage) in at least 50% of gnomAD-SV samples | 1,893 | 2,524 | Sensitivity (for large CNVs) | 97.1% |
| **3. Hardy-Weinberg equilibrium** | Fraction of autosomal biallelic SVs in HWE | 12,653 | 321,140 | HWE rate | 85.8% |
| **4. SV & SNV/indel linkage disequilibrium** | Median maximum genotypic correlation coefficient between common (AF≥1%) SVs with <30% coverage by simple repeats and segmental duplications and all SNVs/indels within ±1Mb from a subset of 5,353 overlapping AFR and EUR samples in this study and Karczewski et al. (2019) | 5,353 | 23,597 | Pearson Correlation Coefficient ($R^2$) | 0.85 |
| **5. Doubleton genotype analysis** | Fraction of doubleton (i.e., AC=2) SVs with ≤10% coverage by simple repeats and segmental duplications that also appear in two samples from the same population among all doubleton SVs appearing in any two samples (excluding 129 samples with uncertain population assignments) | 12,524 | 32,044 | Fraction of intra-population concordant doubleton SVs | 79.0% |
| **6. Comparisons to 1000 Genomes Project** | Correlation of AFs for biallelic autosomal SVs appearing at AF≥1% in either gnomAD and/or 1000 Genomes Project (Sudmant et al., 2015) | N/A | 37,907 | $R^2$ | 0.72 |
| **7. Long-read WGS comparison** | Fraction of SVs with SR support and <30% coverage by simple repeats and segmental duplications that also have long-read WGS support as computationally evaluated by VaPoR (Zhao *et al.*, 2017) | 4 | 4,829 per sample (mean) | PPV | 94.0% |

FDR = false discovery rate; FNR = false negative rate; CMA = chromosomal microarray; HWE = Hardy-Weinberg equilibrium; AC = allele count

#### Supplementary Table 5 | Common SVs in strong LD with SNV/indels

*Table too large to be reproduced here; provided separately as supplementary file*

#### Supplementary Table 6 | Carrier frequencies of genomic disorder CNVs in gnomAD-SV

| | Start | End | | | UK BioBank | | | gnomAD-SV | | |
|---|---|---|---|---|---|---|---|---|---|---|
| Chr | (Mb) | (Mb) | Genomic Disorder | CNV | *Samples* | *Carriers* | *Freq.* | *Samples* | *Carriers* | *Freq.* |
| 1 | 145.39 | 145.63 | TARdel | DEL | 396,725 | 69 | 0.017% | 10,001 | 2 | 0.020% |
| 1 | 145.39 | 145.63 | TARdup | DUP | 396,725 | 408 | 0.103% | 10,154 | 7 | 0.069% |
| 1 | 146.53 | 147.39 | 1q21.1del | DEL | 396,725 | 106 | 0.027% | 10,164 | 5 | 0.049% |
| 1 | 146.53 | 147.39 | 1q21.1dup | DUP | 396,725 | 168 | 0.042% | 10,164 | 10 | 0.098% |
| 2 | 96.74 | 97.68 | 2q11.2del | DEL | 396,725 | 29 | 0.007% | 10,164 | 1 | 0.010% |
| 2 | 96.74 | 97.68 | 2q11.2dup | DUP | 396,725 | 26 | 0.007% | 10,122 | 1 | 0.010% |
| 2 | 110.86 | 110.98 | 2q13delNPHP1 | DEL | 396,725 | 2,330 | 0.587% | 10,164 | 41 | 0.403% |
| 2 | 110.86 | 110.98 | 2q13dupNPHP1 | DUP | 396,725 | 1,868 | 0.471% | 10,164 | 118 | 1.161% |
| 2 | 111.39 | 112.01 | 2q13del | DEL | 396,725 | 51 | 0.013% | 10,162 | 1 | 0.010% |
| 2 | 111.39 | 112.01 | 2q13dup | DUP | 396,725 | 71 | 0.018% | 10,164 | 3 | 0.030% |
| 2 | 131.48 | 131.93 | 2q21.1del | DEL | 396,725 | 40 | 0.010% | 10,164 | 1 | 0.010% |
| 2 | 131.48 | 131.93 | 2q21.1dup | DUP | 396,725 | 55 | 0.014% | 10,164 | 5 | 0.049% |
| 3 | 195.72 | 197.35 | 3q29del | DEL | 396,725 | 9 | 0.002% | 10,164 | 0 | 0.000% |
| 3 | 195.72 | 197.35 | 3q29dup | DUP | 396,725 | 5 | 0.001% | 10,164 | 0 | 0.000% |
| 7 | 72.74 | 74.14 | WBSdup | DUP | 396,725 | 13 | 0.003% | 10,164 | 1 | 0.010% |
| 7 | 75.14 | 76.06 | 7q11.23dupdistal | DUP | 396,725 | 23 | 0.006% | 10,164 | 0 | 0.000% |
| 8 | 8.10 | 11.87 | 8p23.1dup | DUP | 396,725 | 5 | 0.001% | 10,164 | 0 | 0.000% |
| 10 | 49.39 | 51.06 | 10q11.21q11.23del | DEL | 396,725 | 56 | 0.014% | 10,164 | 0 | 0.000% |
| 10 | 49.39 | 51.06 | 10q11.21q11.23dup | DUP | 396,725 | 40 | 0.010% | 10,164 | 1 | 0.010% |
| 10 | 82.05 | 88.93 | 10q23dup | DUP | 396,725 | 7 | 0.002% | 10,164 | 0 | 0.000% |
| 13 | 20.98 | 21.10 | 13q12delCRYL1 | DEL | 396,725 | 363 | 0.091% | 10,164 | 5 | 0.049% |
| 13 | 20.98 | 21.10 | 13q12dupCRYL1 | DUP | 396,725 | 10 | 0.003% | 10,164 | 0 | 0.000% |
| 13 | 23.56 | 24.88 | 13q12.12del | DEL | 396,725 | 84 | 0.021% | 10,164 | 0 | 0.000% |
| 13 | 23.56 | 24.88 | 13q12.12dup | DUP | 396,725 | 212 | 0.053% | 10,163 | 7 | 0.069% |
| 15 | 23.68 | 28.39 | PWSdup | DUP | 396,725 | 16 | 0.004% | 10,164 | 0 | 0.000% |
| 15 | 29.16 | 30.38 | 15q11q13delBP3BP4 | DEL | 396,725 | 14 | 0.004% | 10,164 | 1 | 0.010% |
| 15 | 29.16 | 30.38 | 15q11q13dupBP3BP4 | DUP | 396,725 | 50 | 0.013% | 10,164 | 4 | 0.039% |
| 15 | 29.16 | 32.46 | 15q11q13dupBP3BP5 | DUP | 396,725 | 9 | 0.002% | 10,164 | 0 | 0.000% |
| 15 | 31.08 | 32.46 | 15q13.3del | DEL | 396,725 | 37 | 0.009% | 10,164 | 0 | 0.000% |
| 15 | 31.08 | 32.46 | 15q13.3dup | DUP | 396,725 | 224 | 0.056% | 10,164 | 2 | 0.020% |
| 15 | 32.02 | 32.45 | 15q13.3delCHRNA7 | DEL | 396,725 | 10 | 0.003% | 10,164 | 3 | 0.030% |
| 15 | 32.02 | 32.45 | 15q13.3dupCHRNA7 | DUP | 396,725 | 2,843 | 0.717% | 10,161 | 55 | 0.541% |
| 15 | 72.90 | 78.15 | 15q24dup | DUP | 396,725 | 8 | 0.002% | 10,164 | 0 | 0.000% |
| 16 | 15.51 | 16.29 | 16p13.11del | DEL | 396,725 | 124 | 0.031% | 10,164 | 1 | 0.010% |
| 16 | 15.51 | 16.29 | 16p13.11dup | DUP | 396,725 | 783 | 0.197% | 10,164 | 37 | 0.364% |
| 16 | 21.95 | 22.43 | 16p12.1del | DEL | 396,725 | 235 | 0.059% | 10,164 | 3 | 0.030% |
| 16 | 21.95 | 22.43 | 16p12.1dup | DUP | 396,725 | 192 | 0.048% | 10,164 | 8 | 0.079% |
| 16 | 28.82 | 29.05 | 16p11.2distaldel | DEL | 396,725 | 54 | 0.014% | 9,691 | 5 | 0.052% |
| 16 | 28.82 | 29.05 | 16p11.2distaldup | DUP | 396,725 | 127 | 0.032% | 10,164 | 2 | 0.020% |
| 16 | 29.65 | 30.20 | 16p11.2del | DEL | 396,725 | 103 | 0.026% | 9,548 | 2 | 0.021% |
| 16 | 29.65 | 30.20 | 16p11.2dup | DUP | 396,725 | 131 | 0.033% | 10,164 | 0 | 0.000% |
| 17 | 14.14 | 15.43 | 17p12delHNPP | DEL | 396,725 | 219 | 0.055% | 10,119 | 4 | 0.040% |
| 17 | 14.14 | 15.43 | 17p12dupCMT1A | DUP | 396,725 | 116 | 0.029% | 10,164 | 3 | 0.030% |
| 17 | 16.81 | 20.21 | PotockiLupski | DUP | 396,725 | 5 | 0.001% | 10,164 | 0 | 0.000% |
| 17 | 29.12 | 30.27 | 17q11.2delNF1 | DEL | 396,725 | 9 | 0.002% | 10,164 | 0 | 0.000% |
| 17 | 34.81 | 36.22 | 17q12del | DEL | 396,725 | 7 | 0.002% | 10,095 | 1 | 0.010% |
| 17 | 34.81 | 36.22 | 17q12dup | DUP | 396,725 | 99 | 0.025% | 10,164 | 1 | 0.010% |
| 22 | 19.04 | 21.47 | 22q11.2del | DEL | 396,725 | 10 | 0.003% | 10,164 | 0 | 0.000% |
| 22 | 19.04 | 21.47 | 22q11.2dup | DUP | 396,725 | 266 | 0.067% | 10,164 | 0 | 0.000% |

Genomic disorder coordinates & UK BioBank frequencies obtained from Owen et al., 2018.[11]

### SUPPLEMENTARY NOTES

**Technical benchmarking and quality assessment of the gnomAD-SV callset**

After SV discovery in 14,237 samples as described in the Supplementary Methods, we next aimed to comprehensively assess the performance of our SV discovery pipeline and the technical qualities of the gnomAD-SV callset. Benchmarking SVs from short-read WGS remains a major challenge with no gold standard.[12] As such, we evaluated the technical qualities of gnomAD-SV using seven orthogonal approaches, enumerated briefly here but also detailed in **Extended Data Figures 2-3**, **Supplementary Figures 6-12**, and **Supplementary Tables 4-5**. First, we assessed Mendelian inheritance in 970 parent-child trios (2,910 genomes), finding an average Mendelian violation rate of 3.8% and an apparent heterozygous *de novo* rate of 3.0% per trio. Given that almost all SVs that violate Mendelian transmission patterns represent a combination of false positives and/or negatives, these estimates provide proxies for SV detection performance and genotyping accuracy. Second, we found 97.1% sensitivity to detect large CNVs (>40 kb) previously reported from microarrays in 1,893 individuals.[13] Third, we calculated that 86% of SVs across all populations were in Hardy-Weinberg Equilibrium. Fourth, we found that most common (AF≥1%) SVs were in linkage disequilibrium (LD) with a nearby SNV or indel (median best $R^2$=0.85). Fifth, we observed that doubleton SVs overwhelmingly appeared isolated to specific populations, as expected (79.0% of doubletons were intra-population vs. 35.0% expected by chance; $P<10^{-100}$, one-sided binomial test). Sixth, most (57%) SVs documented in the 1000 Genomes Project were also found in gnomAD-SV, and the AFs of variants overlapping between the 1000 Genomes Project and gnomAD-SV were well correlated ($R^2$=0.72);[7] conversely, 86% of SVs in gnomAD-SV were novel compared to the 1000 Genomes Project, reflecting the ~7-fold increase in size, ~5-fold increase in average coverage, and improved sensitivity of gnomAD-SV. Seventh, we used long-read WGS data available for four individuals to perform an *in silico* confirmation of SVs predicted from short-read WGS in gnomAD-SV,[8,9,14] estimating a positive predictive value of 94.0% for SV not with breakpoint-level read evidence (92.8% of all SVs). We also evaluated breakpoint accuracy among the SVs with long-read WGS support by comparing the coordinates reported in gnomAD-SV to preexisting SV calls generated directly from the long-read WGS data,[8,14] finding that 59.8% of gnomAD-SV breakpoint coordinates were accurate to within a single nucleotide and 75.9% were accurate to within ±10bp. Importantly, these estimates assume the long-read WGS SV calls represent absolute ground truth, which will not hold for 100% of SVs (*e.g.*, spurious long-read misassembly, breakpoints with high homopolymer content, etc.). In conclusion, while the seven analyses listed above are imperfect measures of technical performance given the potential for confounding population substructure, admixture, recombination, recurrence, and other assumptions that will not universally hold true, they nevertheless establish the gnomAD-SV dataset as sufficiently sensitive and specific for most applications as a resource in contemporary human genomics.

### METHODS

#### WGS data aggregation

We processed a subset of the WGS data collected from population genetics and common disease genomics sequencing projects as part of the Genome Aggregation Database (gnomAD; https://gnomad.broadinstitute.org). Details of sample collection are provided in Karczewski *et al.*, 2019. Due to the availability of WGS BAM files at the time of SV callset generation, 8,540 genomes in this study overlap with those included in the gnomAD SNV and indel callset generation, and are described in Karczewski *et al.*, 2019. In addition, we included 6,351 genomes from other studies, either for the purposes of quality control or for population-based analyses.[1,15] These 6,351 additional genomes were collected primarily from three sources: (1) a subset of genomes (n=4,266) from the Multi-Ethnic Study of Atherosclerosis (MESA) cohort in the Trans-Omics for Precision Medicine (TOPMed) initiative, which has already been analyzed for common and rare variation;[15-18] (2) a subset of genomes we had previously analyzed and published from the Simons Simplex Collection (SSC; n=2,076 genomes), which were included for family-based quality control and benchmarking analyses, disease association and population screening, but were not consented for public release of site-frequency data;[1,13] and (3) nine genomes from the Human Genome Structural Variation (HGSV) consortium.[19] Note that we excluded all affected individuals from the SSC cohort prior to all analyses presented herein, with the exception of callset benchmarking. All variant and individual-level data from the SSC can be accessed by qualified researchers in SFARIbase (http://base.sfari.org; see *Data availability*). The nine HGSV samples were sequenced to very deep (~75X) coverage, but were downsampled to ~30X prior to being included in this study. **Supplementary Table 2** provides an explicit comparison to the WGS data also included in the gnomAD SNV and indel analyses.[6] We jointly processed and analyzed these 14,891 genomes, with the public release of genetic site-frequency data provided for 10,847 samples with appropriate consent, and the remaining samples released to appropriate repositories (see **Supplementary Tables 1-2** and **Supplementary Figure 23**).

#### Computational platform

Most WGS processing, SV discovery, and downstream analyses for gnomAD-SV was conducted on the FireCloud platform (https://software.broadinstitute.org/firecloud/), recently renamed to "Terra" (https://terra.bio/), which is a secure open platform for collaborative genome analysis developed as part of the NCI Cloud Pilot program.[2] Where relevant, all workflows and methods used in this study have been made publicly available via the FireCloud Methods Repository (https://portal.firecloud.org/#methods).

#### SV discovery

We performed discovery of SVs using an extension of a previously described modular, multi-algorithm integrative pipeline,[1] as integration of multiple independent algorithms has been shown to be an effective approach for SV discovery with balanced sensitivity and specificity.[1,19] The gnomAD-SV discovery pipeline is segmented into eight sequential modules, an overview of which is depicted in **Supplementary Figure 3**. Each module is described in detail below.

*Module 00: Preprocessing*

The first module of the gnomAD-SV pipeline collects all data and metadata required for SV discovery during the subsequent seven modules. This process involves five steps: (1) ploidy estimation and sex inference, (2) sample QC, including sequencing dosage bias scoring, (3) sample batching, and (4) execution of SV discovery algorithms. These steps are described below:

*Ploidy estimation & sex inference*

We estimated per-chromosome ploidy (*i.e.*, whole-chromosome copy number) and inferred genetic sex per sample using read depth in 1Mb sequential bins, excluding any bins where >5% of samples had zero observed coverage (*e.g.*, N-masked heterochromatic regions). We next normalized coverage values for each sample by dividing the total coverage for each 1Mb bin by the median coverage value across all autosomal 1Mb bins. We assigned ploidy per chromosome per sample as two times the median normalized coverage per 1Mb bin (**Extended Data Figure 1)**. For sex assignments, we rounded sex chromosome ploidy to the nearest integer copy state. Samples with predicted sex aneuploidies (not XX or XY) were assigned as "other". Finally, we screened for particularly large unbalanced rearrangements, such as somatic or mosaic aneuploidy and extremely large CNVs by assigning Z-scores and corresponding Benjamini-Hochberg (FDR) corrected q-values per 1Mb bin sample corresponding to the divergence of that sample's estimated copy number compared to the rest of the samples in the dataset (**Extended Data Figure 1b-e**).

*Sequencing dosage bias scoring*

We have previously observed that CNV calling from WGS can be confounded in samples with highly non-uniform coverage, which we here term "dosage bias", and that these dosage biases are antipodal between PCR+ and PCR- protocols (**Supplementary Figure 4a**).[1,3,20] To control for dosage bias during SV discovery, we developed a model named Whole-Genome Dosage (*WGD*) to quantify the extent of bias per sample. The *WGD* model produces a single metric (∂) that summarizes the directionality and magnitude of bias per sample, which we used to inform sample QC and batching. In brief, we compute ∂ by measuring the weighted mean of normalized coverage values per sample across 3,202 autosomal 100bp bins throughout the genome. These bins were selected on the basis of three features: (1) they have a high likelihood of being copy number-invariant between samples (**Supplementary Figure 4b**),

(2) they can significantly discriminate between PCR+ and PCR- samples across multiple independent sequencing batches and centers (**Supplementary Figure 4c**), and (3) they are roughly representative of all 22 autosomes (**Supplementary Figure 4d**). We confirmed the selection and weighting of these 3,202 bins by comparing to an independent test set of gnomAD-SV samples (**Supplementary Figure 4e**), and provide these bins as a public resource for quality control in future WGS-based studies (https://github.com/RCollins13/WGD). As anticipated, this model was able to improve read depth-based CNV discovery by grouping samples with similar dosage bias profiles (**Supplementary Figure 4g-h**), and also identify outlier samples with extreme biases to be excluded during sample QC.

*Sample QC*

We assessed the WGS properties for all 14,891 samples prior to SV discovery to exclude samples likely to introduce excessive noise into downstream analyses and subsequently reduce the overall quality of the SV dataset. Based on a combined analysis of all available QC metadata, we applied filters to 10 features measured per sample (**Supplementary Figure 1**). Definitions for these features are provided below:

- *Median 100bp bin coverage*: median sequencing coverage, measured in 100bp bins.
- *Dosage bias score (∂)*: measurement of uniformity of coverage (described above).
- *Autosomal ploidy spread*: absolute difference between highest and lowest ploidy estimates for any two autosomes.
- *Z-score of outlier 1Mb bins*: median absolute Z-score of number of 1Mb bins per chromosome with normalized copy number estimates < 1.5 or > 2.5. Z-scores were calculated separately for PCR+ and PCR- samples.
- *Chimera rate*: chimeric read pairs as a percentage of total read pairs.
- *Pairwise alignment rate*: fraction of all read pairs where both reads per pair aligned successfully.
- *Library contamination*: the maximum value of either adapter contamination fraction or estimated sample contamination fraction.
- *Read length*: mean read length.
- *Ambiguous sex genotypes*: normalized copy number estimates for chromosomes X and Y; chromosome X and Y copy numbers were considered ambiguous if outside the interval (1.1, 1.9) and (0.1,0.9), respectively.
- *Discordant inferred and reported sex:* samples where inferred and reported sex designations disagree, given that the sample had binary (male/female) sex assignments for both inferred and reported sex.

For quantitative features, we assigned filter thresholds separately for PCR-amplified (PCR+) and PCR-free (PCR-) WGS library preparation protocols. Given that samples for this study were aggregated across sequencing projects, centers, and dates, 34.7% lacked information for at least one of the 10 filtered features, though 99.8% had at least 9/10 filtered features. Filter thresholds and number of samples excluded per filter are provided in **Supplementary Table 1**. Any sample failing at least one filter was excluded from all SV discovery and downstream analyses.

*Sample batching*

We designed a batching scheme to subdivide the full cohort into smaller sample sets for raw SV discovery, and the final resolved SVs per batch were subsequently merged and re-genotyped across all samples (**Supplementary Figure 5**). This procedure was designed to control for potential batch effects and confounders, to leverage the opportunity of increased cloud-based parallelization, to surmount early computational challenges of simultaneous SV discovery in tens of thousands of genomes, and to mitigate the risk of decreased SV breakpoint accuracy due to large-sample joint SV discovery (*i.e.*, "overclustering" of non-identical breakpoints across samples). This batching scheme was executed as follows: all samples passing all initial sample QC filters were first split by PCR status (PCR+/PCR-). Within each PCR status, samples were next split on chrX ploidy rounded to the nearest whole integer. Samples with ≥2 copies of chrX were assigned to one batch ("female"), and samples with <2 copies of chrX were assigned to another batch ("male"). Each of the four PCR-sex groups were further split into quartiles based on median 100bp binCov values, yielding a total of 16 smaller groups where all samples per batch were matched on sex, coverage, and PCR status. Next, within each of these 16 groups, we ranked all samples by $\partial$ and split them into smaller groups of ~100 samples each. In the interest of keeping a uniform number of total batches of males and females, we optimized the number of ~100 sample groups based on all male samples, then split female samples into an equal number of batches. As is detailed below, we performed read depth-based CNV discovery with cn.MOPS on these ~100 sample batches.[21] This step was necessary because the computational requirements for cn.MOPS at sub-kilobase resolution become intractable for sets of >150-200 samples on most available servers. Finally, we merged every two batches of ~100 male samples with their corresponding two batches of ~100 female samples while maintaining ordering corresponding to both coverage and $\partial$. This last step yielded batches of ~400 samples (~200 male & ~200 female), which were matched for PCR status, coverage, and dosage biases. We intentionally did not include sample ancestry, sequencing project, or sample-sample relatedness as covariates in our batching scheme. We reasoned that having entire batches comprised of single ancestry groups, sequencing projects, or related samples would introduce unwanted batch-to-batch variability and technical artifacts in our final SV callset, so we aimed for a random distribution of these variables across all batches.

*Execution of individual discovery algorithms for SVs*

We refined our previous SV discovery approach[1] to incorporate four algorithms: Manta,[22] DELLY,[23] MELT,[24] and cn.MOPS.[21] Collectively, these algorithms consider three primary raw signals present in WGS data that can be used for SV discovery, namely split reads (SR), anomalous paired-end (PE) reads, and read depth (RD).[25] Each algorithm was selected with a specific rationale based on previous analyses:[1,19] Manta had the best all-around single-algorithm performance among all PE/SR algorithms we evaluated, DELLY maximizes sensitivity for small and balanced SV when run with default parameters, MELT specifically captures mobile element insertions (MEIs) with high sensitivity, and cn.MOPS is a flexible RD-based algorithm designed for cohort-based analyses with high sensitivity for rare CNVs. All four algorithms were run on all 14,891 samples in the gnomAD-SV cohort as described below:

*Manta*

We executed Manta v1.0.3 in single-sample mode with default parameters for 7,075 samples on the FireCloud platform[2] and 5,740 samples on a local cluster of 6,700 CPUs maintained by The Broad Institute. We also retrieved existing Manta calls we had previously generated for 2,076 samples as described in a recent publication.[1]

*DELLY*

We executed DELLY v0.7.7 in single-sample mode for deletions, duplications, insertions, and inversions for 7,075 samples on FireCloud and DELLY v0.7.6 for 5,740 samples on the local Broad Institute cluster. Like Manta, we retrieved existing DELLY calls for 2,076 samples analyzed as part of an earlier study.[1]

*MELT*

We executed MELT v2.0.5 in single-sample mode for 7,075 samples on FireCloud and 5,740 samples on the Broad Institute cluster. As for Manta and DELLY, we retrieved existing MELT calls for 2,076 samples analyzed previously.[1]

*cn.MOPS*

We executed a custom implementation[1] of cn.MOPS v1.20.1 on FireCloud for all 14,891 samples in ~100-sample batches as generated during sample batching (see above). For each 100-sample batch, we composed coverage matrixes across all samples at 300bp and 1kb bin sizes per chromosome, excluded any samples with a median bin coverage of zero per contig, then ran cn.MOPS with R v3.3.3, split raw calls per sample, segregated calls into deletions (copy number < 2) and duplications (copy number > 2), merged the 300bp and 1kb resolution calls per sample per

CNV type using BEDTools merge, and subtracted any N-masked bases from all CNV calls using BEDTools subtract.[26]

After raw SV calls from all four algorithms were aggregated for each sample, we next standardized each VCF or BED file to match specifications expected by the downstream pipeline modules using *svtk standardize* (https://github.com/talkowski-lab/svtk).[1] We stripped all raw SV calls on chrX and chrY for samples with non-canonical inferred sexes from our ploidy estimation procedure.

During module 00, we also collected PE, SR, RD, and SNP B-allele frequency (BAF) evidence per sample. We collected discordant PE evidence and SR evidence using *svtk collect-pesr*, RD evidence using binCov, and BAF evidence from GATK HaplotypeCaller-generated VCFs using a custom script (vcf2baf) included in the gnomAD-SV pipeline codebase on FireCloud.[27] We were unable to obtain GATK VCFs for 0.2% (32/14,891) of samples, and thus lacked BAF data for these samples. Following evidence collection per sample, we constructed PE, SR, RD, and BAF matrices merged across all samples in each 400-sample batch.

All subsequent modules (modules 01-07) were executed in FireCloud unless otherwise specified.

*Module 01: Clustering*

The second module of the gnomAD-SV pipeline involves clustering of all variant calls per algorithm across all samples in each batch of samples. For each 400-sample batch (described above), we used *svtk vcfcluster* to cluster all calls for all samples per PE/SR algorithm (Manta, DELLY, and MELT) while requiring a maximum of 300bp distance between breakpoints and at least 10% reciprocal overlap by size. We excluded any variants whose breakpoints mapped within our PE/SR clustering blacklist, as previously described.[1] In parallel, we clustered cn.MOPS calls for all samples per batch using *svtk bedcluster* while requiring 80% reciprocal overlap by size and no constraints on breakpoint distance. For both PE/SR and RD calls, where two or more calls met the above clustering criteria, we collapsed all clustered calls into a single record using the median coordinates across all clustered variants. The output of module 01 was three VCFs and one BED file per 400-sample batch, corresponding to one file each for each of the four SV algorithms used (Manta, DELLY, MELT, and cn.MOPS).

*Module 02: Evidence Collection*

The third module of the gnomAD-SV pipeline involves querying four modes of raw evidence present in the original aligned WGS BAM files for all samples per batch for each SV call. While this process is described in extensive detail elsewhere,[1] we also briefly summarize it here. For each SV call, we collect the following information:

*PETest & SRTest (all SVs except RD-only CNVs)*

We assess the number of discordant read-pairs and split-reads per sample that supports the called SV, and require the orientation of reads per pair to match the expected signatures for the corresponding SV class.[25] The count of supporting discordant pairs or split-reads is tabulated per sample predicted to carry the SV, and also in a randomly selected background population of 160 samples predicted to not carry the SV. These counts are subsequently compared between predicted SV carriers and predicted non-carriers with a Poisson test to derive one P-value each for PE and SR evidence.

*RDTest (CNVs only)*

Like *PETest* and *SRTest*, we also assess the difference in RD between predicted CNV carriers and non-carriers. RDTest compares the median normalized coverage values between carriers and non-carriers with a two-sample t-test or a one-sample Z-test, depending on the number of predicted CNV carriers, and emits a P-value and a RD separation metric for each putative CNV.

*BAFTest (CNVs only)*

Finally, we also compare the normalized BAF for heterozygous SNVs within predicted CNVs between carriers and non-carriers. The distribution of BAFs is compared between groups of predicted carriers and non-carriers with a Kolmogorov-Smirnov test for duplications or a Gaussian mixture model for deletions, which both produce a P-value and test statistic for each CNV.

The output of module 02 is four matrices per batch, corresponding to one each for Manta, DELLY, MELT and cn.MOPS. Each matrix contains the test statistics and evidence for every SV call made by that algorithm in that batch, and this evidence matrix is fed directly into the random forest filtering step in module 03.

<u>*Module 03: Variant Filtering*</u>

The fourth module of the gnomAD-SV pipeline filters predicted SV calls per batch based on the strength of raw evidence supporting each call. This step is essential to exclude the overwhelming number of spurious false-positive SV calls emitted from each algorithm and retain a subset of SV enriched for true-positive SVs. We perform this filtering with a series of random forest classifiers, which have already been described in detail elsewhere.[1] In brief, this process uses the four modes of evidence produced in module 02 to assign each SV to one of three categories: predicted valid SV, predicted invalid SV, or uncertain. The predicted true SV and false SV are used for training in the random forest, which are then applied across all variants providing an estimated probability of being a valid SV. To correct for overfitting during random forest training we perform a series of ROC optimizations for all evidence metrics produced by module 02, after which we compute a joint probability that each SV is a true variant across all available forms of evidence (PE, SR, RD, BAF). Next, we permanently exclude all SVs with an integrated probability < 0.5. These variants are categorized as false-positive SV calls by the initial algorithms, and are not considered for any subsequent analyses. Finally, we apply a strict heuristic cutoff of ≥5kb in size for CNVs discovered from read depth-based analyses alone, as we have previously observed the false discovery rate for CNVs uniquely detected by read depth increases dramatically below this threshold.[1] We acknowledge that true mosaic and sub-integer copy-state SVs will likely also be filtered out at this stage, as they will exhibit suboptimal support despite being biologically valid SVs. Thus, we emphasize that the filtered SVs retained during module 03 are heavily biased towards germline SVs. Finally, we filtered samples from each batch that remained SV call count outliers even after random forest filtering of SV sites. To determine which samples were SV call count outliers, we counted the number of non-reference SV sites per SV type per algorithm for each sample per batch. Within each batch, we considered a sample to be an outlier if it was outside of six times the IQR for any SV type. Outlier samples were stripped from the cohort and excluded from all subsequent SV discovery and analyses (**Supplementary Table 1**).

*Module 04: Genotyping*

The fifth module of the gnomAD-SV pipeline assigns a genotype and quality score for each sample for every SV based on support from three forms of evidence (RD, PE, SR). Prior to genotyping, all nonredundant SVs discovered in any batch are collated to form a master set of all SVs across the full gnomAD-SV cohort, and each sample is genotyped for this master set of variants. This process is described below:

*RD genotyping (CNVs only)*

For each CNV, a median normalized RD value is calculated per sample by taking the median normalized RD value across all 100bp bins located within that CNV after excluding bins with mapping quality of zero, unless the removal of these unmappable bins would result in fewer than 10 eligible bins within the CNV. CNVs > 1Mb are restricted to the inner 1Mb as a proxy, consistent with the behavior of RdTest (see *Module 02*). RD genotyping thresholds are first trained on a set of 64 previously characterized multiallelic sites (available from: https://github.com/talkowski-lab/RdTest),[28] which exhibit tight normal distributions of normalized RD values centered at each integer copy state. After determining the expected distributions of normalized RD values for each copy state, we next assign a copy state for each sample at every CNV based on a Z-test against each copy state distribution. Samples are automatically assigned homozygous reference genotypes if they do not exceed the minimum RD separation threshold determined by the random forest stage of module 03. Finally, genotype quality (GQ) is assigned as a Phred score based on the P-value from the most likely copy state minus the Phred score for the second most likely copy state. GQ scores are capped at 999, similar to GATK.[27]

*PE/SR genotyping (all SVs except RD-only CNVs)*

For each SV, counts of discordant pairs and split reads supporting the SV are tallied per sample. Genotype assignment is carried out in two phases, as follows. First, a binary determination is reached for each sample as to whether or not that sample's genome carries the SV by comparing the PE or SR evidence in that sample to the cutoffs determined by the random forest step of module 03. Second, for samples predicted to carry each SV, a genotype is assigned based on PE or SR distributions matched to genotyped copy states for CNVs > 1kb determined during RD genotyping (see above). Similar to RD genotyping, both PE and SR counts for predicted SV carriers are normally distributed at each integer copy state, and therefore a similar GQ can also be assigned per sample. For predicted non-carrier samples, GQs are assigned according to a Poisson test, given that PE and SR counts for non-carrier samples do not match those for predicted SV carriers. GQ scores are capped at 999, similar to GATK.[27]

*Consensus genotype integration*

After PE, SR, and RD genotypes have been assigned to each sample for every SV, an integrated genotype is composed according to SV class and size. For each SV, one of the three evidence types (PE/SR/RD) is considered "primary", and the others are considered "secondary". The primary evidence is used to assign the overall genotype, and the secondary evidence provides a bonus to GQ scores if concordant with the primary evidence: if the other pieces of evidence support the primary, a bonus of $(999-GQ_{primary}) \times (0.5 \times GQ_{secondary} / 999)$ is added to the primary GQ. For CNVs > 5 kb, RD is primary and the better non-reference genotype between PE or SR is secondary. For CNVs between 1-5kb, the higher-quality non-reference genotype between PE or SR is primary and RD is secondary. For all other variants, the higher-quality non-reference genotype between PE or SR is primary, and the other is secondary. Once all samples are genotyped per SV, each variant is assigned a QUAL score based on the median GQ across all non-reference samples for that SV.

*Module 05: Batch Integration*

The sixth module of the gnomAD-SV pipeline involves the codification of genotyped SV calls across all batches in the cohort, merging of PE/SR and RD calls, and subsequent resolution of these merged SV calls into complete genomic variants. Components of this process have been described previously,[1] but this module also includes multiple new and modified processes. This occurs in four steps, listed below:

*Cross-batch call clustering*

The first step of module 05 clusters each genotyped VCF from module 04 across all batches. This is performed once for the three PE/SR algorithms and once for cn.MOPS. We first perform a column-wise join across all VCFs from each of the 36 batches, and subsequently run *svtk vcfcluster* on the new cohort-wide joined VCF to collapse overlapping variants.[1] When clustering, we require a minimum of 50% of samples with non-reference genotypes to overlap between records. For PE/SR algorithms, we additionally require a maximum breakpoint distance of ±300bp and a minimum reciprocal overlap of 10% by size, whereas for cn.MOPS, we required a maximum breakpoint distance of ±500kb and a minimum reciprocal overlap of 50% by size. For instances of two or more variants being clustered, each sample retains the non-reference genotype (if any) with the highest genotype quality score among all variants in the cluster. The output of this step is two clustered VCFs for the entire cohort: one containing all PE/SR-based SV calls, and one containing all RD-based CNV calls.

*PE/SR and RD call merging*

The second step of module 05 merges the cohort-wide PE/SR-based and RD-based SV calls output from the previous step. In this merging, we first construct a graph of all overlapping PE/SR and RD SV calls while requiring 50% reciprocal overlap by size, matching SV classes, and at least 50% overlap among samples with non-reference genotypes. Each cluster in this graph is collapsed into a single record, where the SV coordinates from the PE/SR record are retained but the union of non-reference sample genotypes are assigned as in module 05i. The output of this step is a single VCF containing all SV calls across the full cohort.

*Variant resolution*

The third step of module 05 examines predicted alternate allele structures from individual breakpoints to construct SV consisting of multiple breakpoints. This process is performed twice in parallel: once while including all SVs, and once while restricting to inversion breakpoints alone to capture large inversion-mediated complex SV. Variant resolution is performed with *svtk resolve,* the framework for which has been described at length in two previous publications.[1,3] For clarity, we also provide a brief description of this process here. In summary, *svtk resolve* first performs single-linkage clustering of all overlapping SV while requiring a maximum breakpoint distance of ±300bp and 50% overlap among samples with non-reference genotypes. It next compares the coordinates and SV classes of each cluster of SVs against a dictionary of known SV signatures, which resolves canonical translocational insertions, canonical inversions, canonical reciprocal translocations, and 11 complex SV subclasses (see **Figure 2**).[1,3] Non-CNV SVs involved in a multi-SV cluster that are unable to be resolved are marked as unresolved, and are converted to BNDs to accordingly. This entire process is performed two times, sequentially: first when requiring the relatively strict (±300bp) breakpoint distance to capture easily resolved SVs, then a second time while considering any remaining unresolved variants with a more relaxed breakpoint distance criteria of ≥2kb to capture complex SV with large (≥2kb) flanking CNVs. SVs that do not cluster with any other SV, or those that cannot possibly form a complex SV (*e.g.*, two partially overlapping deletions), are left unchanged. The last step of this process is to resolve discrepancies between the outputs of *svtk resolve* when run on all variants and when restricted to only inversions: if an SV is incorporated into a resolved SV in one output but not the other, we retain the resolved SV and discard the unresolved alternative. The output of this step is one VCF one containing all variants, including resolved canonical SV, resolved complex SV, and unresolved BNDs.

*Complex variant regenotyping*

The final step of module 05 is to confirm predicted complex SV structures via RD regenotyping of predicted CNV intervals. To accomplish this, we perform RD genotyping for all 36 batches for all predicted CNV intervals involved in candidate complex SVs with the same procedure as described in module 04, collect the copy state predictions across all samples from all 36 batches, and compare the ratio of samples with expected copy states (*i.e.*, copy state < 2 for a predicted complex deletion and copy state > 2 for a predicted complex duplication) between predicted carriers and non-carriers. For all CNVs > 1kb, we then compute the "confirmation rate" for predicted carrier and non-carrier samples as the fraction of samples with expected copy states divided by the total number of samples. We consider a CNV to be confirmed if the difference in confirmation rates between predicted carriers and non-carriers is at least 40% than for non-carriers (*e.g.*, at least 40% of carriers and 0% of non-carriers, or 90% of carriers and 50% of non-carriers). We restrict this comparison to only consider female samples on chromosome X and male samples on chromosome Y. CNVs ≤ 1kb are assessed for confirmation, but a failure to confirm small CNVs in this size range does not count as a regenotyping failure. Once all CNVs involved in a candidate complex SV are labeled based on this regenotyping procedure, we consider the entire complex SV as confirmed unless any CNVs fail to regenotype (or are <1kb, as described above). Candidate complex SV with at least one involved CNV labeled as a regenotyping failure are rejected and converted to unresolved BND variants.

Following the four steps above, the output from module 05 is a single cohort-wide genotyped VCF with resolved canonical SVs, resolved complex SVs, and unresolved SVs. This VCF is passed to the final VCF refinement step in module 06, described below.

*<u>Module 06: VCF Refinement</u>*

The seventh module of gnomAD-SV pipeline corrects inconsistencies in RD-based CNV genotyping that arise due to difficulties in predicting copy state for overlapping CNVs. Namely, the gnomAD-SV pipeline uses copy number as predicted by RD evidence as the primary source for assigning genotypes to CNVs > 5kb in size; however, in instances of overlapping CNVs, this approach can be confounded without deconvolving each haplotype by phase. To account for this, we apply a correction to CNVs > 5 kb that are not multiallelic (*i.e.*, more than three distinct copy states observed) as follows: per sample, we first isolate pairs of CNVs with at least 50% overlap by size, using BEDTools coverage.[26] The strength of evidence supporting each CNV is then assessed based on CNV size, where larger CNVs are considered to have stronger support, and type(s) of evidence with P ≥ 0.5 from the module 03 random forest (*e.g.*, RD, PE). For each pair, we then correct copy state and genotype for the CNV with weaker support. Concurrent with this overlapping CNV correction, we also explore nested compound heterozygous deletions and duplications, where one of the CNVs may have what appears to be a reference copy state due to the change in copy number being masked by the opposing CNV on the other allele. After correction of copy states, new genotypes are assigned for all samples, and a final multiallelic tag is assigned to CNVs > 5 kb with at least 1% of samples having copy states at least 2 deviations away from expectation (*e.g.*, a deletion call with a maximum copy number of four or more). CNVs tagged as multiallelic are relabeled as "MCNV". In addition to the overlapping CNV correction, this module also handles sex chromosome genotype correction, which is evaluated in a sex-aware manner. For those individuals with a predicted sex chromosome abnormality (*e.g.*, XXY; also see **Extended Data Figure 1**) genotypes are automatically assigned as null on sex chromosome.

*Module 07: Gene annotation*

The eighth and final module of the SV discovery pipeline annotates all SV against known protein-coding genes. We used protein-coding gene annotations from the Gencode v19 comprehensive annotation file.[29] Where multiple transcripts were available for a single gene, we restricted analyses to the transcript matching the Ensembl definition of canonical transcript (see https://useast.ensembl.org/Help/Glossary?id=346). UTRs were defined as the elements designated as UTRs in Gencode v19 that also corresponded to the Ensembl canonical transcript. Promoters were defined as the 1kb window directly preceding each gene body in Gencode v19 on the transcribed strand. We annotated each canonical SV for a range of possible predicted effects on coding sequences, as is graphically outlined in **Supplementary Figure 17** and described below:

- *Loss of function (pLoF)*: we predicted an SV to cause genic pLoF on a SV class-specific basis, as follows:
    - Deletions: any overlap with at least one exon.
    - Duplications: both breakpoints wholly contained within exons of the same gene.
    - Insertions: insertion of any sequence directly into an exon.
    - Inversions: any inversion where one breakpoint is contained within a gene (exon or intron) and the other breakpoint is outside of the same gene, or any inversion where both breakpoints are contained within the same gene and the inversion overlaps at least one exon from that gene.

    ○ Translocations: any translocation breakpoint that overlaps an exon or intron.

- *Copy gain (CG)*: we predicted an SV to cause a whole-gene CG if and only if the SV involved a duplicated segment that completely spanned an entire gene (defined as the first nucleotide of the first exon extending to the last nucleotide of the last exon from the canonical transcript).
- *Intragenic exonic duplication (IED)*: we predicted an SV to cause IED if and only if the SV involved a duplicated segment where both breakpoints were contained within the same gene, at least one exon was intronic, and the duplication overlapped at least one exon.
- *Partial gene duplications*: we predicted an SV to result in a partial gene duplication if the SV involved a duplicated segment where one breakpoint was contained within a gene (exon or intron) but the other breakpoint was found outside that same gene. The functional consequence of these rearrangements is unclear, and likely to infrequently result in altered gene function; thus, for most analyses, these partial gene duplications were not considered to be gene-altering.
- *Whole-gene inversion*: we predicted an SV to invert an entire gene if the SV involved an inverted segment that completely covered an entire gene, using the same definition as for CG annotations (see above). Given that we would not predict any direct alterations to coding sequence from whole-gene inversions, we did not consider these whole-gene inversions as gene-disruptive in our analyses, although we cannot rule out the possibility that a subset of these variants might have context-specific positional effects on gene regulation in *cis*.
- *Multiallelic exon overlap*: we noted all MCNVs that overlap at least one exon, but did not consider these SVs to categorically cause any one functional effect (per above). We did not count MCNVs towards any site-level analyses of genic effects, but instead evaluated the predicted effects of each MCNV on a per-sample basis according to each sample's predicted copy state (*i.e.*, genotype). Samples with a predicted copy state < 2 were treated as MCNV (pLoF), whereas samples with a predicted copy state > 2 were treated as MCNV (CG). We fully anticipate these MCNV designations are oversimplifying the true complexity of these MCNV haplotypes and their diploid arrangement; however, given the relative sparsity of MCNVs in the genome, and absent tedious manual curation, improved MCNV phasing methods, and/or other positional information, we used the generalization outlined here as a rough proxy for the genic effects of MCNVs.
- *UTR SVs*: we labeled SVs as UTR-disruptive if at least one breakpoint was contained within a gene's 5' or 3' UTR, but the gene did not meet any of the above criteria to otherwise be considered gene-disruptive.
- *Promoter SVs*: we labeled SVs as promoter-disruptive if at least one breakpoint was contained within a gene's promoter, but the gene did not meet any of the above criteria to otherwise be considered gene-disruptive.

- *Intronic SVs*: we labeled SVs as intronic if both breakpoints were contained within the same gene, but the SV did not meet any of the above criteria to otherwise be considered gene-disruptive (including promoter disruptions).
- *Intergenic SVs*: all SVs not meeting any of the above criteria were considered intergenic. For these SVs, we also noted the gene with the nearest TSS by linear distance.

Given their multiple interleaved distinct SV signatures, we treated complex SV separately from all canonical SV during gene annotation. For each complex SV, we first deconstructed the rearrangement into its component intervals (labeled as "CPX_INTERVALS" in the VCF INFO field), annotated each interval according to its SV class and coordinates, then composed a consensus annotation for the overall complex SV as the union of predicted effects from all of the component intervals.

The output from this annotation process in module 07, the final module in the gnomAD-SV discovery pipeline, is a genotyped VCF containing all SVs across all samples, with functional genic annotations assigned to each SV (as above).

**Sample and variant QC after SV discovery**

Following SV discovery, we performed a series of per-sample and per-variant QC steps and filters, in the order described below. These *post hoc* callset adjustments are also outlined in **Supplementary Figure 23**.

<u>*Optimizing per-sample genotype quality filters*</u>

We first aimed to control false positive genotypes per sample by applying a series of conditional filters to the genotype quality (GQ) statistic for each genotype at each SV site. To accomplish this, we considered the rate of apparently *de novo* SVs among the 1,173 parent-child trios present in our SV callset, as we reasoned that a large fraction of apparently *de novo* SVs would represent spurious false-positive genotypes in the child. Given that the SV genotyping procedure in module 04 relies on different combinations of evidence for different SV classes, we performed a GQ threshold optimization procedure separately for each PCR status (PCR+, n=203 trios; PCR-, n=970 trios) across six SV classes (DEL, DUP, INS, INV & CPX, BND, and all SV classes), four size ranges (<1kb, 1-5kb, ≥5kb, and all sizes), four allele frequency ranges (<1%, 1-10%, ≥10%, and all frequencies), five VCF filter statuses (PE/SR support for both sides of the breakpoint, high SR background rate, PESR genotyping overdispersion, everything else, and all filter statuses), and four per-sample genotype evidence categories (RD-only, SR-only, everything else, and all evidence categories), for a total of 1,448 distinct filter conditions tested for each PCR status after removing impossible combinations of filters (*e.g.*, RD-only balanced SVs).

For each filter condition, we executed a GQ threshold assignment procedure as follows: we first extracted genotypes and GQ metrics for every trio across all biallelic autosomal SVs where the child was genotyped as heterozygous and the SV matched the specified filter parameters. For trios where >1,000 SVs met the criteria for a given filter combination, we randomly downsampled to 1,000 SVs. We next titrated across a range of minimum GQ thresholds from 0 to 999 in increments of 10. At each candidate GQ threshold, we replaced all genotypes with GQ below this threshold with no-call genotypes (*i.e.* "./."), and computed two metrics: (1) the fraction of heterozygous SV retained in the proband that had appeared as inherited prior to minimum GQ filter application, and (2) the percentage of heterozygous SVs retained per child appearing *de novo* among sites where all three members of the trio still retained non-no-call genotypes. We then computed the median for each of these statistics across all trios at each candidate minimum GQ threshold and performed a receiver operating characteristic (ROC) analysis to find the optimal GQ. We constrained this ROC analysis to find the lowest GQ cutoff such that we could maximize percentage of inherited heterozygous SV retained while also satisfying a maximum tolerated apparent *de novo* rate of 5%.

After determining the optimal minimum GQ threshold for each of the 1,448 filter condition listed above, we discounted the results from any condition with a median number of heterozygous SVs per child less than 11. For these conditions with ≤10 heterozygous SVs per child, we adopted a minimum GQ threshold from a closely related condition that satisfied the minimum requirement of heterozygous SVs per child; this "closely related" condition was determined based on ascending a hierarchical tree of most-to-least effective filter combinations for each SV type, where filters were ranked on effectiveness by maximizing the fraction of inherited heterozygous SVs retained per child at the ROC-optimal minimum GQ threshold while also maintaining apparent *de novo* rate ≤5%.

Once all minimum GQ thresholds were determined for each filter condition for PCR+ and PCR- samples separately, we replaced all homozygous reference or heterozygous biallelic genotypes to no-call genotypes per SV for any sample with GQ below the corresponding threshold based on that sample's PCR status. We did not apply any GQ filtering to homozygous genotypes, multiallelic sites (MCNVs), or chromosomal translocations. SVs without any remaining non-reference genotypes after minimum GQ filtering were dropped from the callset. Finally, we identified SVs that suffered significant shifts in AF after filtering by comparing allele counts and allele numbers before and after minimum GQ filtering with a chi-squared test, which was performed separately for PCR+ and PCR- samples. SVs with (i) at least 2% of samples with null genotypes and (ii) a significant P-value for the difference of allele counts before and after filtering were labeled as having an unstable AF estimate. We considered a P-value as significant only after Bonferroni correction for the total number of SVs evaluated. Variants with unstable AF estimates in PCR+ samples had "UNSTABLE_AF_PCRPLUS" to the VCF info field and all PCR+ sample genotypes were assigned as null, while variants with unstable AF estimates in PCR- samples had "UNSTABLE_AF_PCRMINUS" added to the VCF filter field and all PCR- sample genotypes were assigned as null.

*Outlier sample exclusion*

After filtering genotypes on GQ (see above), we next evaluated whether any samples were outliers in terms of total number of SVs per genome. We counted the total number of non-reference autosomal biallelic SV observed per sample for each SV class with an average of more than 100 SVs per genome (DEL, DUP, INS, BND) after excluding SVs with the PESR genotype overdispersion VCF filter to protect against high rates of homozygous genotypes of these sites masking true outlier samples. We labeled samples as outliers if they had SV counts from any class that was either more than six times the inter-quartile range (IQR) more than the third quartile or less than first quartile across all samples for that SV class. We performed this process separately for PCR+ and PCR- samples. Outlier samples were pruned from the callset, and SVs without any remaining non-reference genotypes after outlier sample exclusion were also excluded outright from the callset.

*Assessment of batch effects*

We next assessed the concordance of SV calls between all pairs of the 36 batches used during SV discovery. We were particularly interested in identifying any SVs that may have been preferentially discovered in one or a subset of batches due to factors other than sex or ancestry, and thus may be unevenly represented across the full gnomAD-SV cohort and skew AF distributions. To accomplish this, we first computed batch-specific AF statistics for every variant for samples from each of four major populations (African/African-American, Asian, European, or Latino) based on unrefined preexisting sample labels corresponding to ancestry inferred from SNV analyses on the same samples (where available) or self-reported race or ethnicity (only when necessary).[6] For MCNVs, we computed AF as the total count of non-diploid individuals divided by the total number of individuals genotyped at that site. We excluded all children from known parent-child trios from all batches when calculating AF to improve the accuracy of AF estimates. For each nonredundant pair of batches (n=630 pairs), we restricted to sites where at least one non-reference allele was observed in either batch and at least one population had at least 60 non-null (*i.e.*, genotyped) alleles in both batches, or 30 individuals with non-null genotypes for MCNVs. We controlled for differences due to ancestry by restricting to the same population on a per-variant basis, and further maximized the accuracy of these comparisons by restricting to the optimal population separately for each variant, where "optimal" was defined as the population with the largest minimum number of non-null alleles between both batches that also met the criteria above (≥60 non-null alleles in both batches & ≥1 non-reference allele in either batch). For each

SV, we tabulated the observed AF in the selected optimal population for both batches, and assessed the significance of any differences in AF between the two batches with a chi-squared test, and performed a Bonferroni correction on the resulting chi-squared P-values to control for the many thousands of sites being compared between any two batches. In parallel, we performed an identical analysis for batch-specific variants, where we compared the AFs for all sites observed in a single batch against the AF of those sites summed across all other 35 batches. We considered a variant to have evidence of batch effects if it had a Bonferroni-corrected P-value < 0.05 in at least 12/630 possible pairwise comparisons or any of the 36 batch-specific variant comparisons. For each variant with batch effects, we subsequently determined whether the batch effect was being driven predominantly by PCR+ or PCR- samples by calculating the fraction of batch-batch pairs with significant batch effects that involved a PCR+ batch. Since 4/36 batches (~11%) were PCR+, we used 11% as a cutoff to discriminate between PCR+ and PCR- batch effects. SVs with significant batch effects were handled as follows:

- If at least 11% of failed comparisons involved a PCR+ batch, and the average AF was higher in PCR+ batches than in PCR- batches, that variant was marked with a “PCRPLUS_ENRICHED” tag in the VCF filter column.
- If at least 11of failed comparisons involved a PCR+ batch and the average AF was lower in PCR+ batches than in PCR- batches, all genotypes from PCR+ samples were rewritten as no-calls and a “PCRPLUS_DEPLETED” tag was added to the VCF INFO field, but no new VCF filter status was assigned. Sites with zero non-reference alleles remaining after excluding PCR+ samples were dropped from the callset.
- If less than 11% of failed comparisons involved a PCR+ batch, the variant was marked with a “VARIABLE_ACROSS_BATCHES” tag in the VCF info field.

*Assignment of final VCF filter labels*

Despite efforts to balance sensitivity and specificity throughout SV discovery and genotyping in this cohort, we nevertheless wanted to categorically partition the final SV callset into a high quality (*i.e.* analysis-ready) subset and a second subset corresponding to variants of lower quality. In particular, unresolved variants like BNDs can dramatically inflate variant counts both cohort-wide and per-sample, but do not have resolved structures and thus are largely uninterpretable for downstream analyses. We also noticed an enrichment of apparent false-positive deletions ranging from 350bp-1kb that were characterized by many samples being genotyped with low GQ. Therefore, our motivation for partitioning the gnomAD-SV callset into a high-confidence subset was twofold: first, for ease of use and clarity when distributed to the broader community, and second, for the formal analyses conducted in this study. To this end, we labeled all variants with a final filter status as “PASS” in the VCF unless they met any of the following criteria:

1. The variant was unresolved; or
2. The variant had >15% of genotypes from PCR- samples masked during minimum GQ filtering as described above; or
3. The variant was a deletion between 300bp and 1kb in size and had >5% of genotypes from PCR- samples masked during minimum GQ filtering as described above; or
4. The variant had a VCF filter status including any of the following terms: “PCRPLUS_ENRICHED,” “UNSTABLE_AF_PCRMINUS,” or “MULTIALLELIC”.

All analyses presented in this study were restricted to SVs with PASS or MULTIALLELIC filter statuses, unless otherwise specified.

*Variant quality score recalibration*

Following all *post hoc* genotype- and site-level adjustments described above, we recalibrated variant quality scores (*i.e.*, “QUAL” values in the VCF) to reflect the median GQ among all samples with non-reference genotypes for each variant, irrespective of FILTER status. For this purpose, samples with homozygous non-reference genotypes were treated as if they had a GQ of 999 to reflect the high probability of at least one non-reference allele (even if the exact genotype is correct). For MCNVs, we treated individuals with copy states of one or three (*i.e.*, one copy different from diploid) as being heterozygous, and treated individuals with copy states of zero or at least four (*i.e.*, at least two copies different from diploid) as homozygous during QUAL score recalibration.

*Population assignments*

We next assigned samples to one of four populations based on genetic similarity inferred from SV genotypes. We first restricted to autosomal SVs with a global AF ≥1%, a VCF filter status of “PASS,” a variant quality (QUAL) score ≥100, lacking VCF INFO tags of “PCRPLUS_DEPLETED,” “UNSTABLE_AF_PCRPLUS,” and “VARIABLE_ACROSS_BATCHES,” and non-null genotypes for ≥99% of samples. We subsequently pruned SVs on a maximum linkage disequilibrium value of $R^2$ ≤0.2 over a rolling 1Mb window with BCFTools v1.9,[30] and filled missing genotypes with the mean allele count per site. We performed a principal component (PC) analysis of allele dosage per sample for these filtered variants. We assigned samples to one of four population labels (African [AFR]; East Asian [ASN]; European [EUR]; and Latino [LAT]) based on the top four PCs as labeled by a support vector machine (SVM) with a Gaussian kernel and 10-fold cross-validation using the e1071 package in R. We trained this SVM classifier on known population labels inferred from SNV data for a subset of samples

(N=7,575) as part of the gnomAD SNV & indel analysis,[6] and assigned each sample to a population if the SVM-estimated probability of membership for that sample was at least 0.8. Samples with membership probability below 0.8 for every population were assigned to an "other" [OTH] population category for purposes of analysis.

*Relatedness inference*

To infer genetic relatedness between samples, we filtered all SV to autosomal SVs with a global AF ≥0.015% (*i.e.*, observed in ≥2/14,000 samples), a VCF filter status of "PASS," a variant quality (QUAL) score ≥100, lacking VCF INFO tags of "PCRPLUS_DEPLETED," "UNSTABLE_AF_PCRPLUS," and "VARIABLE_ACROSS_BATCHES," and non-null genotypes for ≥98% of samples. Following preprocessing with PLINK v2.00a2LM,[31] we calculated kinship coefficients and identity-by-descent (IBD) fractions between all pairs of individuals with KING v2.2.2.[32] We filtered all 101 million nonredundant sample-sample pairs to the most likely related pairs of samples based on KING metrics, including HetConc > 0.2, IBS0 < 0.006, and Kinship > 0.1. From this subset, we trained an SVM on the KING results using ground truth family relationships available for a subset of samples with at least one known parent-child or sibling relationship to another sample in the cohort combined with 10,000 randomly selected pairs of samples where neither sample was known to have a first-degree relative in the cohort. This classifier was able to perfectly discriminate between ground truth pairs of first-degree relatives from presumably unrelated sample pairs (*i.e.*, zero sample pairs were misclassified), and also was able to distinguish between parent-child and sibling-sibling relationships with a near-perfect sensitivity and false positive rate: only 1/2362 ground truth parent-child relationship was misclassified as a sibling-sibling relationship, and reciprocally only 1/494 ground truth sibling-sibling relationships were misclassified as a parent-child relationship. We applied this SVM classifier to the KING metrics for all sample pairs passing our minimum KING thresholds to learn parent-child and sibling relationships. One sample from each pair of samples involved in relationships corresponding to predicted parent-child or sibling relationships were pruned from the dataset; we optimized this selection process to exclude the fewest possible samples such that all inferred sample pair relationships had at least one member excluded. Finally, we supplemented this agnostic, data-driven classification scheme with a list of children from known parent-child trios present in the dataset to account for rare situations where the above relatedness inference process did not optimally prioritize the exclusion of children over their parents. A breakdown of samples excluded at this step is provided in **Supplementary Table 2**.

*Final variant modifications and callset curation*

As the final step of callset curation, we performed multiple tiers of manual review. In some cases, this resulted in altered metadata and/or reclassifications for certain variants. These changes are summarized below:

- We assessed RD genotyping evidence for all autosomal CNVs ≥500kb, finding that 3/697 (0.4%) variants did not feature strong visual support for the expected copy number alterations. These three SVs were assigned the UNRESOLVED FILTER status in the final VCF.
- We assessed RD genotyping evidence for all autosomal CNV intervals ≥50kb involved in complex rearrangements, finding that 1/326 (0.3%) CNV intervals (corresponding to 1/249 [0.4%] unique complex SVs) did not exhibit strong visual support for the expected copy number alterations. This one complex SV lacking clear RD support was assigned the UNRESOLVED FILTER status in the final VCF.
- We reviewed all 120 SVs predicted to cause pLoF, IED, or CG of ≥10 different genes, and found that none were annotated incorrectly upon manual scrutiny of the affected genes and their positions relative to each SV.
- We identified 362 SVs with opposing predicted effects on the same gene, such as pLoF and partial gene duplication. All of these SVs were duplication-associated complex SVs, where computational prediction of the genic consequence is more challenging given the possibility of preserving a fully intact endogenous copy of the genes within the SV's associated duplications.[33] We manually reviewed the predicted rearrangement intervals for all 362 variants, and modified as appropriate for 96/362 (26.5%) of variants, while not modifying any predicted effects for genes without multiple conflicting annotations.
- We identified 2,873 SVs that had ≥50% coverage by regions of somatic hypermutability, such as T-cell receptor genes. These variants were excluded outright from the final VCF.
- We identified 18 SVs that had ≥20% coverage by N-masked regions of the reference genome. These variants were assigned the UNRESOLVED FILTER status in the final VCF.
- We identified 23 pairs of SVs that had identical coordinates and SV classes but were not merged due to sharing less than 50% of sample genotypes. We merged these variants in the final VCF, retaining the nonredundant union of non-reference sample genotypes in the merged record.
- We identified 740 RD-only canonical CNVs with ≥80% coverage by CNVs of the same class involved in a complex SV with ≥50% shared sample genotypes. Most (77.6%; 76/98) of the complex SVs corresponding to these overlapping CNVs were paired-CNV-flanked inversions, and in general these complex SVs were also twenty-fold larger (median = 43.9kb) than the average complex SV in the rest of the callset (median = 2.1kb). Therefore, we concluded that these canonical CNVs were likely redundant variants that did not meet our prior CNV redundancy consolidation criteria (50% reciprocal overlap with any one complex CNV interval, rather than all CNV intervals within a complex SV), and were thus excluded from the final VCF. The complex SVs overlapping these CNVs were retained, and the nonredundant union of non-reference samples was retained between each complex SV and its corresponding RD-only canonical CNV(s).

- We identified 5,162 SVs with ≥80% reciprocal overlap, ±3kb breakpoint proximity, and ≥50% shared sample genotypes with at least one different SV of the same class. We retained a single modified SV for each cluster of overlapping SVs. This modified SV was based on the median coordinates and QUAL score, and the nonredundant union of all non-reference sample genotypes, FILTER assignments, and other variant metadata. We preferentially retained non-pass FILTER statuses and homozygous alternate genotypes for each sample if multiple records had discordant FILTER or genotype metadata.
- We found examples where rare (AF<1%), RD-only CNVs were being apparently fragmented into multiple smaller, consecutive CNV calls. To correct these sites, we first identified all rare, RD-only CNVs that had ≥50% sample overlap with a different rare, RD-only CNV, and that these CNVs did not feature more than[11] 30% overlap by size and were not separated by more than the length of at least one of the CNVs (*e.g.*, a 100kb CNV must be within at least ±100kb from a non-overlapping smaller CNV). We next processed each cluster of qualifying CNVs as follows: (1) we assigned the largest original CNV as the "index" CNV; (2) we computed the total fraction of bases affected by CNVs in the cluster per sample genotyped as non-reference for any CNV in the cluster; (3) all samples with at least one third of the total CNV bases in the cluster were assigned to the index CNV, and these samples were removed from all other CNVs in the cluster; (4) all samples with less than one third of the total CNV bases in the cluster were removed from the index CNV and assigned back to their original contributing CNVs; (5) all index CNVs (N=1,289) had their coordinates modified to reflect the minimum and maximum coordinates among all CNVs in the cluster, and had predicted genic effects reannotated with svtk annotate; (6) all non-index CNVs in the cluster with at least one remaining non-reference sample (N=1,049) were retained with no additional modifications; and (7) all non-index CNVs in the cluster with no non-reference sample genotypes remaining (N=1,764) were removed outright from the final callset.
- We performed a targeted assessment of fragmented CNVs at known loci of recurrent, large CNVs.[11] This analysis followed the same procedure as above, with the following exceptions: we considered all CNVs, not just rare CNVs, to account for large blocks of segmental duplications that frequently flank these loci, and we required ≥33% sample overlap between pairs of putatively fragmented CNVs. This process identified an additional 139 CNVs requiring modification, including 39 extended index CNVs, 71 secondary CNVs with at least one remaining non-reference sample, and 29 CNVs to be excluded from the final callset due to having no remaining non-reference genotypes.
- We identified 917 insertions within one read length (±150bp) of a CNV breakpoint where: (i) the two SVs shared ≥90% non-reference sample overlap, and (ii) both SVs were detected by a PE/SR algorithm (DELLY or Manta). Given the close proximity of breakpoints and the strong non-reference genotype concordance, we concluded that these insertions were either misclassified by their original PE/SR algorithms, or alternatively might represent "scarring" at CNV breakpoints. Given this

uncertainty in variant classification despite the strong genotype correlations, we did not remove these insertion calls outright from the final VCF, instead marking them with the UNRESOLVED FILTER status.

We manually resolved 8 reciprocal translocations and 2 complex interchromosomal rearrangements that were initially incompletely resolved by the automated variant resolution step in module 05.

**Callset benchmarking**

To benchmark the technical properties and overall quality of the final gnomAD-SV map, we applied seven distinct analyses, as described below.

*Assessment of Mendelian violation rate from parent-child trios*

We counted the number of autosomal SV genotype combinations inconsistent with Mendelian segregation in 970 parent-child trios with PCR- WGS in this study. As we expected all inherited SVs to follow Mendelian segregation, and also expected less than one true *de novo* SV per generation (see **Figure 3a**),[1,7] we reasoned that nearly all Mendelian violations represent a combination of false-positive and false-negative genotypes in the child and/or parents. Per trio, we first isolated all biallelic, autosomal SVs where the child and both parents had non-null genotypes and at least one member of the trio had a non-reference genotype, then computed the fraction of those SVs that qualified as a Mendelian violation. We considered the following three possible cases of Mendelian violations:

- Apparent *de novo*: SVs where the child is heterozygous and neither parent carries any non-reference alleles.
- Spontaneous heterozygote: SVs where the child is homozygous for the alternate allele, and at least one parent is homozygous for the reference allele (*i.e.*, it should be impossible for the child to inherit two copies of the SV, without invoking extremely rare phenomena like uniparental disomy).
- Untransmitted homozygote: SVs where the child is homozygous for the reference allele, and at least one parent is homozygous for the alternate allele (*i.e.*, it should be impossible for the child to *not* have inherited at least one copy of the alternate allele).

For CNVs that were labeled as apparent *de novo*, we performed a secondary analysis of the child's CNV based on coverage by CNV calls in either parent to account for infrequent circumstances where parents and/or their children were assigned to different overlapping CNVs calls that likely represent the same genomic variant. If a CNV had ≥50% coverage by a CNV of matching type (*i.e.*, DEL or DUP) in at least one parent, that variant was no longer considered as an apparent *de novo* SV. To control for compound heterozygosity of large and small CNVs that might confound this analysis, we restricted parent CNVs to <10kb for proband CNVs <1kb, but considered all sizes of parent CNVs for proband CNVs ≥1kb.

For each trio, we computed the Mendelian violation rate to be the fraction of all qualifying SVs that met the above three criteria. We calculated the median across all 966 trios as the overall Mendelian violation rate for gnomAD-SV, and used these data to calculate other trio-based measurements of genotyping error rates as indicated in **Supplementary Table 4**.

*Comparison to chromosomal microarray data on matched samples*

We assessed the sensitivity of the gnomAD-SV discovery pipeline for large CNVs by comparing SVs from 1,893 samples in this study to existing CNV calls for those same samples from chromosomal microarray analysis in an earlier study.[13] Among these 1,893 samples was a subset of ASD-affected samples; as described above, these individuals were included for callset benchmarking analyses such as microarray comparisons, but were subsequently excluded prior to all other analyses presented in this study. We first converted the coordinates of CNV from microarray to GRCh37 using UCSC liftOver,[34] filtered to autosomal CNVs ≥ 40kb, and restricted to high-confidence calls by requiring pCNV $< 10^{-9}$ per recommendation of the authors. Next, we computed the number of PCR- samples in this study expected to carry each microarray CNV, and the fraction of the expected samples that also had at least 50% of the CNV covered by either a canonical or complex CNV from the WGS analyses in this study. Coverage was computed using BEDTools.[26] We considered each microarray CNV as captured in gnomAD-SV if the fraction of expected samples that had a matching WGS SV was at least 50%, and evaluated our sensitivity at two thresholds: one while considering all autosomal CNVs, and a second, more conservative threshold where we also excluded all microarray CNVs with ≥ 30% coverage by segmental duplications and/or simple repeats. We calculated overall sensitivity as the total number of microarray CNVs captured in this WGS analysis divided by the total number of eligible microarray CNV calls (**Supplementary Table 4**).

*Analysis of Hardy-Weinberg equilibrium across populations*

We evaluated the genotype distributions per SV under the null expectations set by the Hardy-Weinberg equilibrium (HWE; $1 = p^2 + 2pq + q^2$). While there are many biological reasons why some variants might violate HWE, such as recessive selection, mutational recurrence, or population stratification, the rate at which sites violate HWE can be used as a rough proxy of genotyping accuracy. Thus, we tabulated genotype distributions per population for each biallelic, autosomal SV, and computed a HWE P-value using the "HardyWeinberg" package in R.[5] We considered an SV to be in violation of HWE if its P-value was less than 0.05 following Bonferroni correction for the number of SVs tested per population.

*Comparison to long-read WGS data on matched samples*

Four samples in this study had also previously undergone PacBio long-read WGS as part of separate studies.[9,14,19] We used these data to assess the PPV of the gnomAD-SV discovery pipeline with an orthogonal long-read sequencing technology. SVs from this study for three of the four samples with long-read WGS were converted to hg38 coordinates via UCSC liftOver,[34] a necessary step to match the hg38-based alignment of the available PacBio data. We assessed support for each SV individually from the raw PacBio with VaPoR, a software package designed to autonomously validate SV calls *in silico* by performing comparative local realignments of long-read WGS reads.[35] Given that the performance of VaPoR is known to be sensitive to breakpoint coordinate precision (*e.g.*, see Figure 4 from the original VaPoR publication),[35] we restricted to biallelic, autosomal, FILTER "PASS" SVs from gnomAD-SV that were supported by SR evidence and lacked breakpoint overlap with annotated simple repeats, segmental duplications, or sites of somatic hypermutability. We also restricted the SVs considered here to the SV classes able to be evaluated by VaPoR, which included canonical, biallelic CNVs (*i.e.*, deletions & duplications), insertions, and inversions. Given the modest dependency of VaPoR validation rate on PacBio sequencing depth, we computed a study-level estimate of PPV from long-read WGS by averaging the PPV from each of the four samples analyzed here, and weighted each sample's PPV by the square root of their average PacBio sequencing depth.

We also estimated the accuracy of SV breakpoints reported in gnomAD-SV based on available long-read WGS data. To accomplish this, we compared the gnomAD-SV calls validated by VaPoR (above) to publicly available long-read WGS-derived SV callsets generated using matched methods for two of the four samples used in the VaPoR analyses.[8,9] We converted gnomAD-SV coordinates to hg38 where necessary using UCSC liftOver,[34] and then identified matching SVs between gnomAD and these external callsets by requiring ±1kb breakpoint proximity for both deletions and insertions, and imposing a further 50% reciprocal overlap requirement for deletions. For each SV from gnomAD-SV where a candidate match was found in the corresponding long-read WGS callset, we next computed the difference in reported coordinates for the left/lower and right/higher breakpoint respectively. We pooled these estimates of breakpoint accuracy across both samples and reported overall study-wide breakpoint accuracy estimates based on the pooled dataset.

*Comparison to SVs from the 1000 Genomes Project*

We obtained the 1000 Genomes phase 3 SV VCF as described by its original publication,[7] and converted it from VCF to BED format using *svtk vcf2bed*.[1] We performed minimal additional curation of this dataset: we left all information as provided by the 1000 Genomes Project, except for summing the frequency of all alternate alleles at sites where multiple alleles were listed (*e.g.* MCNVs). We then compared the gnomAD-SV callset to the 1000 Genomes Project callset while requiring at least 50%

reciprocal overlap by size and/or both breakpoints within ±300bp using BEDTools intersect. We further evaluated this comparison at two different levels of stringency: "strict" criteria, which required SV classes to match between candidate overlapping variants, and "loose" criteria, which did not apply this same requirement. We reported overlaps in this study using the "loose" criteria, but provide results from both criteria in **Supplementary Figure 8**. Furthermore, for each SV from the 1000 Genomes Project, we noted the AF of the overlapping SV in gnomAD-SV, if any. Where multiple candidate SVs in gnomAD-SV matched one call from the 1000 Genomes Project, we retained the AF most similar to the 1000 Genomes Project reported AF. We also performed these comparisons on a population-specific basis for the four populations matching between gnomAD-SV and the 1000 Genomes Project (AFR, AMR, EAS, EUR).

*Cross-population genotype analysis for doubleton SVs*

Fundamental population genetic principles dictate that most rare variants should be private to a single global population or subpopulation.[36] Taken to the most extreme case, variants that appear as two heterozygous genotypes in the population (i.e., "doubleton" variants) should disproportionately appear within, rather than across, populations. Despite the many factors that might cause deviations from this expectation, such as recurrent mutation and admixture, we nevertheless assessed the cross-population concordance for doubleton SVs in gnomAD-SV. For this analysis, we remove individuals with nonspecific population assignments (*i.e.*, "OTH"), and restricted to autosomal, resolved SVs appearing as heterozygous genotypes in exactly two unrelated individuals. We further restricted to SVs with ≤10% coverage by segmental duplications or simple repeats, as these genomic contexts are known to drive recurrent SV formation.[37] We computed the intra-population concordance rate for doubleton SVs by counting the number of SVs passing the filters above where both observations occurred in the same population, and divided that count by the total number of all SVs passing the filters above.

*Analysis of linkage disequilibrium between SNVs/indels and SVs*

We explored patterns of genotype correlation, or linkage disequilibrium (LD), between SVs discovered in this study and SNVs/indels discovered in a matched set of samples from a sister study.[6] As LD patterns are variable across populations, we restricted these analyses to the two populations (AFR and EUR) with at least 1,000 QC-pass samples overlapping between studies. Given the finer population substructure available for the SNV/indel callset (see Karczewski *et al.*),[6] we restricted samples for SNVs/indels specifically to non-Finnish Europeans (NFE), and matched those to EUR samples in this study. These filters retained 3,470 AFR samples and 1,883 EUR samples for LD analyses. We next considered all SNVs/indels and SVs that were autosomal, biallelic, had at least 1,000 samples with non-null genotypes and appeared at AF≥0.5% per population in both datasets, and calculated genotype correlations between all qualifying pairs of SNVs/indels and SVs within ±1Mb. We discarded SVs appearing in regions of low SNV/indel density (<2,000 SNVs/indels per 1Mb). After calculating LD between SNVs/indels and SVs, we subsequently restricted analyses to SVs with AF≥1% to avoid situations where slight discrepancies between SNV/indel and SV AFs might underrepresent the degree of LD for variants very near the AF~0.5% cutoff.

**Calculating adjusted proportion of singletons (APS)**

Given the strong dependence of SV size, class, genomic context, and WGS evidence on variant AFs and the proportion of singleton SVs (**Figure 1h** and **Extended Data Figure 5**), we aimed to develop a harmonized metric for comparing the proportion of singleton SVs across various subsets, annotations, or features. To accomplish this, we first stratified all autosomal SVs by mutational class. We further partitioned deletions & duplications based on whether an RD algorithm contributed to the CNV call or not, and partitioned insertions based on whether or not they were annotated as mobile element insertions. Third, we partitioned deletions, duplications, and insertions based on whether or not each variant had 5% coverage by annotated segmental duplications and simple repeats. We did not partition inversions or complex SVs given the relative sparsity of those variant classes. Thus, after including inversions and complex SVs as two separate categories, this process resulted in a total of 14 independent variant partitions. Next, as a proxy for near-neutral variation, we restricted to SVs explicitly intergenic SVs, or those with a predicted consequence of whole-gene inversion but no other effects, including promoter/UTR disruption or intronic localization. We further subdivided each subset of filtered SVs into 50 uniform bins ranked by SV size and computed the mean proportion of singletons and mean SV size within each bin. For each binned SV subset, we fit a nonlinear least-squares regression to predict the probability of being a singleton as a function of binned SV size by calculating the 11-bin rolling weighted mean of proportion of singletons for each bin. Finally, we applied these estimated singleton probabilities to all SVs in the gnomAD-SV callset, irrespective of coding effects or gene annotations. For any given subset of SVs, we defined the adjusted proportion of singletons (APS) as the observed proportion of singletons minus the mean of the singleton probabilities for all SVs in that same subset. We restricted all analyses using APS to autosomal biallelic SVs unless otherwise stated.

**Chromosome-level analyses of SV density**

To compute the density of SVs per chromosome, we first segmented all 22 autosomes into sequential 100kb windows, and excluded windows that overlapped centromeres. For each window, we tallied the number of SV per class that had any overlap with the window. For insertions, we only considered the insertion site in this analysis. This returned a matrix of SV counts per 100kb bin for all autosomes and SV classes. We computed the 11-window rolling mean per chromosome per SV class, yielding values per bin smoothed versus the surrounding 1Mb. Finally, we assigned each window to a percentile based on the position of that window on its respective chromosome arm relative to the chromosome's centromere where a value of -1 corresponded to the p-arm telomere, a value of 0 corresponded to the centromere, and a value of 1 corresponded to the q-arm telomere. To compute "meta-chromosome" averages, we segmented the range of normalized window positions (*i.e.*, -1 to 1) into 500 uniform bins, and averaged all windows across all chromosomes based on their chromosome-normalized window positions. We considered normalized positions within the outermost 5% of each chromosome arm to be "telomeric", the middle 90% of each arm to be "interstitial", and the innermost 5% to be "centromeric" for purposes of comparing chromosome contexts.

**Annotated repetitive element correlations**

We assessed correlations between SV density and seven different classes of annotated repetitive elements. Using the same filtered set of 100kb bins as generated for the chromosome-level analyses of SV density (see above), we annotated each 100kb bin for coverage versus segmental duplications, LINE repeats, SINE repeats, long terminal repeats (LTR), satellite repeats, and simple repeats. We downloaded all repeat annotations from the UCSC Genome Browser in native hg19 coordinates, and calculated the fraction of each 100kb bin covered per repeat class using BEDTools coverage.[26,34] We subsequently transformed each per-bin repeat coverage value into a Z-score within each repeat class across all bins. Next, for each SV class, we fit a generalized linear regression model to predict SV density based on the per-bin coverage for each of the seven annotated repeat classes considered here. We assessed the P-values of the fitted regression coefficients for each repeat class at a Bonferroni-corrected significant threshold across all SV classes (n=6) and repeat classes (n=7) considered here; thus, P-values were corrected for 42 total independent tests.

**Estimating SV mutation rates**

We estimated the SV mutation rate using the Watterson estimator with an effective population size ($N_e$) of 10,000, consistent with precedent set by prior SV studies.[7,38,39] Specifically, for each of the five major populations catalogued here, we first computed the Watterson estimator (ϴ) for each SV class as follows:

$$\hat{\theta}_w = \frac{K}{\sum_{i=1}^{n} \frac{i}{1}}$$

Where *K* was the number of SV sites observed per population for a given SV class and *n* was the total number of chromosomes analyzed in each population. We then solved for mutation rate (μ) as follows:

$$\mu = \frac{\hat{\theta}_w}{4N_e}$$

Finally, since the Watterson estimator is sensitive to differences in $N_e$, and the appropriate value of is known to be strongly influenced by population demographic history,[40] we computed the mean mutation rate across all five populations using the same estimate of to arrive at our global mutation rate estimate. We also computed a 95% confidence interval about the mean according to a *t* distribution.

**Comparisons of SVs to metrics of genic constraint against point mutations**

We constructed a simple statistical model to predict the number of rare SVs observed per gene for four different classes of gene-interacting SVs (pLoF, CG, IED, and whole-gene inversions). This approach is also described in the gnomAD SNV/indel study,[6] but is reproduced here for clarity. For each SV class, we first tallied the number of rare SVs per gene for all autosomal protein-coding genes. To prevent genes under strong selection from biasing the fit of this model, we restricted to genes in the $5^{th}$-$9^{th}$ deciles of observed:expected ratios for rare pLoF SNVs as described in the gnomAD SNV/indel analyses.[6] We fit a negative binomial regression model to predict the number of rare SVs per gene while including the following covariates: gene length, number of exons, median exon size, total number of nonredundant nucleotides in protein-coding exons, number of introns, median intron size, total number of nonredundant nucleotides in introns, and annotated overlap with segmental duplications. We applied this model to all protein-coding autosomal genes, which yielded expected counts of rare SVs per gene for each functional class. For comparisons to constraint against missense SNVs, we re-trained these models based on genes in the 5th-9th deciles of observed:expected ratios for rare missense SNVs.

We compared SVs in this study to constraint against damaging point mutations for both pLoF and missense SNVs. For each comparison, we first ordered all autosomal protein-coding genes based on the observed:expected measurement of SNV constraint, and subsequently grouped genes into 100 bins based on SNV constraint percentile. Next, for each bin of genes, we summed the total number of rare SVs observed in gnomAD-SV for all genes in the bin and divided this total by the expected number of rare SVs based on the regression model (described above). This calculation produced an observed:expected ratio of rare SVs for each percentile of SNV constraint scores. We assessed the correspondence between SV and SNV constraint for all 100 bins of genes using a Spearman's rank correlation test. Finally, we repeated this entire analysis while restricting to canonical SVs with precise breakpoints (*i.e.*, SVs with split-read support) to confirm that the inclusion of balanced and complex SVs and/or inaccurate coding annotations weren't unduly influencing these conclusions. We found that restricting to precise, canonical SVs had effectively no impact on the correlations between rare SVs and SNV constraint metrics, nor the conclusions drawn from those data.

**Estimating noncoding selection against *cis*-regulatory annotation classes**

Independent of our gene-based annotations and analyses, we also conducted a series of analyses examining evidence for selection against noncoding SVs across a variety of *cis*-regulatory annotation classes. The relevant data and analyses are described below.

*Definition of noncoding CNVs*

We restricted all SVs in this analysis to canonical biallelic deletions or duplications that did not overlap any protein-coding exons. We also restricted all CNVs and elements in this analysis to relatively unique genomic regions, defined as any CNV or element with <30% coverage by segmental duplications, simple repeats, N-masked unalignable regions of the reference genome, and known somatically hypermutable regions.

*Curation of functional annotation classes*

We curated a set of 14 functional annotation classes to be considered in this analysis. For all annotation classes, we restricted to relatively unique autosomal regions as was performed for CNVs (see above). Additional curation steps for each functional annotation class are described below:

- *Topologically associated domain (TAD) boundaries*: we downloaded a list of TADs as defined by Hi-C in a human fetal fibroblast cell line (IMR90) from GEO accession GSE63525.[41] TAD boundaries were defined as the 10kb intervals (±5kb) centered on the start and end coordinates of each TAD. Overlapping TAD boundaries were collapsed with BEDTools merge.[26]
- *Chromatin loop anchors*: we downloaded a list of chromatin loops as defined by Hi-C in IMR90 cells from GEO accession GSE63525.[41] Chromatin loop anchors were defined as the 10kb intervals (±5kb) centered on the start and end coordinates of each loop. Overlapping loop anchors were collapsed with BEDTools merge.[26]
- *DNAse1 hypersensitive sites (DHS)*: we downloaded the consensus set of clustered DHS peaks derived from 125 cell types by the ENCODE project (V3) from the UCSC Genome Browser in hg19 coordinates.[34,42] We required all DHS clusters to have a score >500 and to have been discovered in ≥50% of cell types (>62/125). Overlapping DHS clusters passing these criteria were collapsed using BEDTools merge.[26]
- *Transcription factor (TF) binding sites (TFBS)*: we downloaded the consensus set of clustered TFBS data derived for 161 transcription factors in 91 cell types by the ENCODE project (V3) from the UCSC Genome Browser in hg19 coordinates.[34,42] We required all TFBS to have a score >200 and to have been discovered in ≥5% of cell types (>4/91), which restricted this dataset to ~23% (37/161) of all TFs assayed. Overlapping TFBS meeting these criteria were collapsed with BEDTools merge.[26]

- *Experimentally validated enhancers*: we downloaded a list of enhancers with experimentally confirmed *in vivo* activity provided by the Vista database.[43] We restricted to enhancers labeled as having "reproducible expression in the same [physiological] structure in at least three independent transgenic [mouse] embryos." (https://enhancer.lbl.gov/aboutproject_n.html).
- *Computationally predicted enhancers*: we downloaded a list of enhancers in 29 primary tissues, primary cells, or primary cell culture from the EnhancerAtlas database in hg19 coordinates.[44] We restricted to enhancers observed in ≥20% of tissues (≥6/29) based on 50% reciprocal overlap by size as calculated with BEDTools intersect.[26] Overlapping enhancers passing these criteria were collapsed using BEDTools merge.[26]
- *Super enhancers*: we downloaded all super enhancers predicted across 99 tissues from dbSUPER in hg19 coordinates,[45] and retained all super enhancers observed in ≥5% (5/99) tissues based on 50% reciprocal overlap by size as calculated by BEDTools intersect. Overlapping super enhancers were collapsed with BEDTools merge.[26]
- *Chromatin states*: we downloaded 15-state ChromHMM annotations in 200bp windows for 129 tissues as provided by the Roadmap Epigenomics Project.[46] For four chromatin states (genic enhancers, enhancers, bivalent enhancers, and polycomb repressed elements), we selected all 200bp windows where at least one-third of tissues (≥43/129) matched that state, and subsequently merged these 200bp windows with BEDTools merge while allowing for up to ±1kb to separate adjacent windows prior to merging.[26]
- *Ultraconserved noncoding elements (UCNEs)*: we downloaded UCNEs in hg19 coordinates from UCNEBase on January 8, 2019.[47] Overlapping UCNEs were collapsed with BEDTools merge.[26]
- *Human accelerated regions (HARs)*: we collected a list of HARs by taking the union of HARs reported by three previously published studies.[48-50] For each study, a list of HARs was lifted over to GRCh37 using the UCSC liftOver tool (where necessary).[34] Intervals across all three studies were collapsed into a nonredundant set using BEDTools merge.[26]
- *Recombination hotspots*: we downloaded a recombination frequency map at 10kb resolution averaged across males and females from deCODE Genetics.[51] We excluded bins with unsequenced bases, then defined any 10kb bin in the top 10% of all remaining recombination frequency scores as a recombination hotspot, and merged adjacent hotspots with BEDTools merge.[26] Finally, we lifted over all recombination hotspots from hg18 reference assembly coordinates to GRCh37 with the UCSC liftOver tool while requiring at least 50% of the original locus to map to GRCh37.[34]

*Calculation of APS for noncoding CNVs overlapping functional annotation classes*

For each of the 14 functional annotation classes defined above, we calculated APS for all noncoding deletions or duplications with any overlap with at least one element from that annotation class. APS for deletions and duplications was calculated separately. We also calculated APS for deletions or duplications that completely covered at least one element from that annotation class. Finally, we considered two additional annotation classes: one defined as "any annotation," which was the union of all 14 annotation classes considered here, and one defined as "no annotations," which was defined as the inverse intersection of deletions or duplications against the union of all 14 annotation classes. For each comparison, we assessed significant deviation from the expected APS value of zero for neutral variation using a two-sided one-sample t-test against an expected mean of zero. Finally, we adjusted P-values for the 32 tests performed here using a Bonferroni correction.

**Intersection of gnomAD-SV and published genome-wide association studies**

We performed a series of analyses to understand the role that SVs documented in this study might play in genome-wide association studies (GWAS) of common SNVs in large cohorts for a spectrum of human traits and diseases. These analyses were performed in a series of steps, as follows.

First, we combined two sources of publicly available GWAS results. Specifically, we collated GWAS results from the NHGRI-EBI catalog of published GWAS results ("GWAS Catalog") v1.0.2 and a recent GWAS analysis of 4,023 phenotypes in 361,194 European samples from the UK BioBank ("UKBB").[52-54] We lifted over the GWAS Catalog loci from hg38 to GRCh37 using UCSC liftOver prior to analysis.[34], and restricted UKBB GWAS results to only genome-wide significant loci with a P-value < $10^{-8}$.

Second, using the LD calculations between SVs and SNVs/indels for a subset of samples in gnomAD-SV as described earlier, we next compared SVs in strong LD ($R^2 \geq 0.8$) against the GWAS results curated above. As there is an established bias toward Europeans in current GWAS databases,[55] we restricted our comparison to SVs in strong LD with at least one SNV or indel specifically in European samples. SVs were considered to be overlapping a GWAS locus if the SV had at least one SNV in strong LD overlapping overlap the same nucleotide as the reported GWAS association.

Finally, we computed enrichments for the set of common SVs in strong LD with at least one GWAS association across all gene-centric functional classes with at least one count, including pLoF, partial CG, promoter UTR, and intronic SVs, and also computed similar enrichments for intergenic SVs as a comparison group. We used a two-sided Fisher's Exact test to compare the fraction of SVs for each functional class in strong LD with any SNV not reported as a significant GWAS association to the fraction of SVs in strong LD with an SV that was reported as a significant GWAS association. Enrichments were reported as odds ratios, 95% confidence intervals, and unadjusted P-values.

##### Analysis of genomic disorder loci

We compared the frequency of CNVs at putatively pathogenic genomic disorder (GD) loci in gnomAD-SV to their corresponding frequencies reported from CMA in a recent analysis of 396,725 participants from the UK BioBank (UKBB).[11] We restricted these analyses to the subset of 10,047 samples in gnomAD-SV with no indication of neuropsychiatric disease. We collected UKBB CNV frequency data from the original publication for the 54 GD loci considered in the UKBB analysis, which required at least five UKBB participants to be GD carriers. We further excluded a total of five GDs: deletions and duplications of 22q11.2 [distal], deletions and duplications of 15q11.2, and deletions of *NRXN1*. We excluded the 22q11.2 distal GD and the 15q11.2 due to their proximity to (or direct overlap with) antibody part genes, which were blacklisted during the creation of gnomAD-SV, and excluded *NRXN1* due to this locus being a known nonrecurrent GD region corresponding to a well-described single-gene deletion syndrome.[56] Per consultation with the authors of the UKBB analysis, we also imposed a restriction on the thrombocytopenia absent radius (TAR) GD locus by requiring CNVs in gnomAD-SV to span at least two of the three segmental duplication tracts within the region. After curation, we retained 49 GDs for analysis. We intersected the coordinates of these 49 GD loci against all biallelic canonical and complex CNVs in the gnomAD-SV dataset using BEDTools intersect requiring CNVs in gnomAD-SV to have 30% overlap of the GD locus, and computed the carrier frequency as the number of individuals with at least one non-reference SV allele. We determined 30% overlap to be the optimal parameter for this analysis by manual inspection of all GD loci, and to account for differences in breakpoint coordinates between WGS and CMA. Where one gnomAD SV matched multiple possible GD loci, we counted each gnomAD SV only once in total towards the GD that best matched the gnomAD SV based on reciprocal overlap. We compared carrier frequencies between UKBB and gnomAD-SV using Fisher's exact text, and significance was assessed after Bonferroni correction for multiple comparisons. We also computed odds ratios and 95% confidence intervals for these CNVs in developmental disorders (DDs) by counting the number of DD patients carrying CNVs matching these GD loci from a large, previously published DD cohort with CNV data from chromosomal microarray,[10] followed by a Fisher's Exact Test.

##### gnomAD-SV callset downsampling analyses

Several of our investigations in this dataset involved either projecting the properties of SV datasets hypothetically attainable from larger sample sizes, or estimating what fraction of the current SV callset would have been obtained had we sequenced fewer individuals. To accomplish this, we first performed a single, standardized set of iterative callset downsamplings, and used this series of downsampled callsets in all pertinent analyses. We randomly downsampled the gnomAD-SV VCF to contain 1, 2, 3, 4, 5, 6, 7, 8, 9, 10, 25, 50, 75, 100, 250, 500, 750, 1,000, 2,500 5,000, 7,500 or 10,000 samples, respectively, and generated five independent downsampled VCFs at each sample size. This combination of 22 sample sizes performed five times each yielded a total of 110 downsampled VCFs, each with a distinct subset of samples from the full gnomAD-SV callset. For each downsampled VCF, we retained variant information exactly as it appeared in the full callset, except for excluding all individuals not selected to be part of that downsampling and subsequently removing all sites that no longer had at least one non-reference allele in the downsampled VCF due to all non-reference allele carriers being excluded during downsampling. As with all per-sample analyses, we restricted this analysis to only include PCR- samples.

**Predicting the rate of clinically reportable incidental findings from SVs**

We estimated the rate of clinically reportable incidental findings in gnomAD-SV to derive a population-based estimate for SV analyses from WGS. We first restricted the gnomAD-SV callset to very rare (AF<0.1%), biallelic, autosomal SVs resulting in pLoF of one of 57 autosomal genes marked as clinically reportable for incidental findings per recommendations by the American College of Medical Genetics (ACMG)[57] and classified variants following the ACMG guidelines for the interpretation of sequence variants.[58] All heterozygous SVs that disrupted a gene associated with an autosomal recessive disorder and/or only have evidence for a gain-of-function (GoF) pathogenic mechanism were classified as benign. We classified SVs that disrupted genes that predominantly have a GoF pathogenic mechanism, but there exists some evidence for a LoF pathogenic mechanism, as a variant of unknown significance (VUS). All SVs that disrupted a gene known to be associated with disease through a LoF mechanism were classified as pathogenic or likely pathogenic, depending on the strength of evidence available from the existing literature. SVs were determined to be clinically reportable for the purposes of this study if they met criteria to be pathogenic or likely pathogenic.

**Evaluation of callset filtering on key results**

We examined the quality of filtering thresholds used in this study and assessed their impact on the results reported herein. We produced three VCFs, each corresponding to a certain set of filtering criteria. The first VCF, which represented a hypothetical callset with looser filtering, included all SVs irrespective of their FILTER status (*i.e.*, included variants with failing FILTER statuses, which we excluded for our analyses presented in this study). The second VCF was the callset exactly as presented in this study. The third VCF, which represented a hypothetical callset with stricter filtering, included only SVs with a PASS FILTER status, and additionally required all SVs to have QUAL > 500 (also see **Supplementary Figures 11-12** for the justification of this additional criterion). For each of these three VCFs, we repeated all analyses exactly as described in the main study and methods, with the single exception being that all BND variants were excluded from all analyses. The outcomes of these analyses are provided in **Supplementary Figure 24**.

##### Compliance with ethical regulations

We have complied with all relevant ethical regulations. This study was overseen by the Broad Institute's Office of Research Subject Protection and the Partners Human Research Committee, and was given a determination of Not Human Subjects Research. Informed consent was obtained from all participants.

##### Data availability

All gnomAD-SV site-frequency data for appropriately consented samples (N=10,847) have been distributed in VCF and BED format via the gnomAD Browser (https://gnomad.broadinstitute.org/downloads/), as well as from NCBI dbVar under accession nstd166. Furthermore, these SVs have been integrated directly into the gnomAD Browser.[59] The architecture of the gnomAD Browser is described in the main gnomAD study,[6] as well as instructions for how to access and query the data hosted therein. Refer to **Extended Data Figure 10** for a highlight of the SV-related features. All VCFs for the analyses of ASD families for disease association comparisons will be deposited in SFARIbase (https://base.sfari.org/) and are available to qualified researchers by applying online.

##### Code availability

The overall structure and availability of code used in this study is outlined on the home page of the main gnomAD-SV github repository (https://github.com/talkowski-lab/gnomad-sv-pipeline). The gnomAD-SV discovery pipeline is publicly available via a series of methods configured for the FireCloud/Terra platform (https://portal.firecloud.org/#methods) under the methods namespace "Talkowski-SV". The svtk software package used extensively in the gnomAD-SV discovery pipeline is publicly available via gitHub (https://github.com/talkowski-lab/svtk). Most custom scripts used in the production and/or analysis of the gnomAD-SV dataset are publicly available via gitHub (https://github.com/talkowski-lab/gnomad-sv-pipeline). All code is made available under the MIT License, unless stated otherwise.
